# CRISPR-Assisted DNA Detection, a novel dCas9-based DNA detection technique

**DOI:** 10.1101/2020.05.13.093062

**Authors:** Xinhui Xu, Tao Luo, Jinliang Gao, Na Lin, Weiwei Li, Xinyi Xia, Jinke Wang

**Affiliations:** State Key Laboratory of Bioelectronics, Southeast University, Nanjing 210096, China; Jinling Hospital, Nanjing University School of Medicine, Nanjing 210002, China

**Keywords:** CRISPR/Cas9, DNA, detection, HPV

## Abstract

Nucleic acid detection techniques are always critical to diagnosis, especially in the background of the present COVID-19 pandemic. The simple and rapid detection techniques with high sensitivity and specificity are always urgently needed. However, the current nucleic acid detection techniques are still limited the traditional amplification and hybridization. To overcome the limitation, we here develop a CRISPR/Cas9-assisted DNA detection (CADD). In this detection, DNA sample is incubated with a pair of capture sgRNAs (sgRNAa and sgRNAb) specific to a target DNA, dCas9, a signal readout-related probe, and an oligo-coated solid support beads or microplate at room temperature for 15 min. During this incubation, the dCas9-sgRNA-DNA complex is formed and captured on solid support by the capture sequence of sgRNAa and the signal readout-related probe is captured by the capture sequence of sgRNAb. Finally the detection result is reported by a fluorescent or colorimetric signal readout. This detection was verified by detecting DNA of bacteria, cancer cell and virus. Especially, by designing a set of sgRNAs specific to 15 high-risk human papillomaviruses (HPVs), the HPV infection in 64 clinical cervical samples were successfully detected by the method. All detections can be finished in 30 minutes at room temperature. This detection holds promise for rapid on-the-spot detection or point-of-care testing (POCT).

## 1. Introduction

Nucleic acid detection is essential to agriculture, food industry and medicine. Therefore, development of nucleic acid detection techniques always attracts intensive attentions in these fields, which promotes the sustainable technique development. In mechanism, three types of DNA detection techniques are widely used. The first is the DNA amplification mainly based on various polymerase chain reaction (PCR) including traditional PCR (tPCR)(1), quantitative PCR (qPCR)(2) and digital PCR (3). DNA amplification also includes various isothermal amplifications (4,5), such as rolling circle amplification (RCA), recombinase polymerase amplification (RPA), loop-mediated Isothermal amplification technology (LAMP), nucleic acid sequence-dependent amplification (NASBA), and nicking enzyme amplification reaction (NEAR). The second is nucleic acid hybridization including traditional Southern and Northern blotting, and DNA microarrays or chips. The third is DNA sequencing including traditional Sanger sequencing, next-generation sequencing (NGS) and nanopore sequencing. Each technique has its advantages and still plays important role in various nucleic acid tests. Nevertheless, in application to the increasingly demanded on spot test or point-of-care diagnosis (POCT), these techniques are now challenged by their intrinsic limitations. Especially, the time-consuming and instrument-dependent processes of enzymatic or hybridization reaction make these techniques difficult to be the simple, rapid and portable diagnostic tools. Moreover, these techniques are still challenged by the unsatisfied sensitivity or specificity resulted from enzyme-introduced mutations and mismatched hybridization. Fortunately, the advent of Clusters of regularly spaced short palindrome repeats (CRISPR) provides new chance to change the situation (6,7).

CRISPR was first discovered in bacteria genome as an immunity weapon to phages (8). Immediately its worth as a new type of gene editing tool was rapidly mined (9,10). Especially, Cas9 of type II CRISPR system was intensively explored due to its benefit requiring only one CRISPR-associated nuclease. This single CRISPR protein can function as sequence-specific DNA endonuclease by associating with CRISPR-related RNA (crRNA) and trans-activated crRNA (tracrRNA), in which tracrRNA can activate Cas9 nuclease and crRNA guide Cas9 to target DNA. TracrRNA and crRNA can be ligated into a single guide RNA (sgRNA)(11), greatly simplifying the application of CRISPR/Cas9 system. This system is then widely used to edit, regulate and target genomes (12). Due to the high specificity of interaction with its targets, Cas9 has also been gradually used to develop new DNA detection methods. For example, the CRISPR/Cas9 system has been early used to detect Zika viruses and can type Zika viruses (13). Recently, several DNA detection methods were developed by using its typical double-stranded DNA (dsDNA) cleavage activity of Cas9, such as CRISPR-typing PCR (ctPCR)(14-16), CUT-LAMP (17), FLASH(18), and. Several DNA detection methods were developed by using the Cas9 nickase activity, such as Cas9nAR (Cas9 nickase-based amplification reaction) (19), and CRISDA (CRISPR-Cas9-triggered nicking endonuclease-mediated Strand Displacement Amplification) (20). In addition, several DNA detection methods were developed by using the catalytically deactivated Cas9 (dCas9), such as CRISPR-Chip (21) and paired dCas9 proteins linked to split halves of luciferase (22). However, the potential of Cas9/sgRNA system in nucleic acid detection and typing has not yet been fully explored by the current studies.

In contrast to CRISPR/Cas9, other CRISPR/Cas proteins with collateral cleavage activity attracted increasing attention and have been intensively explored in recent years (23). Firstly, Cas13a (also called C2c2) of type III CRISPR system was used to develop SHERLOCK (Specific High-sensitivity Enzymatic Reporter UnLOCKing) by using its target-cutting activated collateral cleavage activity for single-stranded RNA (ssRNA) (24-27). Then Cas12a (also known as Cpf1) (28) and Cas14 (also known as Cas12f) (29) with collateral cleavage activity of single-stranded DNA (ssDNA) was used to develop DETECTR (DNA Endonuclease Targeted CRISPR Trans Reporter) (28,29) and HOLMES (a one-HOur Low-cost Multipurpose highly Efficient System) (30,31). Immediately, the HOLMESv2 (HOLMES2.0) and CDetection technology were developed by using the collateral cleavage activity to single-stranded DNA (ssDNA) of Cas12b (also known as C2c1) (32,33). These methods are demonstrated to have attomolar sensitivity and single-base specificity. The ultrahigh sensitivity of these methods partially rely on the signal amplification resulted from collateral cleavage activity of related Cas enzymes. These methods are therefore used to detect various pathogenic viruses such as Zika virus (ZIKV) (24,25), dengue virus (DENV) (24), human papillomavirus (HPV) (28), Japanese encephalitis virus (JEV)(32), African swine fever virus s (ASFV) (34), and *Mycobacterium tuberculosis* (MTB)(35). Recently, these techniques are striving to detect SARS-CoV-2 (36-41). However, these CRISPR Cas-based detection methods typically require DNA pre-amplification with polymerase chain reaction (PCR) (31), reverse transcription-RPA (RT-RPA)(25), recombinase polymerase amplification (RPA) (25,27,28,34,35), Loop-mediated isothermal amplification (LAMP) (32,39), reverse transcription-LAMP (RT-LAMP) (32,39), and asymmetric PCR (32). These pre-amplifications also partially contribute to the ultrahigh sensitivity of these methods. Additionally, SHERLOCK also needs reverse transcription and T7 in vitro transcription when detecting RNA (27). Such amplification dependence still makes these methods difficult to become simple, rapid and portable diagnostic tools.

In this study, we develop a new DNA detection method based on CRISPR/Cas9 named as CRISPR/Cas9-assisted DNA detection (CADD). In this method, a pair of capture sgRNAs (sgRNAa and sgRNAb) are designed for a target DNA. SgRNAa and sgRNAb harbor a different 3′ terminal capture sequence. When target DNA is bound by a pair of dCas9-sgRNA complexes, the dCas9-sgRNA-DNA complex will be captured on surface of beads or microplate via annealing between an oligonucleotide coupled on solid supports and capture sequence of sgRNAa. Then the captured dCas9-sgRNA-DNA complex is reported by a kind of signal reporter captured by the capture sequence of sgRNAb. This method was validated by detecting DNA of bacteria, cancer cell and virus. Especially, by designed a set of sgRNAs specific to 15 high-risk human papillomaviruses (HPVs), this study successfully detected 64 clinical cervical samples with this method. Of importance, a target DNA can be rapidly detected in less than 30 minutes using a signal readout of fluorescent hybridization chain reaction (HCR). This method has its unique advantages over the current methods such as free of enzyme, pre-amplification, modular heaters, simplicity and rapidness.

## 2. Materials and methods

### 2.1. Preparation of sgRNA

SgRNA was designed with online sgRNA design software Chop-Chop (http://chopchop.cbu.uib.no/), using hg19 as the reference genome. The designed sgRNAs are shown in Table S1. Each DNA target had a pair of sgRNA, named sgRNAa and sgRNAb, respectively. According to the designed sgRNA, primers (Table S2) were synthesized to amplify sgRNA template by PCR using our previous protocol (42). The PCR-amplified sgRNA template has a T7 promoter sequence. SgRNA was then prepared by in vitro transcription using sgRNA template as previously described (42). The prepared sgRNA had a 5′-end 20 bp target DNA-specific sequence and a 3′-end capture sequence. SgRNAa had a 3′-end capture sequence named as Capture 1 that was used to anneal with the capture oligonucleotide immobilized on the surface of the magnetic beads or microplate. SgRNAb had a 3′-end capture sequence named as Capture 2 that was used to anneal with a signal reporter-associated oligonucleotide.

### 2.2. Preparation of HPV plasmids

The full-length L1 fragments of 15 high-risk HPV (hrHPV) were cloned in the pMD plasmid (Takara) to prepare HPV plasmids, including: pMD-HPV16, pMD-HPV18, pMD-HPV31, pMD-HPV33, pMD-HPV35, pMD-HPV39, pMD-HPV45, pMD-HPV51, pMD-HPV52, pMD-HPV56, pMD-HPV58, pMD-HPV59, pMD-HPV66, pMD-HPV68, and pMD-HPV73.

### 2.3. Preparation of HCR hairpin

Oligonucleotides FAM-hairpin-1 and FAM-hairpin-2 (Table S3) were synthesized. Oligonucleotides were dissolved in TE buffer (10 mM Tris-HCl, pH 8.0, 1 mM EDTA) at a concentration of 100 μM. For preparing HCR hairpins, 30 μL of FAM-hairpin-1 and FAM-hairpin-2 were mixed. The mixture was incubated at 95 °C for 5 min and then naturally cooled to room temperature (RT). The HCR hairpins (Hairpin 1 and Hairpin 2) were kept at 4 °C. In order to evaluate HCR hairpins and sgRNA, liquid-phase HCR reactions were performed. Each reaction (20 μL) contained 1× Binding buffer (5 mM MgCl_2_, 0.3 M NaCl, 0.1% BSA, 10 mM Tris-HCl, pH 8.0, 1 mM EDTA), 0.2 μM Hairpin 1, 0.2 μM Hairpin 2, and 30 nM Initiator (Table S3) or sgRNA. The reaction was kept at RT for 20 min and detected by agarose gel electrophoresis.

### 2.4. Preparation of Beads@oligo

A biotin-modified oligo RE-flanking (Table S3) was synthesized and dissolved in TE buffer at a concentration of 100 μM. In order to prepare DNA-coupled Beads (Beads@oligo), 1 μL of the magnetic beads (Dynabeads™ M-280 Streptavidin; Invitrogen) (10 μg; 6–7 × 10^5^ beads) was first washed twice with 1× Binding buffer. The beads were then incubated with 2 μL of 10 μM RE-flanking oligo in rotation at RT for 20 min. The beads were washed 3 times with 1× Binding buffer. Finally the beads were resuspended in 1× Binding buffer and kept at 4 °C for later use.

### 2.5. Clinical HPV DNA sample

All procedures used in this research were performed according to the Declaration of Helsinki. This study was approved by the Ethics Committee of Jinling Hospital (Nanjing, China). All participants were recruited from the Jinling Hospital with informed consent. The clinical HPV detection was performed by Jinling Hospital (Nanjing, China) using a human papillomavirus genotyping (type 23) detection kit (PCR-reverse dot hybridization method) (Asia Energy Biotechnology, Shenzhen). The gDNA extraction and HPV detection (a PCR-reverse dot hybridization method) were all performed with this kit. DNA was first tested by hospital and then the left DNA was brought to our laboratory. Two batches of clinical DNA samples were detected by CADD.

### 2.6. Beads-HCR detection

The Beads-HCR detection reaction (20 µL) contained 1× Binding buffer, 15 nM sgRNAa, 15 nM sgRNAb, 30 nM dCas9 protein (dCas9 Nuclease, S. pyogenes, M0652, New England Biolabs), 1–4× 10^4^ Beads@oligo, 40 U RNase inhibitor (RNaseOUT ™ Recombinant Ribonuclease Inhibitor, Invitrogen), and various amount of DNA sample (see figures). The reaction was incubated at RT for 15 min in rotation. The beads were then washed 3 times with 1× Binding buffer. The beads were added 20 µL of HCR solution (0.2 μM FAM-hairpin-1, 0.2 μM FAM-hairpin-2 and 1× Binding buffer) and incubated at RT for 15 min in rotation. The beads were then dropped on slide glass and covered with a cover glass. The beads were imaged with a fluorescence microscope. The beads images were analyzed with Image pro.

### 2.7. Beads-ELISA detection

Single-target Beads-ELISA detection reaction (20 µL) contained 1× Binding buffer, 15 nM sgRNAa, 15 nM sgRNAb, 30 nM dCas9 protein, 1–4×10^4^ Beads@oligo, 0.5 µM oligo re-biotin (Table S3), 40 U RNA Enzyme inhibitor, and DNA sample (see figures for amount). Multi-target Beads-ELISA detection reaction (20 µL) contained 1× Binding buffer, 15 nM each of sgRNAa, 15 nM each of sgRNAb, 15× N nM dCas9 protein, 1–4×10^4^ Beads@oligo, 0.5 µM oligo re-biotin, 40 U RNase inhibitor, and various amount of DNA sample (see figures); where N is the number of targets detected. The reaction was incubated at RT for 15 min in rotation. The beads were then washed three times with a washing buffer (1× dCas9 buffer contained 10 U of RNase inhibitor and 0.5% BSA), in which 1× dCas9 buffer can be purchased from New England Biolabs as NEBuffer 3.1 or prepared at home. The beads were incubated with 20 µl washing buffer containing of 8 ng of HRP-conjugated Streptavidin (Sangon Biotech, Shanghai) for 3 min. The beads were washed three times with the washing buffer and finally re-suspended in 30 µl of washing buffer. The beads were transferred into microplate and added 50 µl of TMP chromogenic solution for ELISA (P0209-100ml; Beyotime). The beads were incubated at RT for 10 min. The microplate was read at 630 nm and imaged with a BioRad gel imager in staining-free mode.

### 2.8. Microplate-ELISA detection

An amino-modified oligo RE-NH_2_ (Table S3) was covalently coupled to the DNA-BIND 96-well microplate (Corning) according to the instructions of manufacturer to prepare the oligo-coated microplate. Single-target Microplate-ELISA detection reaction (80 µL) contained 1×Binding buffer, 15 nM sgRNAa, 15 nM sgRNAb, 30 nM dCas9 protein, 0.5 µM oligo re-biotin (Table S3), 40 U RNA Enzyme inhibitor, and various amount of DNA sample (see figures). Multi-target Microplate-ELISA detection reaction (80 µL) contained 1× Binding buffer, 15 nM each of sgRNAa, 15 nM each of sgRNAb, 15 nM × N dCas9 protein, 0.5 µM oligo re-biotin (Table S3), 40 U RNase inhibitor, and various amount of DNA sample (see figures); where N is the number of targets detected. The reaction was added to the oligo-coated microplate. The microplate was incubated at RT for 15 min on a horizontal mixer and then washed three times with the washing buffer. The microplate was added with 100 µL of washing buffer containing 8 ng of HRP-conjugated Streptavidin and incubated at RT for 3 min. The microplate was then washed three times with the washing buffer. The microplate was then added with 30 μL of washing buffer and 50 μL TMP chromogenic solution for ELISA (Beyotime). The microplate was incubated for 10 min. The microplate was read at 630 nm and imaged with a BioRad gel imager in staining-free mode.

## 3. Results

### 3.1 Beads-HCR CADD detection

To explore the feasibility of CADD, we first designed a Beads-HCR method (Fig. 1A), in which the fluorescent hybrid chain reaction (HCR) was used as signal readout. A pair of sgRNA were designed for a target DNA. Different from the traditional sgRNA, the sgRNA used by CADD was designed to have a short extended 3′ terminal capture sequence that can anneal with other functional oligonucleotides. In detection, a pair of dCas9-sgRNA (dCas9-sgRNAa and dCas9-sgRNAb) first binds the target DNA. The dCas9-sgRNA-DNA complex is then captured onto the surface of beads via annealing between the capture sequence on the sgRNAa and a complementary oligonucleotide coupled on beads. The beads are then washed on magnet and then two HCR Hairpins (Hairpin 1 and Hairpin 2) are added. The capture sequence of sgRNAb can anneal with Hairpin 1 to initiate HCR. Because the Hairpins are labeled by fluorescein, fluorescent signal can be produced on the surface of beads by HCR.

**Fig. 1.**
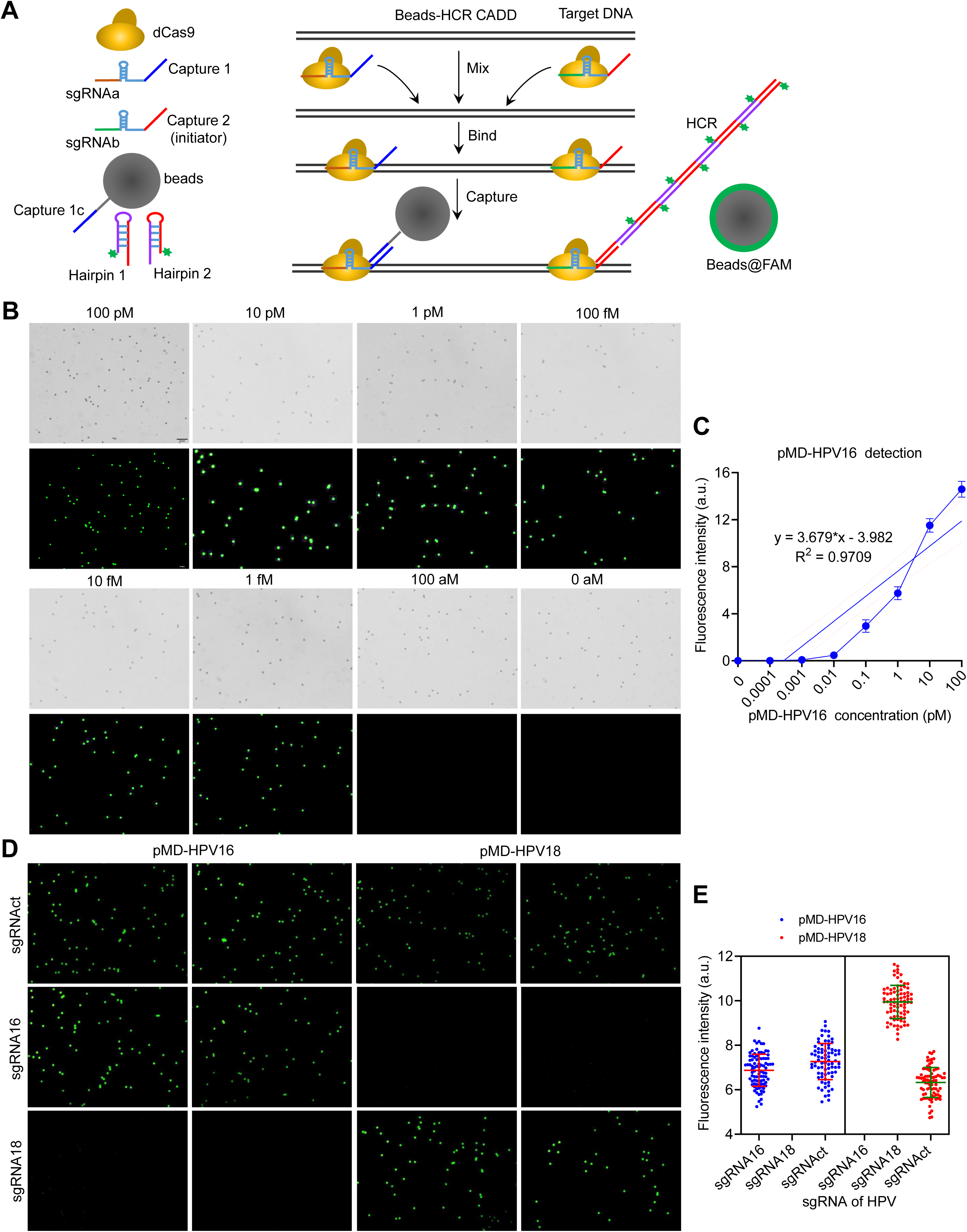
Beads-HCR CADD principle and pilot detection of HPV DNA. **A**. Schematic of Beads-HCR CADD. HCR, Hybridization Chain Reaction; Beads@FAM, microspheres with FAM fluorescence. **B** and **C**. Detection of different concentrations of pMD-HPV16 with sgRNA16. **B**. Beads fluorescent image. **C**: Quantitative analysis of beads fluorescence. There is a linear correlation between DNA concentration and fluorescence intensity in the range of 100 pM to 10 fM. **D** and **E**. Detection of pMD-HPV16 and pMD-HPV18 (each 1 pM) with sgRNA16, sgRNA18 and sgRNAct. **D**. Beads fluorescent image. **E**. Quantitative analysis of beads fluorescence. SgRNAct (sgRNA cocktail) is an equimolar mixture of sgRNAsp targeting 15 hrHPVs.

To investigate whether the designed HCR reaction is feasible, we tested the prepared Hairpin 1 and Hairpin 2 in liquid-phase HCRs with sgRNAa plus sgRNAb, sgRNAa, sgRNAb, and initiator oligo, respectively. The results show that HCR reaction was only initiated by sgRNAb (Fig. S1), indicating that the capture sequence of sgRNAb annealed with Hairpin 1. The oligo Initiator (Table S3) is a positive control that can also anneal with Hairpin 1 to initiate HCR (Fig. S1).

With the reliable Hairpins and sgRNAb, we first detected the HPV16 DNA (pMD-HPV16) with Beads-HCR CADD using sgRNAs targeting HPV16 (sgRNA16). The results show that pMD-HPV16 can be quantitatively detected by the method (Figs. 1B and 1C). To investigate the specificity of Beads-HCR detection, we then detected pMD-HPV16 and pMD-HPV18 with Beads-HCR CADD using sgRNA16, sgRNA18, and sgRNAct, respectively. SgRNAct is an equimolar mixture of sgRNAs of 15 hrHPVs. The results show that the two genotypes of HPVs can be specifically by the method (Figs. 1D and 1E).

To further explore the feasibility and specificity of Beads-HCR CADD to test HPVs, we detected all clinical hrHPVs with the method. Fifteen genotypes of hrHPVs (pMD-HPV16, 18, 31, 33, 35, 39, 45, 51, 52, 56, 58, 59, 66, 68, and 73) were detected with various sgRNAsp and sgRNAct, in which sgRNAsp means sgRNA specific to a particular genotype of HPV (such as sgRNA16). The empty vector pMD was used as a negative control. The results indicate that each HPV can be specifically detected by its cognate sgRNAsp and sgRNAct (Fig. 2, Fig.3, and supplementary File 1). The negative control pMD was not detected by any sgRNAs. These data indicate that the designed sgRNAs have high specificity when detecting 15 hrHPVs with Beads-HCR CADD.

**Fig. 2.**
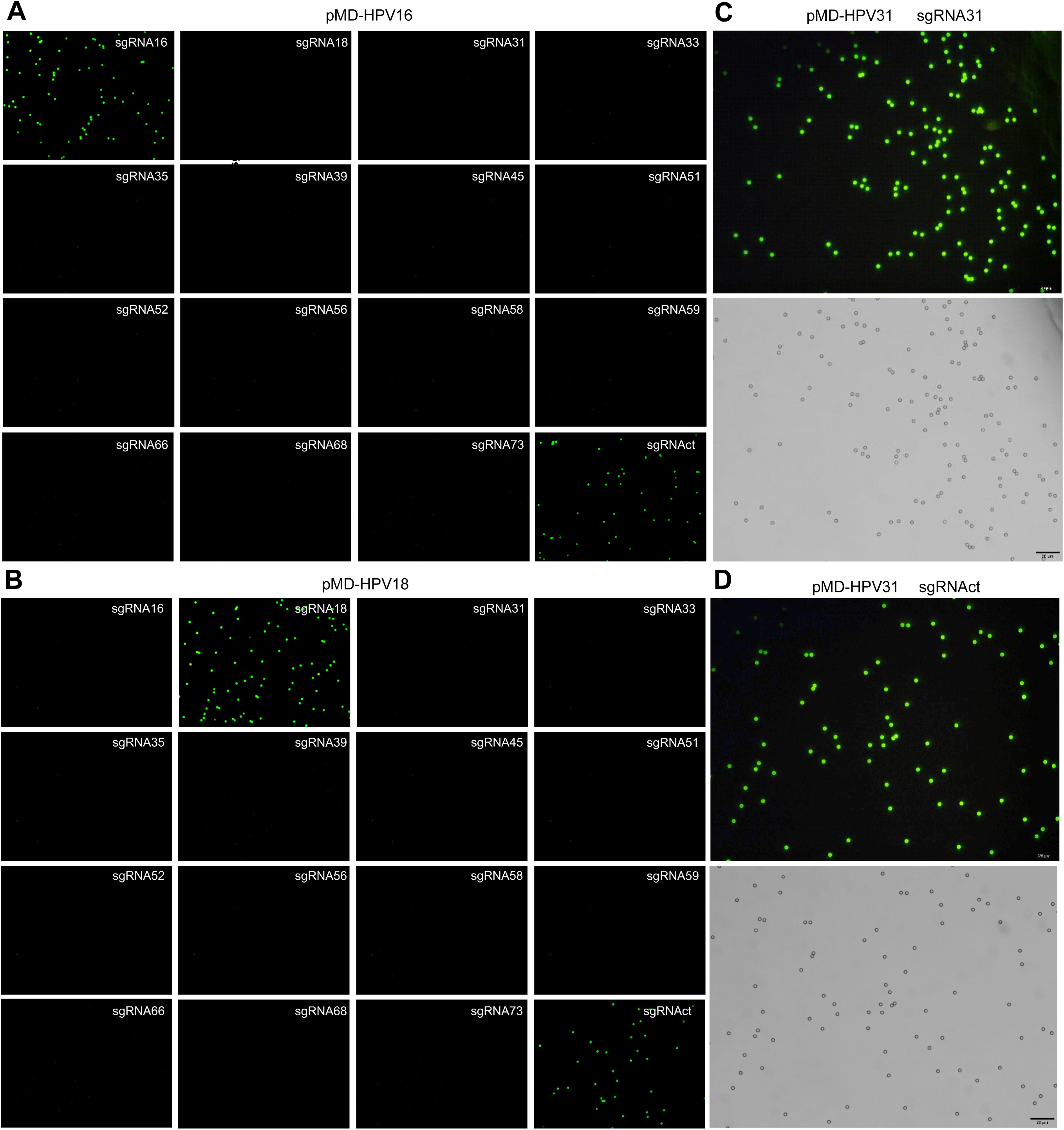
Detection of hrHPVs with Beads-HCR CADD. **A** and **B**. Detection of pMD-HPV16 (**A**) and pMD-HPV18 (**B**) with various sgRNAsp and sgRNAct. Beads fluorescent images are shown. **C** and **D**. Beads fluorescent and light images of pMD-HPV31 detected with sgRNA31 and sgRNAct.

**Fig. 3.**
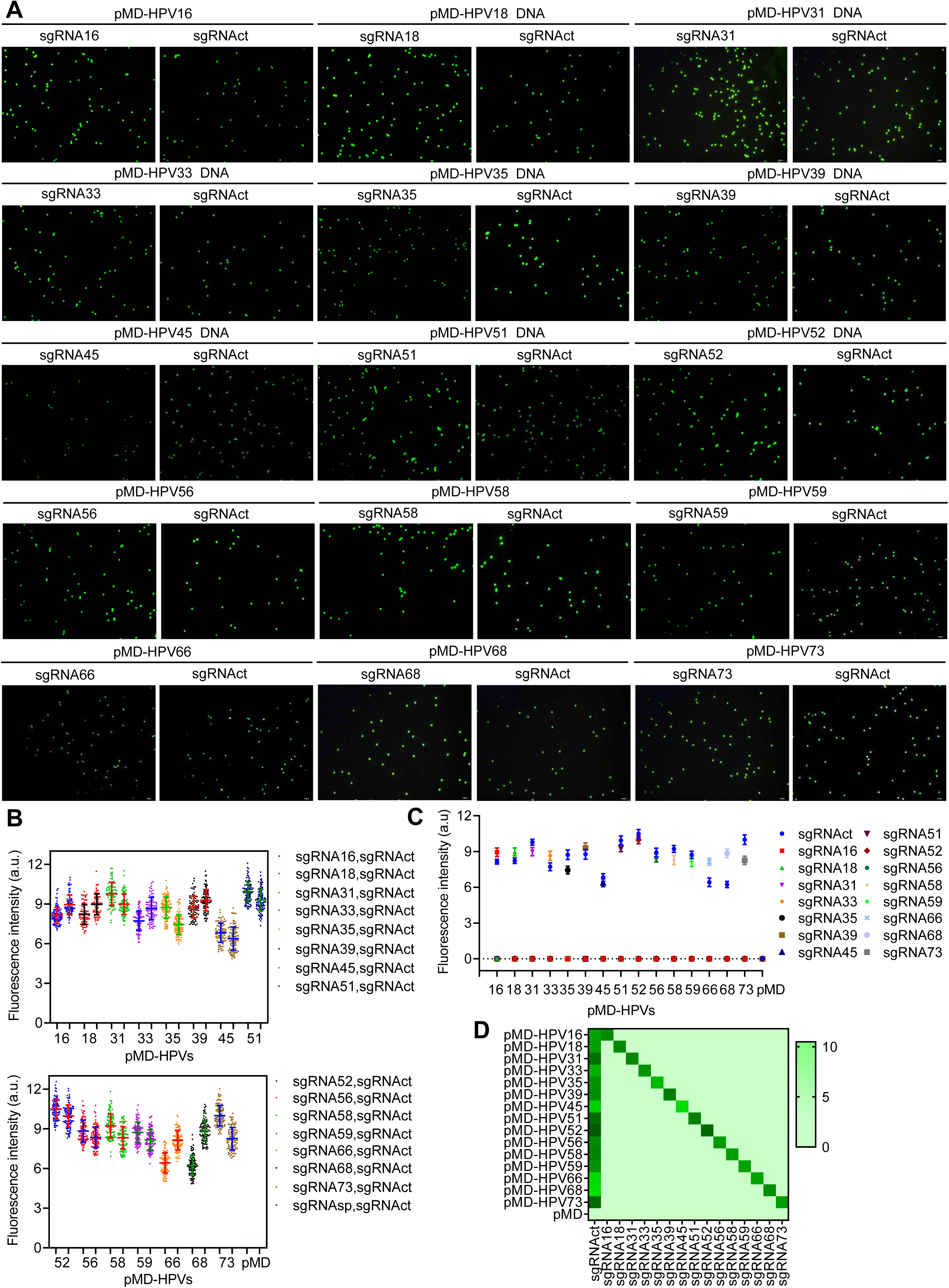
Detection of hrHPVs with Beads-HCR CADD. 15 hrHPVs (pMD-HPV16, 18, 31, 33, 35, 39, 45, 51, 52, 56, 58, 59, 66, 68, and 73) were detected with various sgRNAsp (sgRNA16, 18, 31, 33, 35, 39, 45, 51, 52, 56, 58, 59, 66, 68, and 73) and sgRNAct. **A**. Beads fluorescent image. Only the fluorescent images of each hrHPV detected with its cognate sgRNAsp and sgRNAct are shown here. The fluorescent images of all hrHPVs detected with various sgRNAsp and sgRNAct are shown in File 1. **B**. Quantitative analysis of beads fluorescent images of each hrHPV detected with its cognate sgRNAsp (left) and sgRNAct (right). **C**. Quantitative analysis of beads fluorescent images of each hrHPV detected with various sgRNAsp and sgRNAct. **D**. Quantitative analysis of beads fluorescent images of each hrHPV detected with various sgRNAsp and sgRNAct (heat map of average fluorescence intensity).

In order to investigate whether Beads-HCR CADD can be used to detect HPV DNA in human gDNA, we next detected gDNAs of three cervical cancer cell lines using sgRNA16, sgRNA18 and sgRNAct. HeLa and SiHa cells are known with HPV18 and HPV16 infection, respectively, and C-33a is known without HPV infection. The results indicate that the HeLa gDNA was detected by sgRNA18 and sgRNAct, the SiHa gDNA was detected by sgRNA16 and sgRNAct, and C-33a gDNA was not detected by any sgRNA (Fig. S2). These results indicate that Beads-HCR CADD is qualified for detection of more complicated DNA sample than HPV plasmid.

In order to investigate whether Beads-HCR CADD can be used to detect clinical sample, we then detected 31 clinical DNA samples with sgRNAsp of 15 hrHPVs and sgRNAct. The results are shown as Fig. 4 and supplementary File 2. We compared the HPV detection results of these 31 clinical samples tested by Beads-HCR CADD and PCR-reverse dot hybridization method that was performed by Jinling Hospital (Fig. 4C). In comparison with the hospital tests, the hrHPV infection (yes or no) and genotype are accurately detected by Beads-HCR CADD with 100% sensitivity and specificity. Importantly, the Beads-HCR CADD also found multiple infections in samples 2, 11, and 37 that were not detected by the PCR-reverse dot hybridization (Fig. 4C). These multiple infections were confirmed by a PCR re-detection (PCR-rd) (Fig. S3), in which HPV45 and HPV59 infection were detected by PCR using primers specific to the two HPVs (Supplementary method and Table S3). These results indicate that Beads-HCR CADD can be used to detect hrHPV infections in clinical samples with high sensitivity and specificity.

**Fig. 4.**
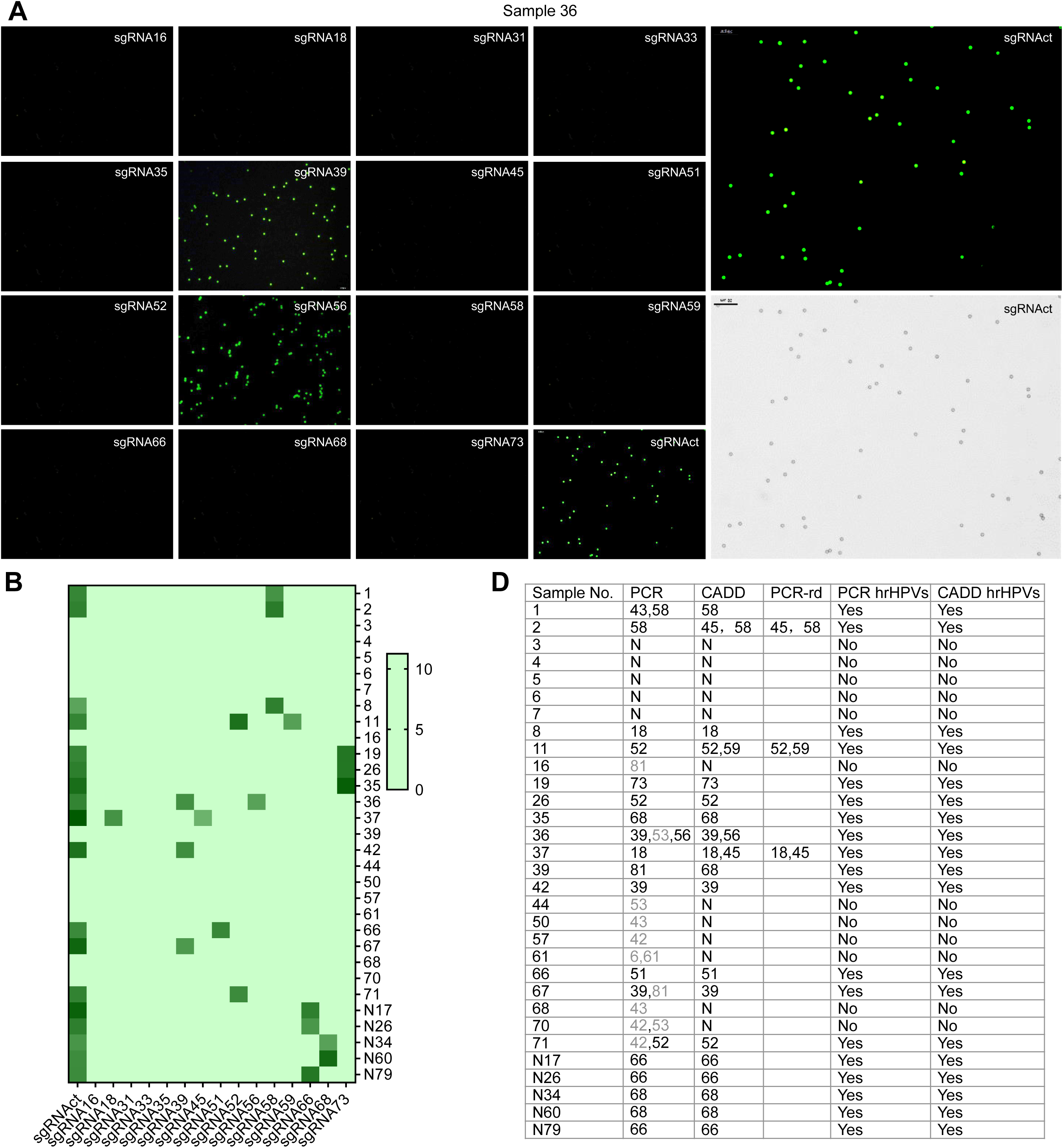
Test of the first batch of clinical cervical samples with Beads-HCR CADD. Totally 31 clinical samples were detected with various sgRNAsp and sgRNAct. The gDNA amount used for each detection was 150 ng. **A**. Beads fluorescent image. Only the fluorescent image of the Sample 36 detected with various sgRNAs are shown here. The beads fluorescent and light images of this sample detected with sgRNAct are shown in large size. The fluorescent images of other clinical samples are shown in File 2. **B**. Quantitative analysis of beads fluorescent images of 31 clinical samples detected with Beads-HCR CADD (heat map of average fluorescence intensity). **C**. Comparison of PCR and CADD test results of HPV infection in 31 clinical samples. PCR test was completed by Jinling Hospital. PCR-rd: HPV45 in Samples 2 and 37 and HPV59 in Sample 11 were detected again by specific PCR amplification (Fig.S3). The genotype in grey is not regarded as hrHPVs.

Finally, to investigate whether DNA other than virus DNA could be detected by Beads-HCR CADD, we also detected two types of other DNA. One is bacterium DNA and the other is human oncogenic DNA (Supplementary methods). The sgRNAs targeting T7 RNA polymerase DNA and oncogenic telomerase reverse transcriptase (TERT) promoter were designed (Table S1). The results indicate that the T7 RNA polymerase DNA fragment could be quantitatively detected by Beads-HCR (Fig. S4A and S4B). In addition, the subsequent detection of gDNA from two different E. coli (DH5α and BL21) indicate that the T7 RNA polymerase DNA in BL21 gDNA could be also specifically and quantitatively detected by Beads-HCR CADD (Fig. S4C and S4D). The DH5α gDNA that contains no T7 RNA polymerase DNA did not produce fluorescence signal even at the highest amount (Fig. S4C and S4D). The detection of TERT promoter DNA indicate that the ontogenetic TERT promoter can be specifically detected by Beads-HCR CADD using a sgRNAs targeting mutated TERT promoter (Fig. S5). The mutant TERT promoter causes expression of telomerase, which results in malignant cell proliferation in more than 90% of cancers. It should be noted that the detected TERT promoter has only one base difference between the wild-type and mutant genotype (Fig. S6), indicating that CADD has high specificity that can discriminate single nucleotide polymorphisms (SNP).

### 3.2. Beads-ELISA CADD detection

Because Beads-HCR CADD is dependent on fluorescent microscope, we then expect to realize a CADD with visual readout. We therefore designed a Beads-ELISA form of CADD (Fig. 5A). In this format of CADD, after dCas9-sgRNA binds target DNA and the dCas9-sgRNA-DNA complex is captured on beads surface via sgRNAa, a biotinylated oligonucleotide is captured by sgRNAb. The HRP-labeled streptavidin is then associated with biotin. Finally a soluble chromogenic substrate TMB is used to develop color signal. The detection results can thus be read either qualitatively with naked eyes or quantitatively with microplate reader.

**Fig. 5.**
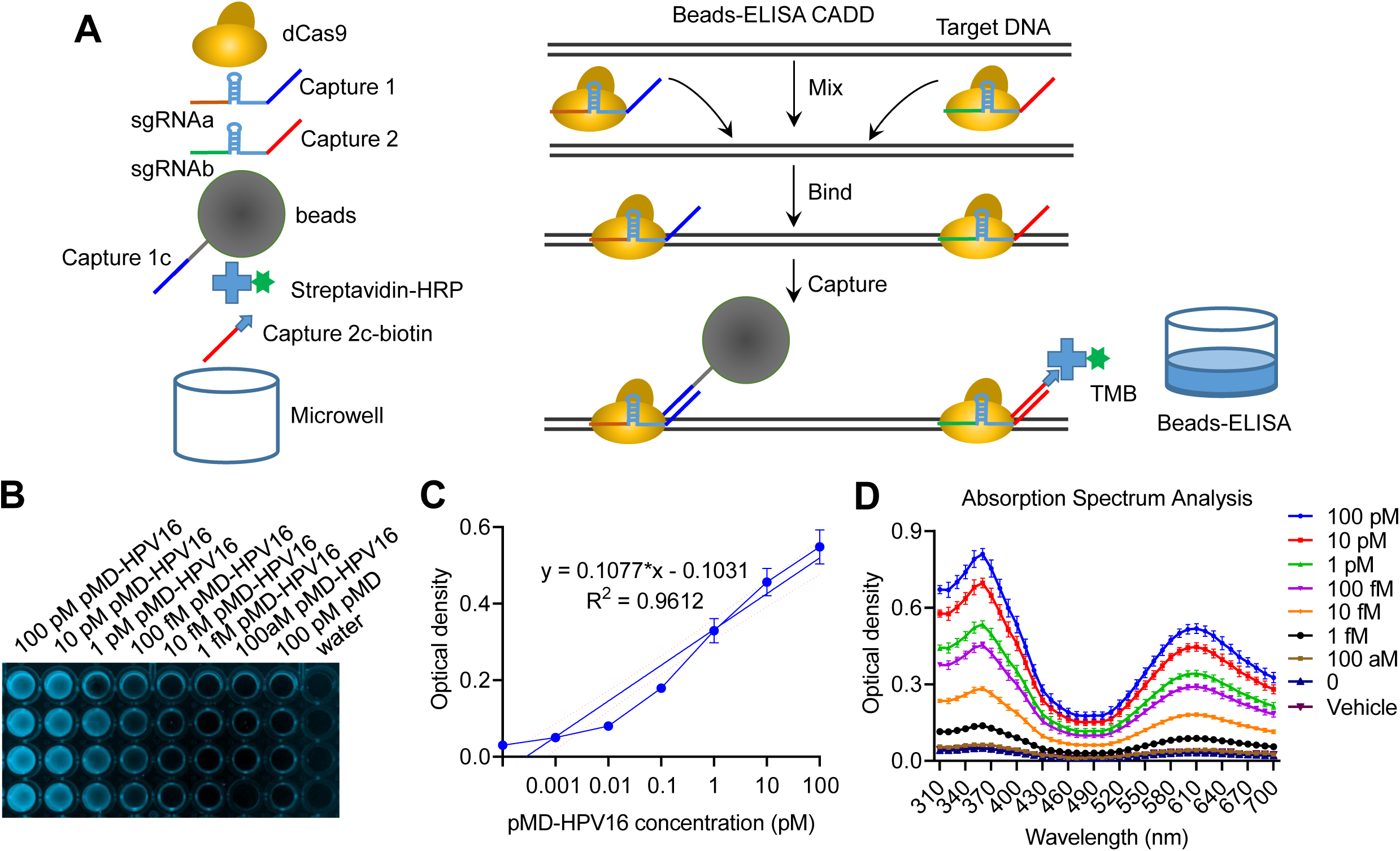
Beads-ELISA CADD principle and pilot detection of HPV. **A**. Schematic of the Beads-ELISA CADD. **B**. Detection of different concentrations of pMD-HPV16 with sgRNA16. The image shows the TMB-developed microplate. Each column is four replicates. **C**. Quantitative analysis of optical absorbance. There is a linear correlation between DNA concentration and absorbance in the range of 100 pM to 1 fM. **D**. Absorption spectrum measured with a microplate reader.

As a pilot assay, we first detected pMD-HPV16 with Beads-ELISA CADD. The results indicate that pMD-HPV16 can be quantitatively detected by this method (Fig. 5B–5D). In this assay, pMD was also used as a negative control. It cannot produce color even at the highest concentration (100 pM). We then detected 15 hrHPVs with this method using various sgRNAsp and sgRNAct. The results indicate that each hrHPV were specifically detected by its cognate sgRNAsp and sgRNAct (Fig. 6A and 6B). The negative control pMD was not detected by any sgRNA. To check if this method can detect multiple infections, we subsequently detected seven mixtures of two different hrHPVs using various sgRNAsp and sgRNAct. The results reveal that the simulated multiple infections were specifically detected by this method (Figs.6C and 6D). To further check the detection specificity, we finally detected several hrHPVs and clinical DNA samples with this method. The results demonstrate that each hrHPV was detected by its cognate sgRNAsp and sgRNAct (Fig. 6E); however, all single or mixed clinical DNA samples were not detected by any sgRNA (Fig. 6E). Because the clinical DNA samples were selected from the Beads-HCR CADD-detected samples, we focused on investigating if false positive can be produced by sgRNA when detecting clinical samples. The results reveal that all selected clinical samples were not detected by any sgRNA, further indicating the high specificity of this method.

**Fig. 6.**
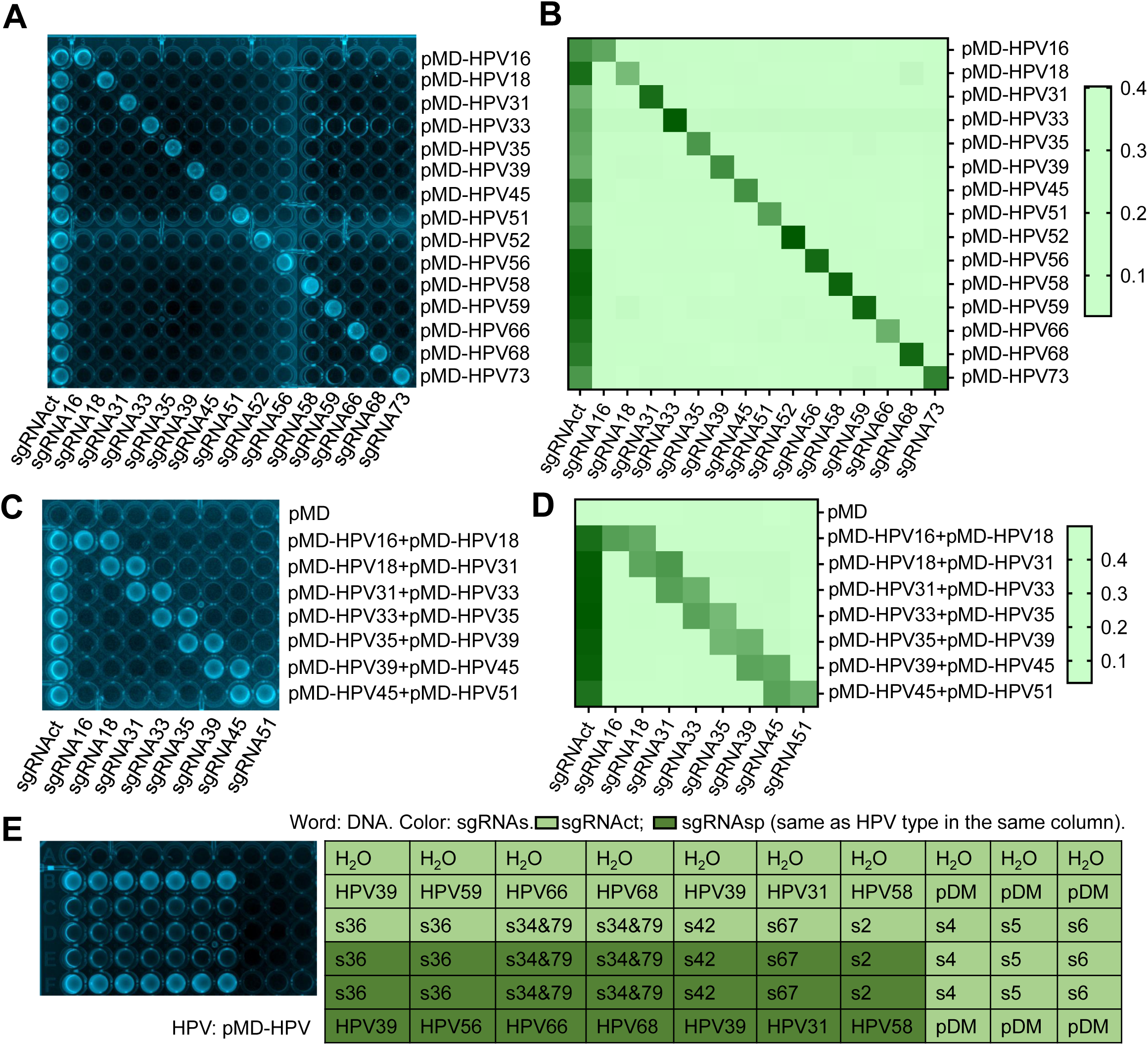
Detection of hrHPV with Beads-ELISA CADD. A and B. Detection of 15 hrHPVs (pMD-HPV16, 18, 31, 33, 35, 39, 45, 51, 52, 56, 58, 59, 66, 68, and 73) with various sgRNAsp and sgRNAct. **A**. Beads-ELISA microplate image. **B**. Quantified absorbance (heat map of absorbance). **C** and **D**. Detection of mixtures of two different HPV DNAs with Beads-ELISA CADD with various sgRNAsp and sgRNAct. **C**. Beads-ELISA microplate image. **D**. Quantified absorbance (heat map of absorbance). The pMD plasmid was used as a negative control. **E**. Detection of various HPV plasmids and clinical samples with various sgRNAsp and sgRNAct. Several pMD-HPVs and clinical samples (150 ng) were detected. The clinical samples were selected from the first batch of clinical samples detected by the Beads-HCR CADD.

### 3.3. Microplate-ELISA CADD detection

To further simplify the Beads-ELISA CADD and also try a new solid support other than beads, we designed a Microplate-ELISA form of CADD (Fig.7A). In this method, a capture oligonucleotide is coupled in microplate. The dCas9-sgRNA-DNA complex will be captured in microplate through annealing between the 3’-end capture sequence of sgRNAa and the capture oligonucleotide. The signal reporting system is completely the same as the Beads-ELISA CADD.

To verify this method, we first detected pMD-HPV16 using sgRNA16. The results indicate that pMD-HPV16 can be quantitatively detected by this method (Figs. 7B and 7C). The negative control pMD did not generate a signal at the highest concentration (Figs. 7B and 7C). We then detected 33 new clinical samples with this method using various sgRNAsp and sgRNAct. The results reveal that all samples were detected by this method (Fig. 8A and 8B). In comparison with the results obtained by hospital tests using the PCR-reverse dot hybridization, the Microplate-ELISA CADD accurately detected the infections of 15 hrHPVs (yes or no) and genotypes with 100% sensitivity and specificity. Additionally, the Microplate-ELISA CADD also identified more multiple infections in Samples 2, 5, 15, and 33. To confirm the test results, we re-detected the five samples with Microplate-ELISA. The results confirm that the Microplate-ELISA CADD can more accurately identify multiple infections in clinical samples than the current method used in clinics (Fig. 8D and 8E).

**Fig. 7.**
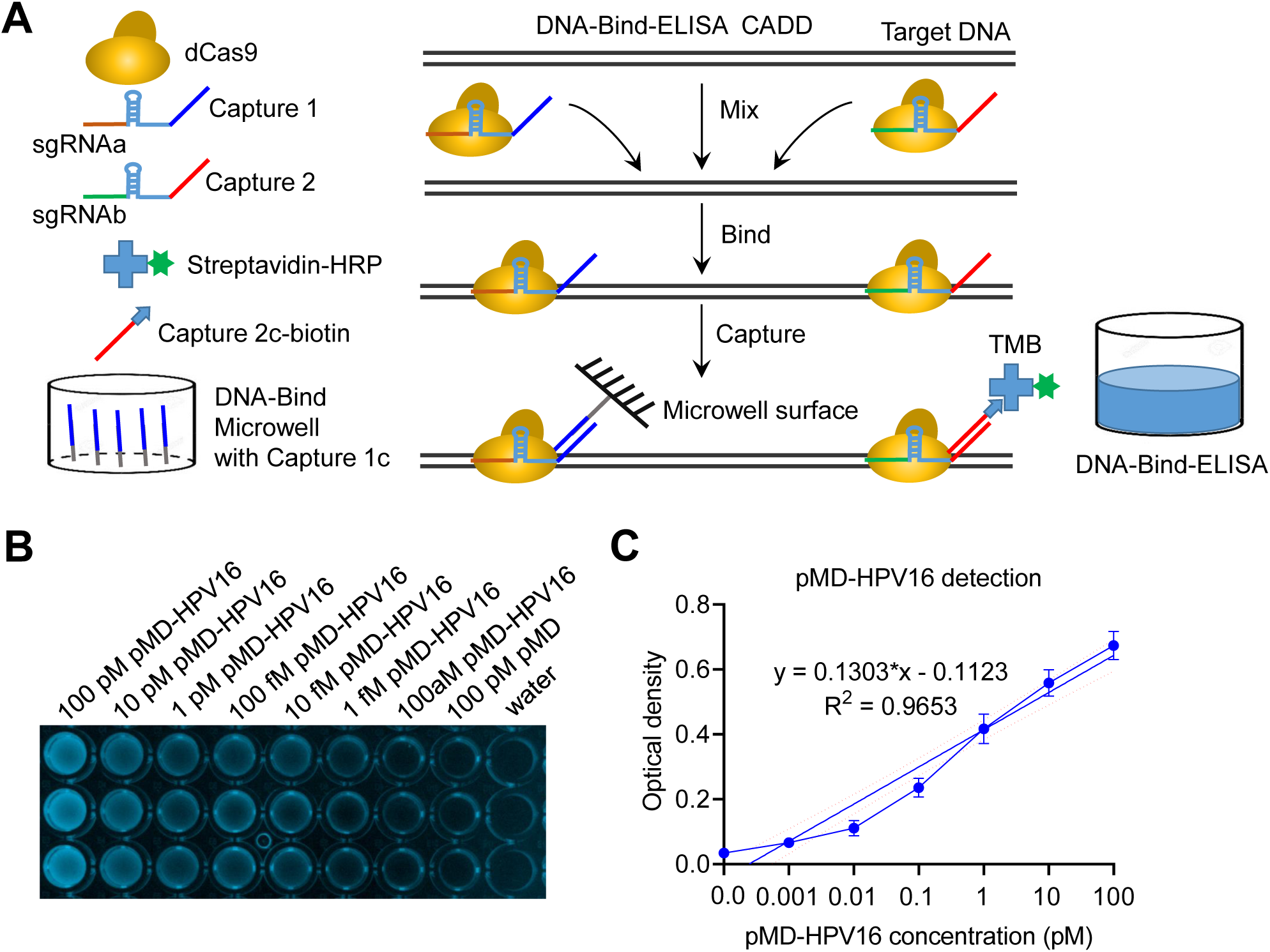
Microplate-ELISA CADD principle and pilot detection of HPV. **A**. Schematic of Microplate-ELISA CADD. B. Detection of different concentrations of pMD-HPV16 with sgRNA16. **B**. Image of TMB-developed microplate. Each column is three replicates. **C**. Quantitative analysis of optical absorbance. There is a linear correlation between DNA concentration and absorbance in the range of 100 pM to 1 fM.

**Fig. 8.**
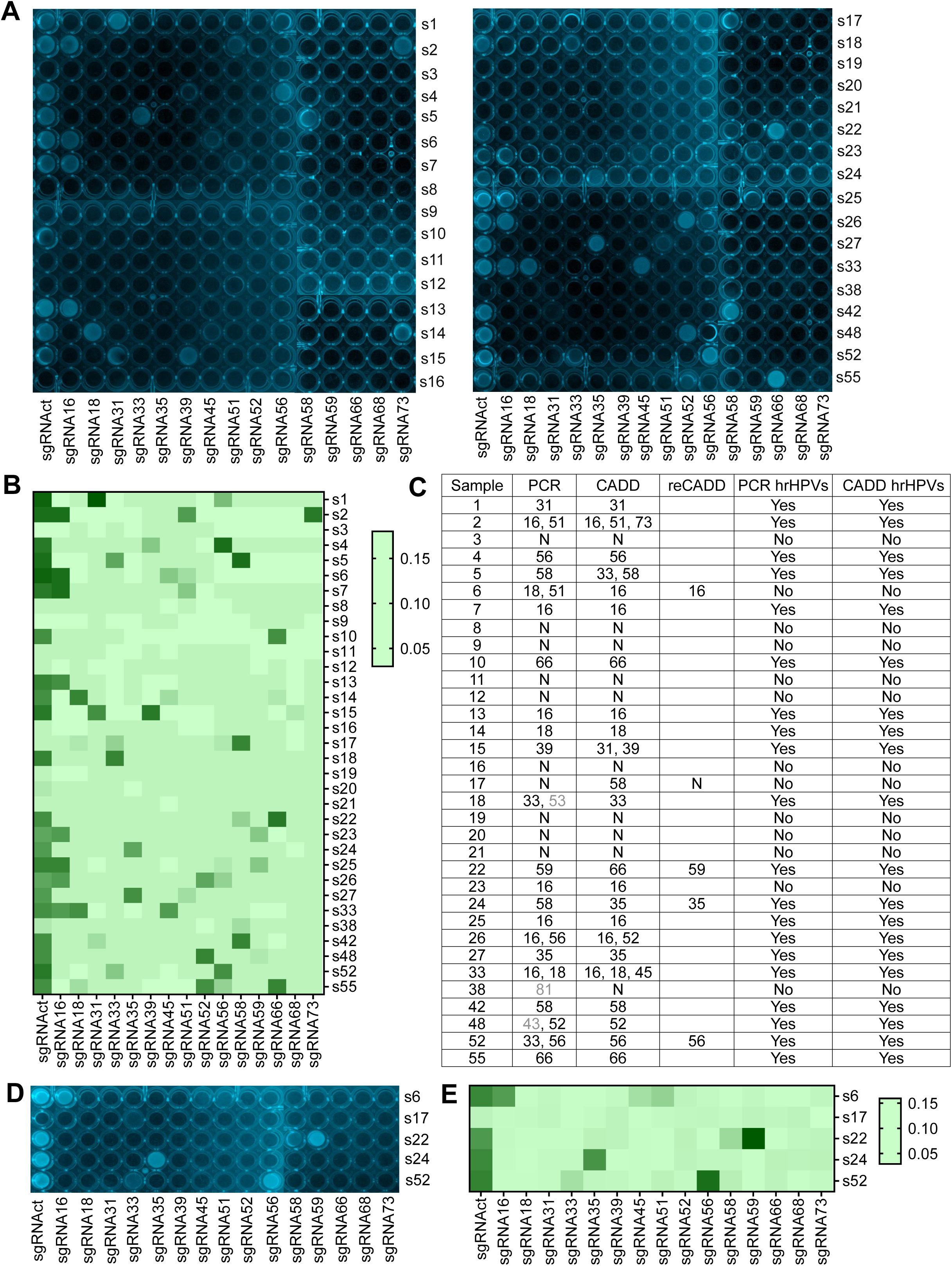
Detection of the second batch of clinical cervical samples with Microplate-ELISA CADD. Totally 33 clinical samples were detected with various sgRNAsp and sgRNAct. The amount of clinical samples used for each test was 150 ng. **A**. Microplate-ELISA microplate imaging. **B**. Quantitative analysis of Microplate-ELISA microplate image (heat map of absorbance). **C**. Comparison of PCR and CADD test results of hrHPV infection in 33 clinical samples. PCR test was completed by Jinling Hospital. Several samples were re-detected with Microplate-ELISA CADD (reCADD). The genotype in grey is not regarded as hrHPVs. **D** and **E**. Re-detection of HPV infection in 5 samples with Microplate-ELISA CADD. **D**. Microplate-ELISA microplate imaging. **E**. Quantified absorbance (heat map).

## 4. Discussion

In this study, we designed a new CRISPR/Cas9-based DNA detection technique and validated it by detecting three types of DNA including bacteria DNA, human cancer cell DNA and virus DNA. We also verified three forms of CADD method by detecting 15 hrHPV plasmids and as many 64 clinical cervical samples. These investigations demonstrate the feasibility and reliability of CADD method. In comparison, CADD has its unique advantages over the current CRISPR-based nucleic acid detection methods. Different from the current widely known CRISPR-based nucleic acid detection methods that mainly rely on Cas enzymes with collateral cleavage activity such as Cas13a, Cas12a, Cas12b, and Cas14, we develops a new DNA detection method based on a most widely used Cas protein, CRSIPR/Cas9, that has no collateral cleavage activity. CADD relies on neither the specific enzymatic activity nor non-specific collateral cleavage activity of CRISPR/Cas proteins as all the current CRISPR-based methods. Therefore, CADD is mechanistically distinct from the current CRISPR-based methods (such as SHERLOCK, DECTECTR, and HOLMES). CADD employs dCas9 that has no enzymatic activity. CADD uses the dCas9-sgRNA as a DNA-binding complex with high sequence specificity.

This study indicates that CADD can be used to detect and type various target DNA molecules with high simplicity, sensitivity and specificity. Although the current CRISPR-based methods are reported to have ultrahigh detection sensitivity (aM), they are all challenged by tedious pre-treatments to DNA/RNA samples including various amplifications using RPA, LAMP, PCR, asymmetric PCR and in vitro transcription. These pre-amplifications not only complicate the detection process and increase the detection time and cost, but also may increase false negative, because pre-amplification with various DNA polymerases may introduce mutations into DNA that is then enzymatically detected by various Cas proteins. Importantly, amplicon spread and contamination is always a serious issue in all detection spots. CADD needs no any pre-treatment to DNA sample. Beads-HCR CADD is an enzyme-free method. Beads/Microplate-ELISA CADD only need a widely used enzyme HRP to develop signal. All three forms of CADD methods can be performed without depending on any modular heaters. Another advantage of CADD is free of traditional hybridization at high temperature. The whole CADD process is carried out at room temperature. The time-consuming and instrument-dependent hybridization and amplification are key limitations to the on-the-spot application of the current nucleic acid tests. Due to these advantages, CADD will have wide applications in future DNA detection.

The CADD signal readout can be versatile. In this study, two forms of CADD signal readouts were verified, fluorescent and colorimetric readout. The Beads-HCR CADD uses fluorescent readout by employing the fluorescently labeled HCR hairpins. The Beads/Microplate-ELISA uses colorimetric readout, in which the TMB color development catalyzed by streptavidin-coupled HRP is used as a visual readout. The Beads/Microplate-ELISA CADD is an instrument-free test. In addition, the Microplate-ELISA assay allows automatic measurement of hundreds of samples on standard plate reader in a high-throughput format. The whole detection process of Beads-HCR and Microplate-ELISA can be finished in 30 minutes, holding promise for rapid on-the-spot detection or point-of-care testing (POCT). In fact, other forms of CADD signal readouts can be realized by a few changes of the solid support and signal reporter, such as lateral flow readout, nano-gold colorimetric assay, and molecular beacon-HCR.

The high specificity of CADD has close relationship with the high sequence specificity of dCas9-sgRNA as a DNA-binding complex. Additionally, the high specificity of CADD is also dependent on its unique detection mechanism, in which a pair of sgRNAs commonly determine the final detection results, providing a double-insurance sequence-specific detection. This overcomes the potential false positive results resulted from potential off-target binding of one dCas9-sgRNA. This problem still challenges all current CRISPR-based methods including SHERLOCK, DECTECTR and HOLMES, in which only one-site target cutting can activate a collateral cleavage activity. Due to the signal amplification produced by collateral cleavage activity, detections based on such one-site activation may be prone to false positives.

HPV is a double-stranded DNA virus that is closely related to the pathogenesis of cervical cancer, anal cancer and other cancers (43). There are about 100 different types of HPV. According to different carcinogenic capabilities, HPV is divided into high-risk HPV (hrHPV) and low-risk HPV (lrHPV). The most common hrHPVs in the world are HPV16 and HPV18, which cause about 70% of cervical cancers (44,45). Other hrHPVs include HPV 31, 33, 35, 39, 45, 51, 52, 56, 58, 59, 66, 68, and 73. Because of its rich DNA polymorphism, HPV is a good experimental material for study DNA detection and typing technology. Therefore, we uses HPV DNA as a material to validate three forms of CADD method in this study. With the results obtained, this study validated a set of sgRNAs targeting hrHPVs and three forms of CADD technique that can be directly used to detect clinical samples. When applied to clinical test, the Beads-HCR test can be finished in 30 min, much faster than the current HPV clinical tests. If clinician only wants to know if a clinical sample is infected by hrHPVs (screening test), the sample can only be rapidly tested with sgRNAct. If the sample is then needed to identify hrHPV genotype, it can then be tested by various sgRNAsp. In addition, many cervical samples can be tested in high throughput with the Microplate-ELISA test by using the current available automated liquid handling systems.

## 5. Conclusion

We designed a new CRISPR/Cas9-based DNA detection technique. We then validated this technique by detecting three types of DNA including bacteria DNA, human cancer cell DNA and virus DNA. This technique has significant advantages over the current CRISPR-based nucleic acid detection techniques. Especially, we designed a set of sgRNAs that are specific to 15 hrHPVs. By detecting plasmids containing the L1 fragment of 15 hrHPVs and 64 clinical DNA samples with three forms of CADD method, we demonstrates that CADD provides a new simple and rapid HPV test technique.

## Supplementary information

The supplementary information includes Supplementary methods, Table S1–S3, Figures S1□S6, and File 1 and File 2. Supplementary methods include (i) Preparation of bacterial gDNA and T7 RNA polymerase DNA fragment; (ii) Preparation of TERT promoter fragments; and (iii) PCR verification of Beads-HCR test results of clinical samples. Table S1. SgRNAs designed and used in this study. Table S2. Primers used to prepared sgRNA template by PCR amplification. Table S3. Oligos and other PCR primers. Fig. S1. Evaluation of HCR reaction. Fig. S2. Detection of gDNA of cervical cancer cell lines using Beads-HCR CADD. Fig. S3. Detection of HPV45 in samples 2 and 37 and HPV59 in sample 11 using specific PCR amplification. Fig. S4. Detection of T7 RNA polymerase DNA using Beads-HCR CADD. Fig. S5. Detection of TERT promoter DNA using Beads-HCR CADD. Fig.S6. DNA sequences of wild-type and mutant TERT promoter. File 1 Characterization of specificity of sgRNAs for 15 high-risk HPV subtypes. File 2 Detection of clinical samples with the Beads-HCR CADD.

## Author Contributions

J.W. conceived the study and designed the experiments. The first author X.X. performed all CADD detections and data analysis. J.G., T.L. and N.L. prepared sgRNA and other reagents. W.L. and the sixth author X.X. performed the HPV test in hospital and provided the left clinical samples that were then detected by CADD. J.W. and X.X. wrote the manuscript with supports from all authors.

