## Supplementary information for "CRISPR-Assisted DNA Detection, a novel dCas9-based DNA detection technique"

##### Preparation of bacterial gDNA and T7 RNA polymerase DNA fragment

The bacteria BL21 and DH5a were cultured in LB liquid medium, and genomic DNA (gDNA) was extracted using a genomic DNA extraction kit (B610423; Sangon Biotech, Shanghai). The purified gDNA was sheared by ultrasound and used as a substrate for HCR detection. A T7 RNA polymerase DNA fragment was amplified from BL21 gDNA. The PCR reaction (50 μL) contained 1 × PrimeSTAR® HS, 0.2 μM T7-F (Table S3), 0.2 μM T7-R (Table S3), and 5 ng gDNA. The PCR program was 98 °C for 5 min; 28 cycles of 98 °C 10 s, 58 °C 15 s, 72 °C 1 min; and 72 °C for 5 min. The amplified product was 651 bp. This fragment was purified and quantified as a substrate for HCR detection.

##### Preparation of TERT promoter fragments

The TERT promoter fragment was amplified from the gDNA extracted from HepG2 cells. The PCR reaction (50 μL) contained 1 × PrimeSTAR® HS, 0.2 μM TERT-F (Table S3), 0.2 μM TERT-R (Table S3), and 20 ng gDNA. The PCR program was: 98 °C for 5 min; 98 °C 10 s, 58 °C 15 s, 72 °C 1 min, 28 cycles; 72 °C, 5 min. The amplified product is 235 bp. The fragment was purified by gel recovery. After sequencing, the target base (rs1327649395) in this sequence was the wild-type TERT promoter. A pair of point mutation primers PM-F and PM-R (Table S3) were designed. The framed base is the base for introducing the mutation. The wild-type TERT promoter is at CG, and the mutant TERT promoter is at TA. Two PCR amplifications were first performed by using 5 ng of wild-type TERT promoter fragment as template and TERT-F and PM-R, and PM-F and TERT-R as primers, respectively. Two PCR products were recovered by gel recovery. Using 20 ng each of the two products as template and TERT-F and TERT-R as primers, a mutated TERT promoter fragment was prepared by PCR, in which the first 5 cycles were carried out without primers and then 23 cycles were carried out with primers. The PCR product was purified with gel recovery. The mutation was verified by sequencing.

##### PCR verification of Beads-HCR test results of clinical samples

In order to verify the multiple infection of samples 2, 11 and 37 in the first batch of clinical samples, PCR primers were designed to specifically amplify a L1 fragment of HPV45 and HPV59, including HPV45-test-F, HPV45-test-R, HPV59-test-F, and HPV59-test-R (Table S3). The pMD-HPV45, pMD-HPV59, pMD, gDNAs of clinical samples and HeLa cells were used as templates to amplify the target sequence. The PCR reaction (50 μL) contained 1× Hieff ™ PCR Master Mix, 0.2 μM HPV45-test-F/HPV59-test-F, 0.2 μM HPV45-test-R/HPV59-test-R), and template DNA (20 ng gDNA and 1 ng plasmid). PCR program was 98 °C 5 min; 98 °C 10 s, 58 °C 15 s, 72 °C 1.5 min, 28 cycles; 72 °C, 5 min. The PCR products were detected using 1% agarose gel electrophoresis. The PCR products of HPV45 and HPV59 are 722 bp and 1208 bp, respectively.

**Table S1**. SgRNAs designed and used in this study

| Target gene | sgRNA | sgRNA pair | sgRNA sequence（5'–3'） |
|---|---|---|---|
| HPV16 L1 | sgRNA16 | sgRNA16a | TTAAGGAGTACCTACGACAT |
| | | sgRNA16b | GTATCTTCTAGTGTGCCTCC |
| HPV18 L1 | sgRNA18 | sgRNA18a | TGCTGCACCGGCTGAAAATA |
| | | sgRNA18b | GCATCATATTGCCCAGGTAC |
| HPV31 L1 | sgRNA31 | sgRNA31a | ACCTACCTCTAAACCAACAC |
| | | sgRNA31b | TCCTACACCTAGCGGCTCCA |
| HPV33 L1 | sgRNA33 | sgRNA33a | CTGAGAGGTAACAAACCTAT |
| | | sgRNA33b | AAGGAAAAGGAAGACCCCTT |
| HPV35 L1 | sgRNA35 | sgRNA35a | ACACAGACATATTTGTACTA |
| | | sgRNA35b | TCTTTAGGTTTTGGTGCACT |
| HPV39 L1 | sgRNA39 | sgRNA39a | GTCCCCCGTAGATACATTAT |
| | | sgRNA39b | GACCTATAGTAGGGCGCCTG |
| HPV45 L1 | sgRNA45 | sgRNA45a | GGGTCATATGTACTTGGCAC |
| | | sgRNA45b | ACGATATGTATCCACCAAAC |
| HPV51 L1 | sgRNA51 | sgRNA51a | ATCCTACCATTCTTGAACAG |
| | | sgRNA51b | ACAGGCTAAGCCAGATCCTT |
| HPV52 L1 | sgRNA52 | sgRNA52a | GGAATACCTTCGTCATGGCG |
| | | sgRNA52b | CAGTTGTTTTGTCACAGTTG |
| HPV56 L1 | sgRNA56 | sgRNA56a | TATTGGGTTATCCCCGCCAG |
| | | sgRNA56b | CATATTCCTCCACATGTCTA |
| HPV58 L1 | sgRNA58 | sgRNA58a | GCTACGAGTGGTATCAACCA |
| | | sgRNA58b | AATGACATATATACATACTA |
| HPV59 L1 | sgRNA59 | sgRNA59a | TAAGGGTCCTGTTTAACTGG |
| | | sgRNA59b | CTGGTAGGTGTGTATACATT |
| HPV66 L1 | sgRNA66 | sgRNA66a | CAATCAATACCTTCGCCATG |
| | | sgRNA66b | CCCAATAGAGGACGGTGACA |
| HPV68 L1 | sgRNA68 | sgRNA68a | CATAGACCCACTAGGCGAGG |
| | | sgRNA68b | TGATACGCAGCGATTGGTAT |
| HPV73 L1 | sgRNA73 | sgRNA73a | TGATAAGGAGCGCCTAGTAT |
| | | sgRNA73b | TAGGTTGTAGGCCTCCCTTA |
| T7 RNA Pol | sgRNAt7 | sgRNAt7a | CTGCGGTTGAAGCAATGAAC |
| | | sgRNAt7b | CCCACACTACAGTCTTACGA |
| TERT promoter | sgRNAtert | sgRNAterta | GGACCGCGCTCCCCACGTGG |
| | | sgRNAtertb | TCCCCGGCCCAGCCCCTTCC |

**Table S2**. Primers used to prepared sgRNA template by PCR amplification

| Name | Sequence (5'–3') |
|---|---|
| F1 | GTTTTAGAGCTAGAAATAGCAAGTTAAAATAAGGCTAGTCCGTTATCAACTTG |
| Ra | AAAAAAAAGCATCTGGTATTCGTAAGGTTCCGCACCGACTCGGTGCCACTTTTTC |
| Rb-1 | GTGGGAGTCGTCTGTAACATGAAGTAGCACCGACTCGGTGCCACTTTTTC |
| Rb-2 | AAAAAAAGGTATGTTTGAGGTTCCACTAGATGCACCGACTCGGTGCCACTTTTTC |
| sgRa | AAAAAAAAGCATCTGGTATTCGTAAGGTTC |
| sgRb-1 | GTGGGAGTCGTCTGTAACATGAAGTAG |
| sgRb-2 | AAAAAAAGGTATGTTTGAGGTTCCACTAG |
| 16L1a-F2 | TTAAGGAGTACCTACGACATGTTTTAGAGCTAGAAATAGCAAG |
| 16L1a-F3 | TTCTAATACGACTCACTATAG*TTAAGGAGTACCTACGACAT*G |
| 16L1b-F2 | GTATCTTCTAGTGTGCCTCCGTTTTAGAGCTAGAAATAGCAAG |
| 16L1b-F3 | TTCTAATACGACTCACTATAG*GTATCTTCTAGTGTGCCTCC*G |
| 18L1a-F2 | TGCTGCACCGGCTGAAAATAGTTTTAGAGCTAGAAATAGCAAG |
| 18L1a-F3 | TTCTAATACGACTCACTATAG*TGCTGCACCGGCTGAAAATA*G |
| 18L1b-F2 | GCATCATATTGCCCAGGTACGTTTTAGAGCTAGAAATAGCAAG |
| 18L1b-F3 | TTCTAATACGACTCACTATAG*GCATCATATTGCCCAGGTAC*G |
| 33L1a-F2 | CTGAGAGGTAACAAACCTATGTTTTAGAGCTAGAAATAGCAAG |
| 33L1a-F3 | TTCTAATACGACTCACTATAG*CTGAGAGGTAACAAACCTAT*G |
| 33L1b-F2 | AAGGAAAAGGAAGACCCCTTGTTTTAGAGCTAGAAATAGCAAG |
| 33L1b-F3 | TTCTAATACGACTCACTATAG*AAGGAAAAGGAAGACCCCTT*G |
| 35L1a-F2 | ACACAGACATATTTGTACTAGTTTTAGAGCTAGAAATAGCAAG |
| 35L1a-F3 | TTCTAATACGACTCACTATAG*ACACAGACATATTTGTACTA*G |
| 35L1b-F2 | TCTTTAGGTTTTGGTGCACTGTTTTAGAGCTAGAAATAGCAAG |
| 35L1b-F3 | TTCTAATACGACTCACTATAG*TCTTTAGGTTTTGGTGCACT*G |
| 45L1a-F2 | GGGTCATATGTACTTGGCACGTTTTAGAGCTAGAAATAGCAAG |
| 45L1a-F3 | TTCTAATACGACTCACTATAG*GGGTCATATGTACTTGGCAC*G |
| 45L1b-F2 | ACGATATGTATCCACCAAACGTTTTAGAGCTAGAAATAGCAAG |
| 45L1b-F3 | TTCTAATACGACTCACTATAG*ACGATATGTATCCACCAAAC*G |
| 51L1a-F2 | ATCCTACCATTCTTGAACAGGTTTTAGAGCTAGAAATAGCAAG |
| 51L1a-F3 | TTCTAATACGACTCACTATAG*ATCCTACCATTCTTGAACAG*G |
| 51L1b-F2 | ACAGGCTAAGCCAGATCCTTGTTTTAGAGCTAGAAATAGCAAG |
| 51L1b-F3 | TTCTAATACGACTCACTATAG*ACAGGCTAAGCCAGATCCTT*G |
| 52L1a-F2 | GGAATACCTTCGTCATGGCGGTTTTAGAGCTAGAAATAGCAAG |
| 52L1a-F3 | TTCTAATACGACTCACTATAG*GGAATACCTTCGTCATGGCG*G |
| 52L1b-F2 | CAGTTGTTTTGTCACAGTTGGTTTTAGAGCTAGAAATAGCAAG |
| 52L1b-F3 | TTCTAATACGACTCACTATAG*CAGTTGTTTTGTCACAGTTG*G |
| 56L1a-F2 | TATTGGGTTATCCCCGCCAGGTTTTAGAGCTAGAAATAGCAAG |
| 56L1a-F3 | TTCTAATACGACTCACTATAG*TATTGGGTTATCCCCGCCAG*G |
| 56L1b-F2 | CATATTCCTCCACATGTCTAGTTTTAGAGCTAGAAATAGCAAG |
| 56L1b-F3 | TTCTAATACGACTCACTATAG*CATATTCCTCCACATGTCTA*G |

| | |
|---|---|
| 58L1a-F2 | GCTACGAGTGGTATCAACCAGTTTTAGAGCTAGAAATAGCAAG |
| 58L1a-F3 | TTCTAATACGACTCACTATAG*GCTACGAGTGGTATCAACCA*G |
| 58L1b-F2 | AATGACATATATACATACTAGTTTTAGAGCTAGAAATAGCAAG |
| 58L1b-F3 | TTCTAATACGACTCACTATAG*AATGACATATATACATACTA*G |
| 59L1a-F2 | TAAGGGTCCTGTTTAACTGGGTTTTAGAGCTAGAAATAGCAAG |
| 59L1a-F3 | TTCTAATACGACTCACTATAG*TAAGGGTCCTGTTTAACTGG*G |
| 59L1b-F2 | CTGGTAGGTGTGTATACATTGTTTTAGAGCTAGAAATAGCAAG |
| 59L1b-F3 | TTCTAATACGACTCACTATAG*CTGGTAGGTGTGTATACATT*G |
| 31L1a-F2 | ACCTACCTCTAAACCAACACGTTTTAGAGCTAGAAATAGCAAG |
| 31L1a-F3 | TTCTAATACGACTCACTATAGACCTACCTCTAAACCAACAC |
| 31L1b-F2 | TCCTACACCTAGCGGCTCCAGTTTTAGAGCTAGAAATAGCAAG |
| 31L1b-F3 | TTCTAATACGACTCACTATAGTCCTACACCTAGCGGCTCCA |
| 39L1a-F2 | GTCCCCCGTAGATACATTATGTTTTAGAGCTAGAAATAGCAAG |
| 39L1a-F3 | TTCTAATACGACTCACTATAGGTCCCCCGTAGATACATTAT |
| 39L1b-F2 | GACCTATAGTAGGGCGCCTGGTTTTAGAGCTAGAAATAGCAAG |
| 39L1b-F3 | TTCTAATACGACTCACTATAGGACCTATAGTAGGGCGCCTG |
| 66L1a-F2 | CAATCAATACCTTCGCCATGGTTTTAGAGCTAGAAATAGCAAG |
| 66L1a-F3 | TTCTAATACGACTCACTATAGCAATCAATACCTTCGCCATG |
| 66L1b-F2 | CCCAATAGAGGACGGTGACAGTTTTAGAGCTAGAAATAGCAAG |
| 66L1b-F3 | TTCTAATACGACTCACTATAGCCCAATAGAGGACGGTGACA |
| 68L1a-F2 | CATAGACCCACTAGGCGAGGGTTTTAGAGCTAGAAATAGCAAG |
| 68L1a-F3 | TTCTAATACGACTCACTATAGCATAGACCCACTAGGCGAGG |
| 68L1b-F2 | TGATACGCAGCGATTGGTATGTTTTAGAGCTAGAAATAGCAAG |
| 68L1b-F3 | TTCTAATACGACTCACTATAGTGATACGCAGCGATTGGTAT |
| 73L1a-F2 | TGATAAGGAGCGCCTAGTATGTTTTAGAGCTAGAAATAGCAAG |
| 73L1a-F3 | TTCTAATACGACTCACTATAGTGATAAGGAGCGCCTAGTAT |
| 73L1b-F2 | TAGGTTGTAGGCCTCCCTTAGTTTTAGAGCTAGAAATAGCAAG |
| 73L1b-F3 | TTCTAATACGACTCACTATAGTAGGTTGTAGGCCTCCCTTA |
| T7_a_F2 | CTGCGGTTGAAGCAATGAACGTTTTAGAGCTAGAAATAGCAAG |
| T7_a_F3 | TTCTAATACGACTCACTATAGCTGCGGTTGAAGCAATGAAC |
| T7_b_F2 | CCCACACTACAGTCTTACGAGTTTTAGAGCTAGAAATAGCAAG |
| T7_b_F3 | TTCTAATACGACTCACTATAGCCCACACTACAGTCTTACGA |
| TERT-a-sgRNA-F2 | GGACCGCGCTCCCCACGTGGGTTTTAGAGCTAGAAATAGCAAG |
| TERT-a-sgRNA-F3 | TTCTAATACGACTCACTATAGGGACCGCGCTCCCCACGTGG |
| TERT-b-sgRNA-F2 | TCCCCGGCCCAGCCCCTTCCGTTTTAGAGCTAGAAATAGCAAG |
| TERT-b-sgRNA-F3 | TTCTAATACGACTCACTATAGTCCCCGGCCCAGCCCCTTCC |

Note: The sgRNAa template was prepared with primers F1, Ra and sgRa. The sgRNAb template was prepared with primers F1, Rb-1/Rb-2 and sgRb-1/sgRb-2. The template prepared with primers F1, Rb-1 and sgRb-1 was used in Beads-HCR detection. The template prepared with primers F1, Rb-2 and sgRb-2 was used in Beads-ELISA and Microplate-ELISA detections.

**Table S3**. Oligos and other PCR primers

| Name | Sequence | Usage |
|---|---|---|
| FAM-hairpin-1 | 5'-FAM-ACAGACGACTCCCACATTCTCCAGGTGGGAGTCGTCTGTAACATGAAGTA-3' | oligo |
| FAM-hairpin-2 | 5'-CTGGAGAATGTGGGAGTCGTCTGTTACTTCATGTTACAGACGACTCCCAC-FAM-3' | oligo |
| Initiator | 5'-TACTTCATGTTACAGACGACTCCCAC-3 ' | oligo |
| RE-flanking | 5'-Biotin-TTTTTTTGCATCTGGTATTCGTAAGGTTCCG-3 ' | oligo |
| Re-biotin | 5′-Biotin-TTTTTTGGTATGTTTGAGGTTCCACTAGAT-3′ | oligo |
| RE-$NH_2$ | 5'-$NH_2$-TTTTTTGCATCTGGTATTCGTAAGGTTCCG-3′ | oligo |
| T7-F | 5'-GAGTTCGGCTTCCGTCAACAAGTG-3 ' | primer |
| T7-R | 5'-GTCCAATTGAGACTCGTGCAACTG-3' | primer |
| TERT-F | 5'-AGTGGATTCGCGGGCACAGA-3 ' | primer |
| TERT-R | 5'-CAGCGCTGCCTGAAACTC-3' | primer |
| PM-F | 5'-CGGGTCCCCGGCCCAGCCCCTTCCGGGCCCTCCCAGCCCCTCC-3 ' | primer |
| PM-R | 5'-GGAGGGGCTGGGAGGGCCCGGAAGGGGCTGGGCCGGGGACCCG-3' | primer |
| HPV45-test-F | 5′-TTCTGTGGCCAGAGTTGTCA-3′ | primer |
| HPV45-test-R | 5′-ACAGTTGTTCACGGCGTAGG-3′ | primer |
| HPV59-test-F | 5′-TAGGTGTTGAAATCGGTCGGG-3′ | primer |
| HPV59-test-R | 5′-AGACTTGCGACGCTTAACAC-3′ | primer |

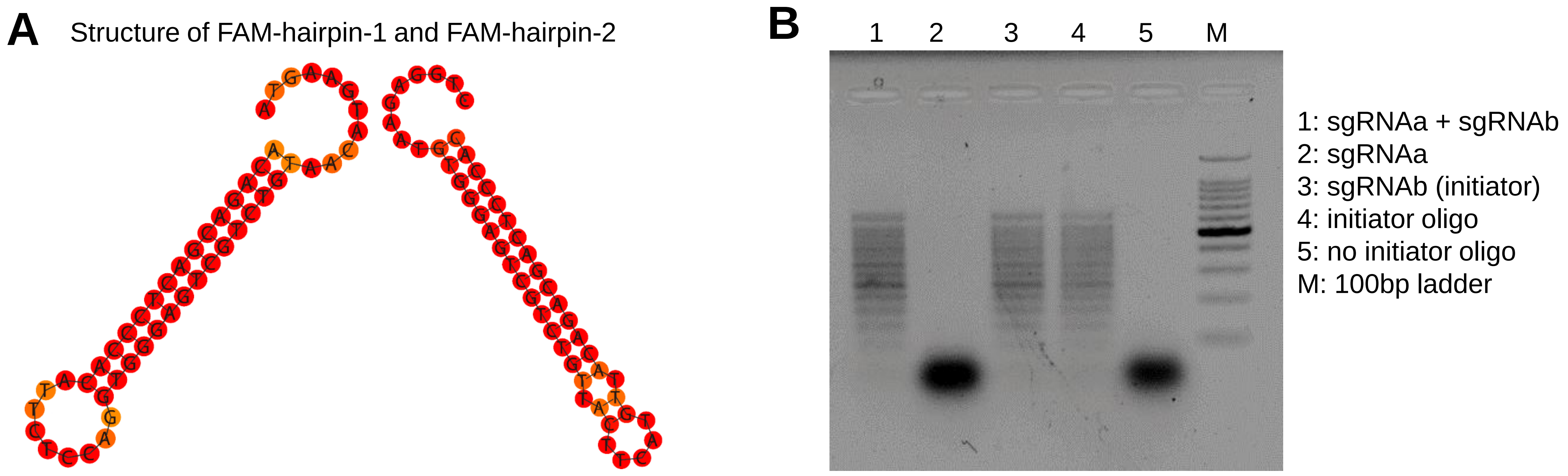


Hairpin 1 5′ -FAM-ACAGACGACTCCCACATTCTCCAGGTGGGAGTCGTCTGTAACATGAAGTA-3′

Hairpin 2 3′-FAM-CACCCTCAGCAGACATTGTACTTCATTGTCTGCTGAGGGTGTAAGAGGCT-5′

**Fig. S1**. Evaluation of HCR reaction. (**A**) The secondary structure of Hairpin 1 and Hairpin 2. The sequence and fluorescence modification of Hairpin 1 and Hairpin 2 were shown. (**B**) Liquid phase detection of HCR reaction. The sgRNAa and sgRNAb, sgRNAa, sgRNAb (initiator), or initiator oligo were added to the HCR reaction solution containing Hairpin 1 and Hairpin 2. The reaction products were detected by agarose gel electrophoresis. The reaction without the initiator oligo was used as a negative control.

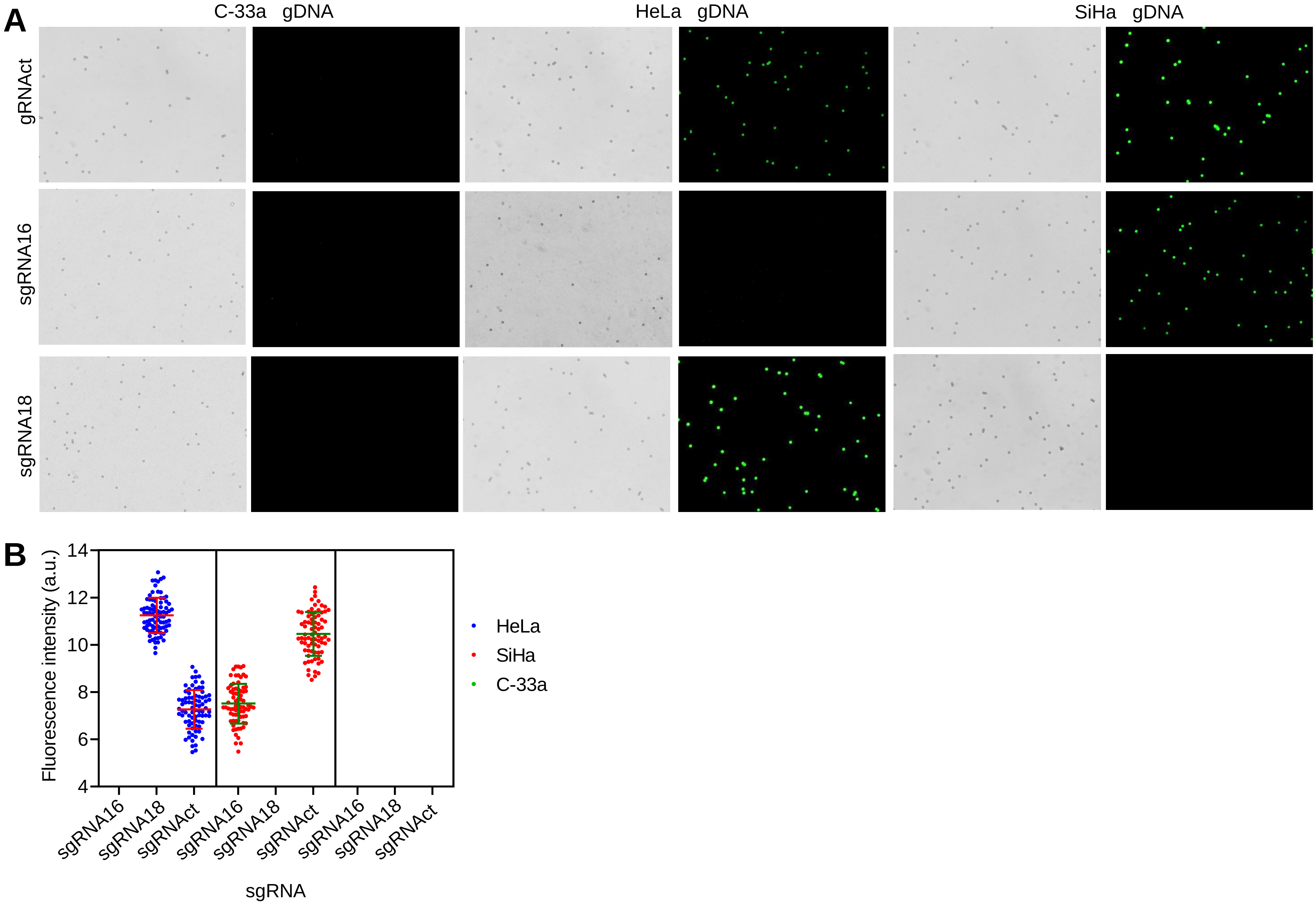


**Fig. S2**. Detection of gDNA of cervical cancer cell lines with Beads-HCR CADD. The gDNA of three cervical cancer cell lines were detected with sgRNA16, sgRNA18 and sgRNAct, respectively. (**A**) Beads fluorescence image. (**B**) Quantitative analysis of Beads fluorescence.

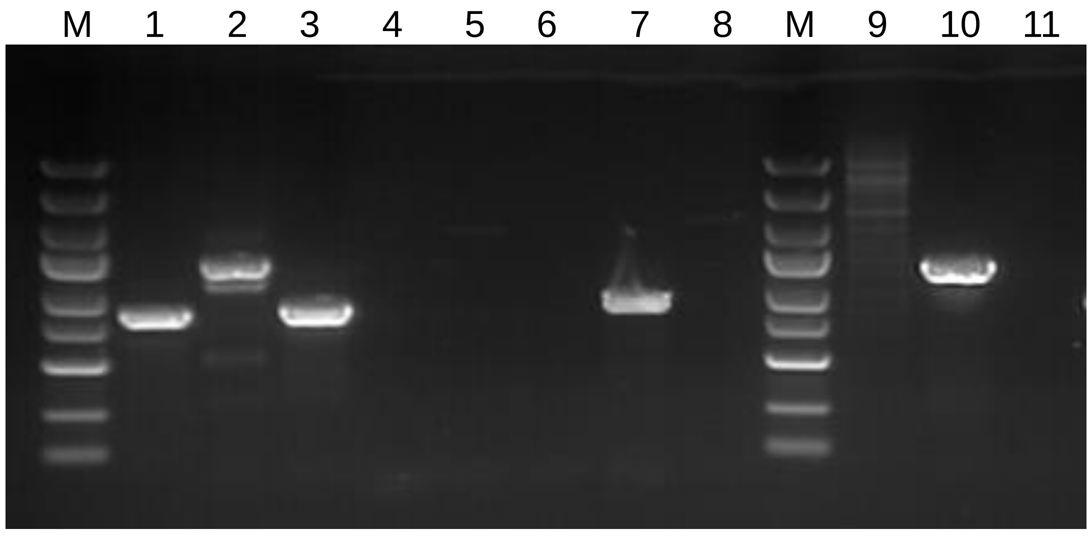


| Lane | 1 | 2 | 3 | 4 | 5 | 6 |
|---|---|---|---|---|---|---|
| Template | pMD-HPV45 | pMD-HPV59 | S2 | S3 | pMD | pMD |
| Primers | 45 | 59 | 45 | 45 | 45 | 59 |

| Lane | 7 | 8 | 9 | 10 | 11 |
|---|---|---|---|---|---|
| Template | S37 | HeLa gDNA | HeLa gDNA | S11 | S3 |
| Primers | 45 | 45 | 59 | 59 | 59 |

**Fig. S3**. Detection of HPV45 in Sample 2 and 37 and HPV59 in Sample 11 with specific PCR amplification. The HPV45- and HPV59-specific primers were used to amplify different DNA samples (template). The pMD-HPV45 and pMD-HPV45 were used as positive controls. The HeLa gDNA, pMD and the Sample 3 were used as a negative control. It can be seen that the Sample 2 and 37 were infected by HPV45 and the Sample 11 was infected by HPV59.

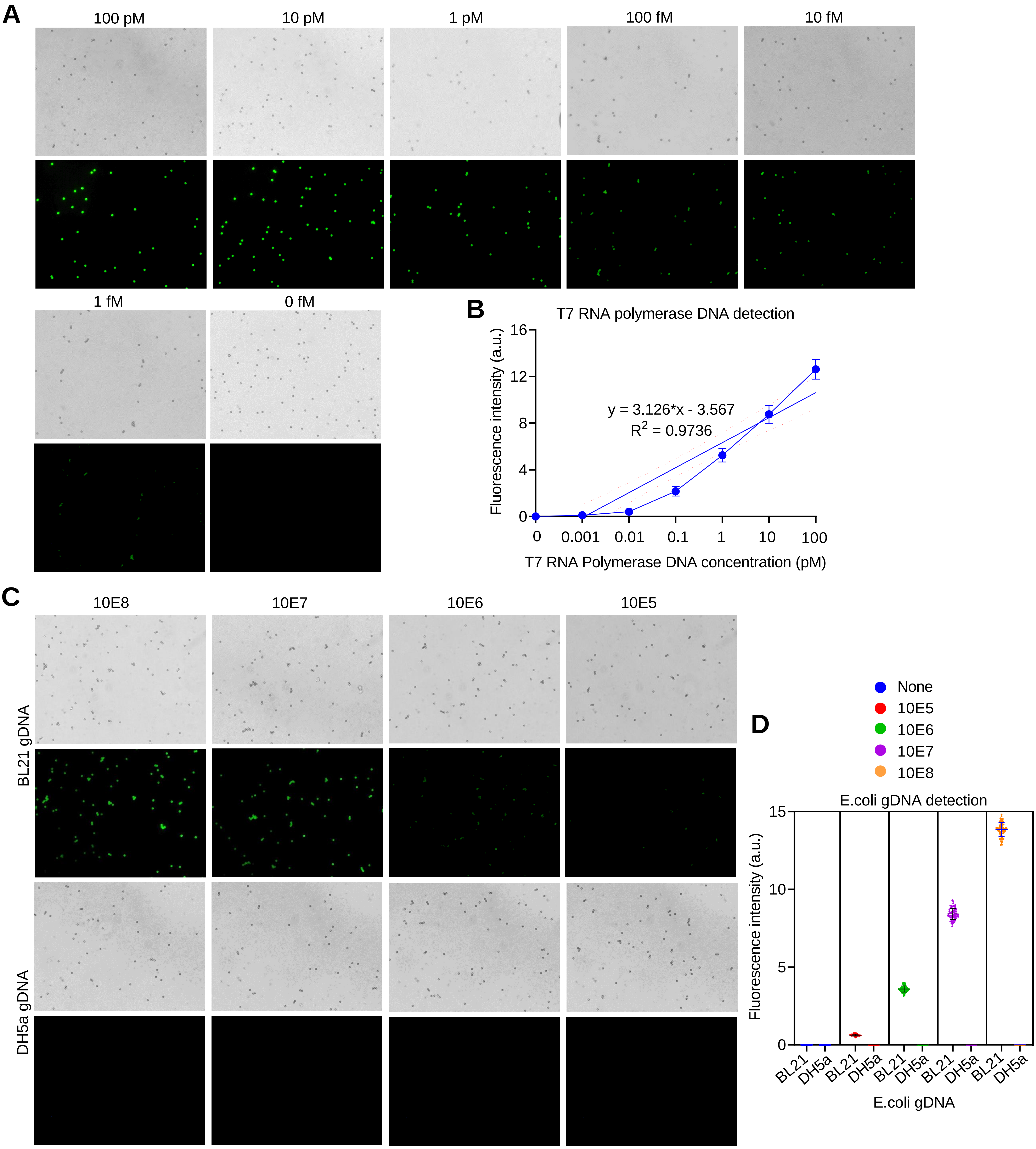


**Fig. S4**. Detection of T7 RNA polymerase DNA using Beads-HCR CADD. A and B. Detection of different concentrations of T7 RNA polymerase DNA fragment with sgRNAt7. (**A**) Beads fluorescence image. (**B**) Quantitative analysis of Beads fluorescence. There is a linear correlation between DNA concentration and fluorescence intensity in the range of 100 pM to 10 fM. (**C** and **D**) Detection of gDNA of BL21 and DH5α at different concentrations with sgRNAt7. (**C**) Beads fluorescence image. (**D**) Quantitative analysis of Beads fluorescence. Each view displays light and fluorescent images.

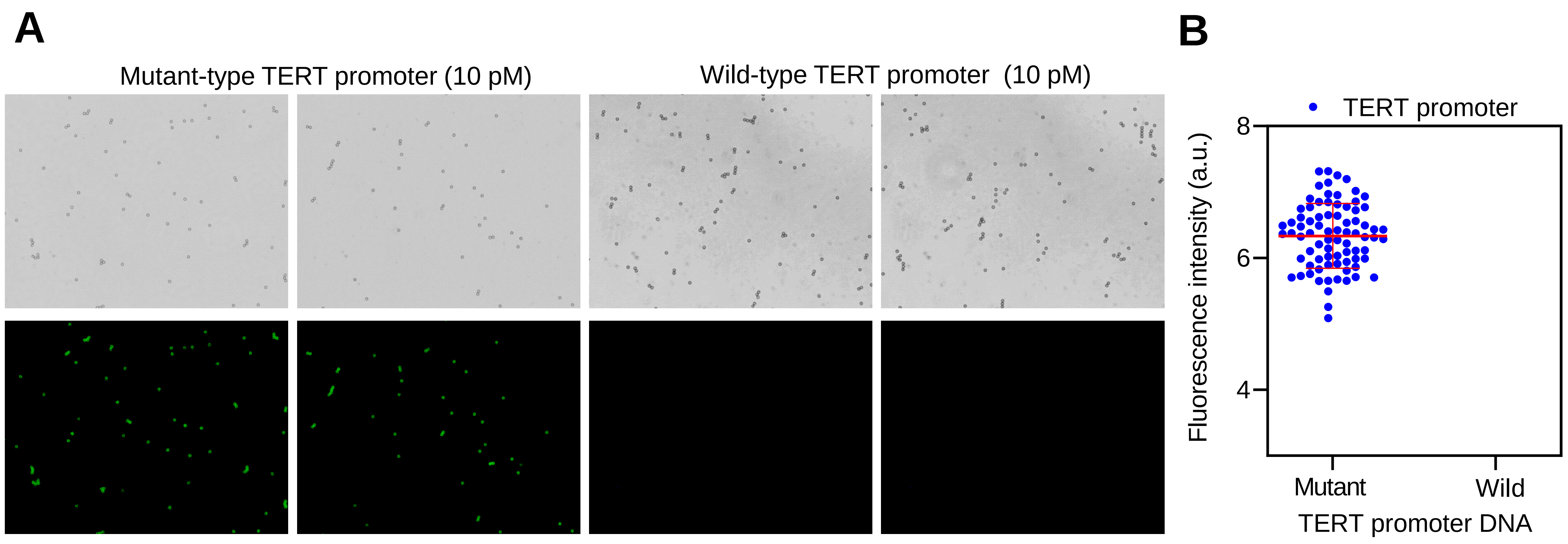


**Fig. S5**. Detection of TERT promoter DNA with Beads-HCR CADD. The wild-type (Wild) and mutant TERT promoter fragments were detected with sgRNAtert. (**A**) Beads fluorescence image. (**B**) Quantitative analysis of Beads fluorescence.

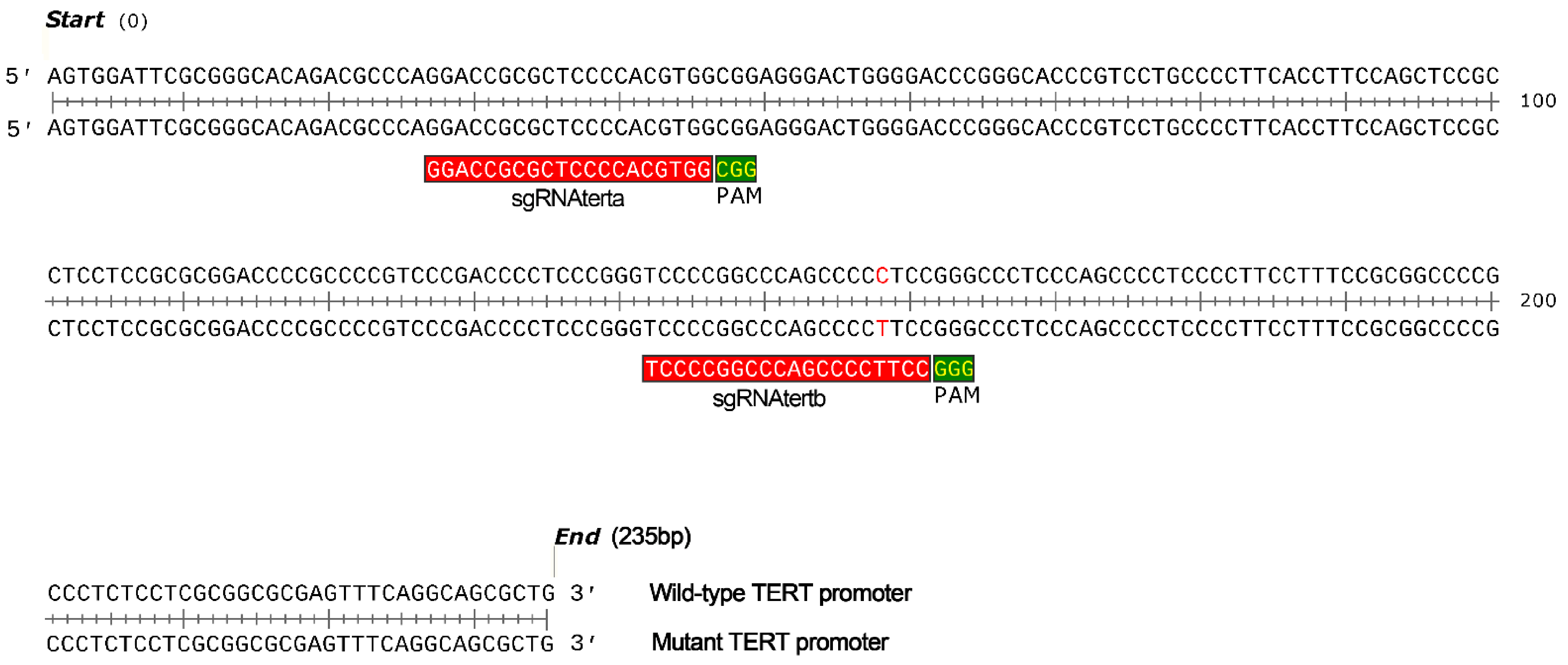


**Fig. S6**. DNA sequences of wild-type and mutant TERT promoter. The position of two sgRNAs targeting TERT promoter are shown. The sgRNAtertb was designed to detect the mutant TERT promoter.

### File 1

### Characterization of specificity of sgRNAs for 15 high-risk HPV subtypes

Detection of each of 15 high-risk HPV subtypes with all sgRNAs separately and together

pMD-HPV16–73: pMD contains L1 fragment of HPV16~73
pMD: pMD contains no L1 fragment (a blank plasmid for cloning L1 fragment )
sgRNA16–73: sgRNA specific to L1 fragment of HPV16~73 (sgRNAsp)
sgRNAct: cocktail sgRNA (an equimolar mixture of sgRNA16–73)

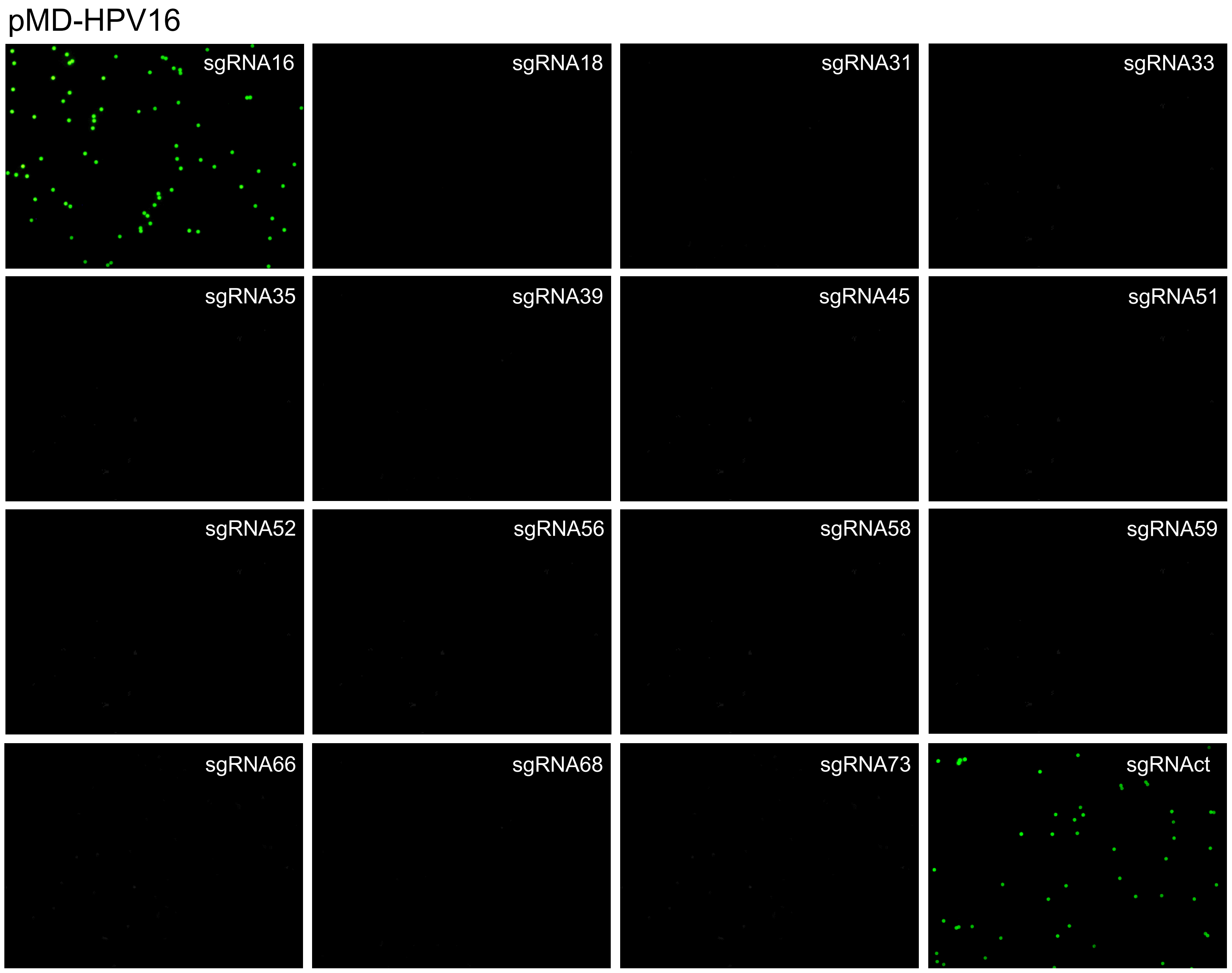
pMD-HPV16
sgRNA16
sgRNA18
sgRNA31
sgRNA33
sgRNA35
sgRNA39
sgRNA45
sgRNA51
sgRNA52
sgRNA56
sgRNA58
sgRNA59
sgRNA66
sgRNA68
sgRNA73
sgRNAct

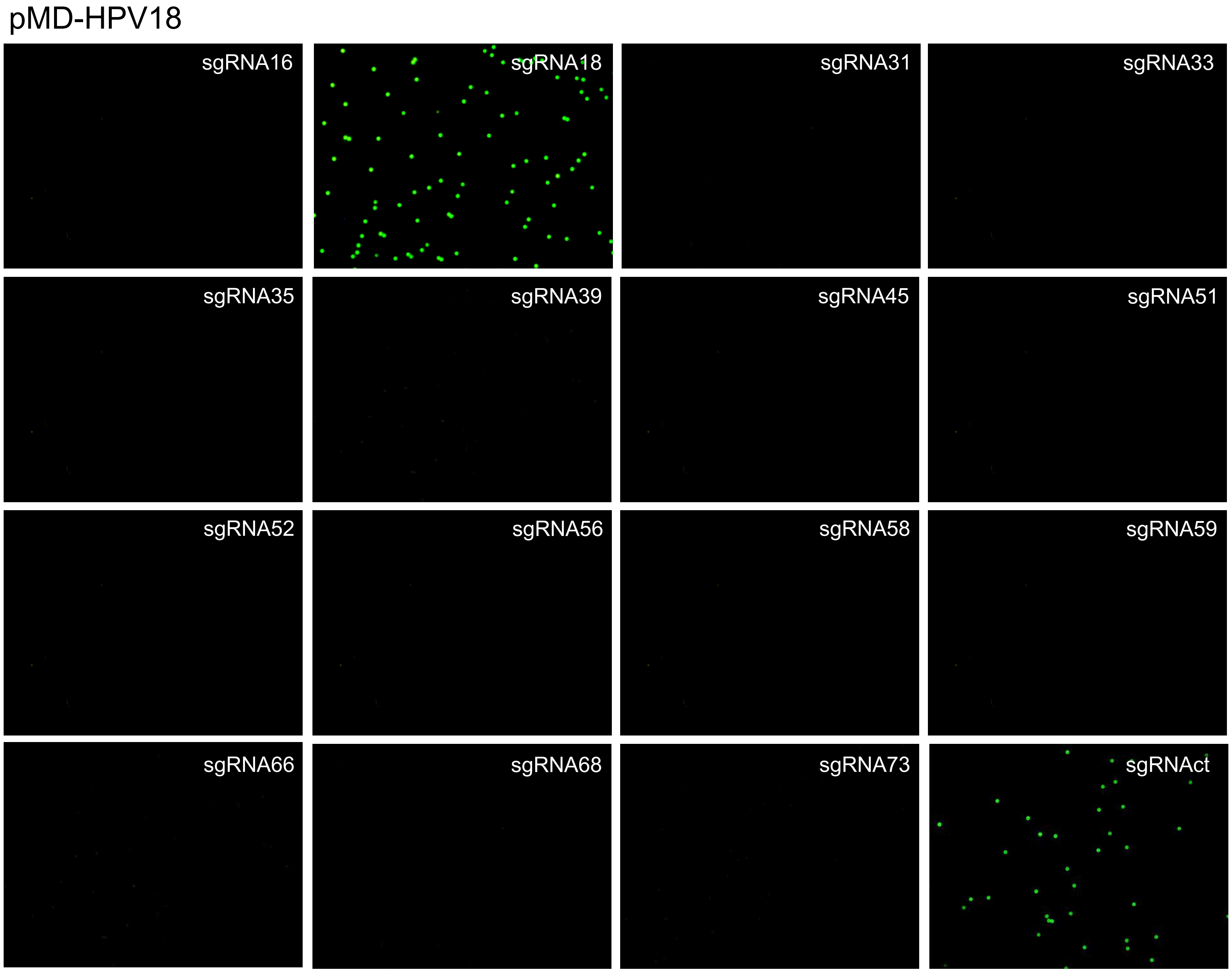
pMD-HPV18
sgRNA16
sgRNA18
sgRNA31
sgRNA33
sgRNA35
sgRNA39
sgRNA45
sgRNA51
sgRNA52
sgRNA56
sgRNA58
sgRNA59
sgRNA66
sgRNA68
sgRNA73
sgRNAct

pMD-HPV31

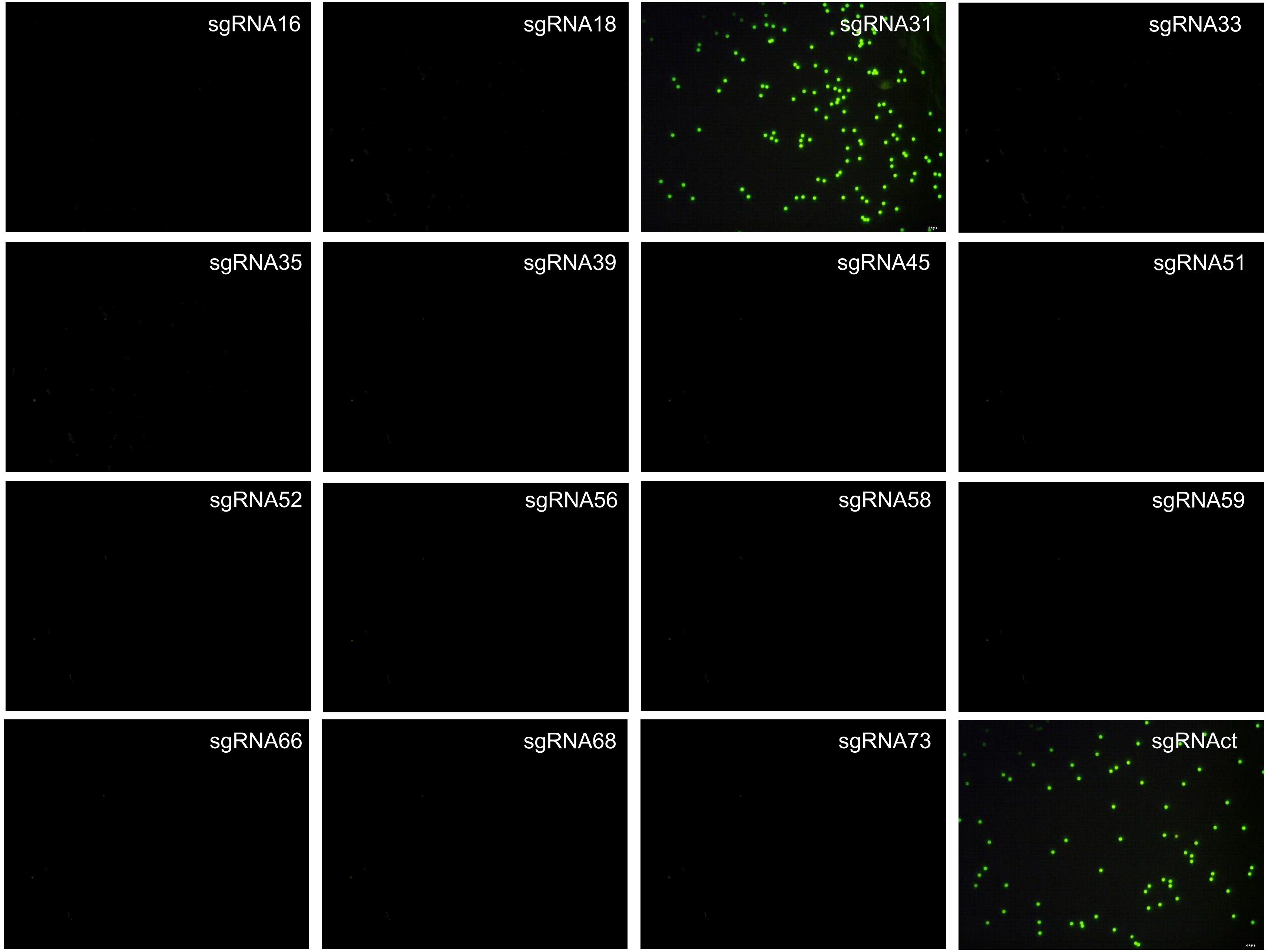
sgRNA16
sgRNA18
sgRNA31
sgRNA33
sgRNA35
sgRNA39
sgRNA45
sgRNA51
sgRNA52
sgRNA56
sgRNA58
sgRNA59
sgRNA66
sgRNA68
sgRNA73
sgRNAct

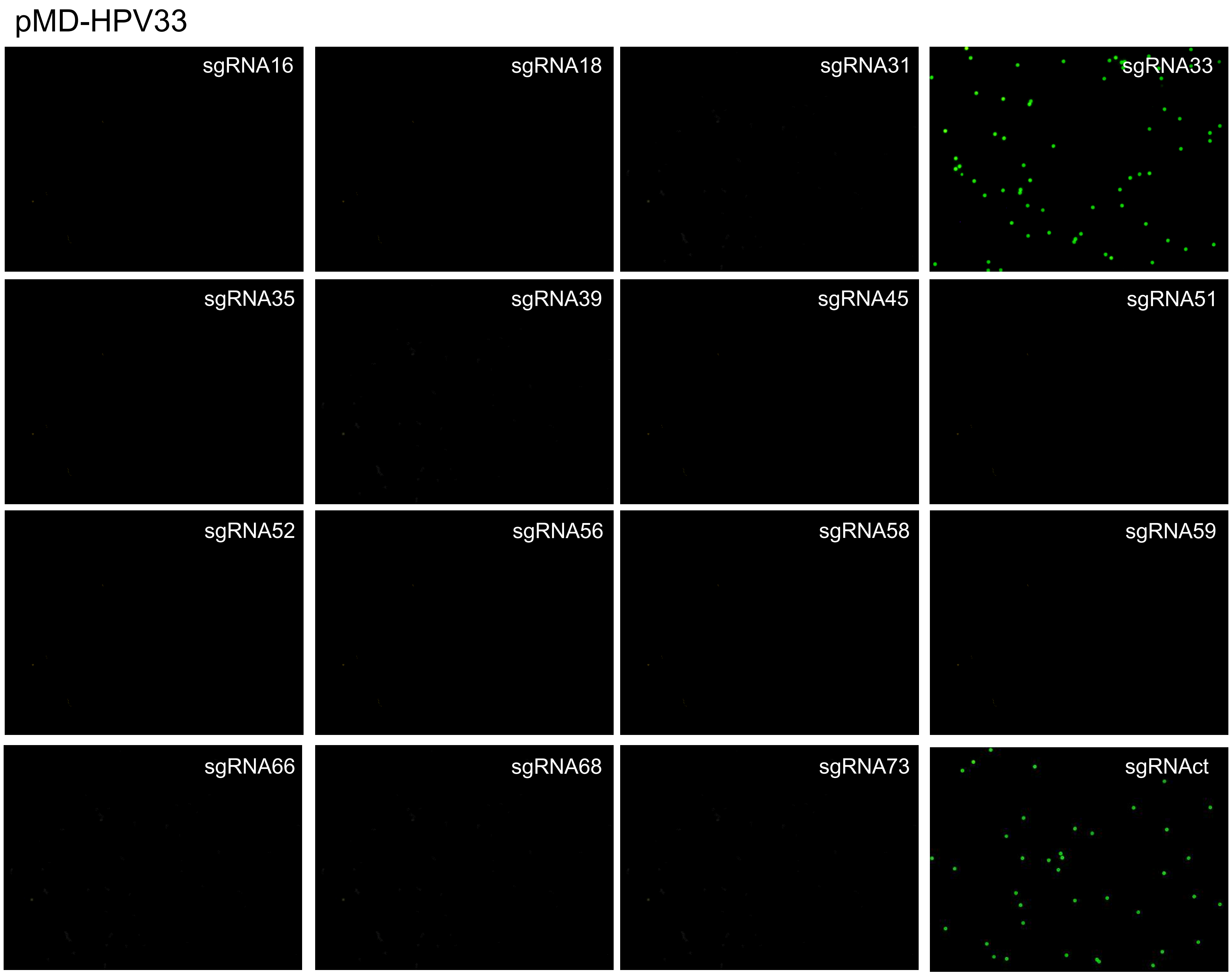
pMD-HPV33
sgRNA16
sgRNA18
sgRNA31
sgRNA33
sgRNA35
sgRNA39
sgRNA45
sgRNA51
sgRNA52
sgRNA56
sgRNA58
sgRNA59
sgRNA66
sgRNA68
sgRNA73
sgRNAct

pMD-HPV35

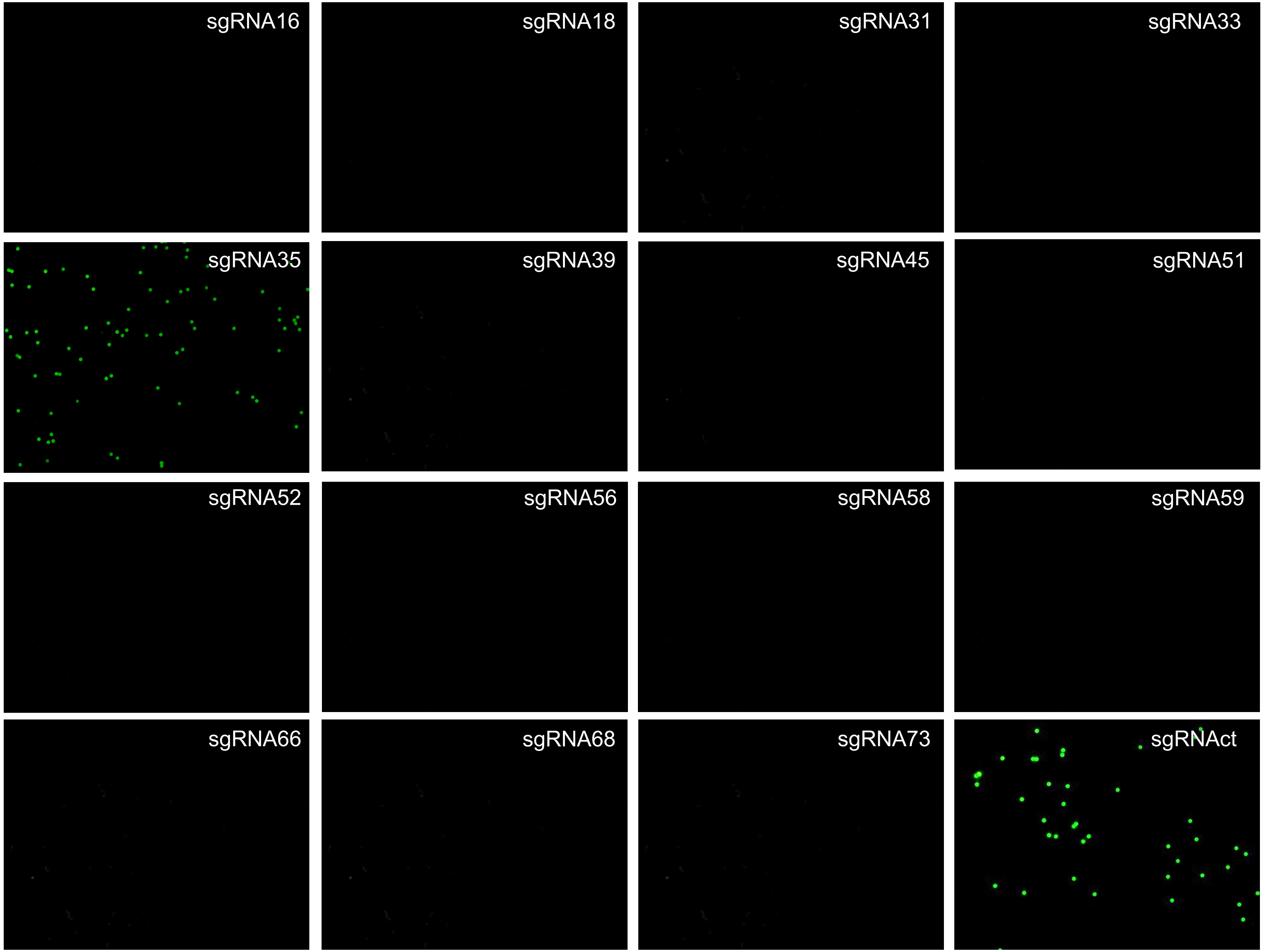
sgRNA16
sgRNA18
sgRNA31
sgRNA33
sgRNA35
sgRNA39
sgRNA45
sgRNA51
sgRNA52
sgRNA56
sgRNA58
sgRNA59
sgRNA66
sgRNA68
sgRNA73
sgRNAct

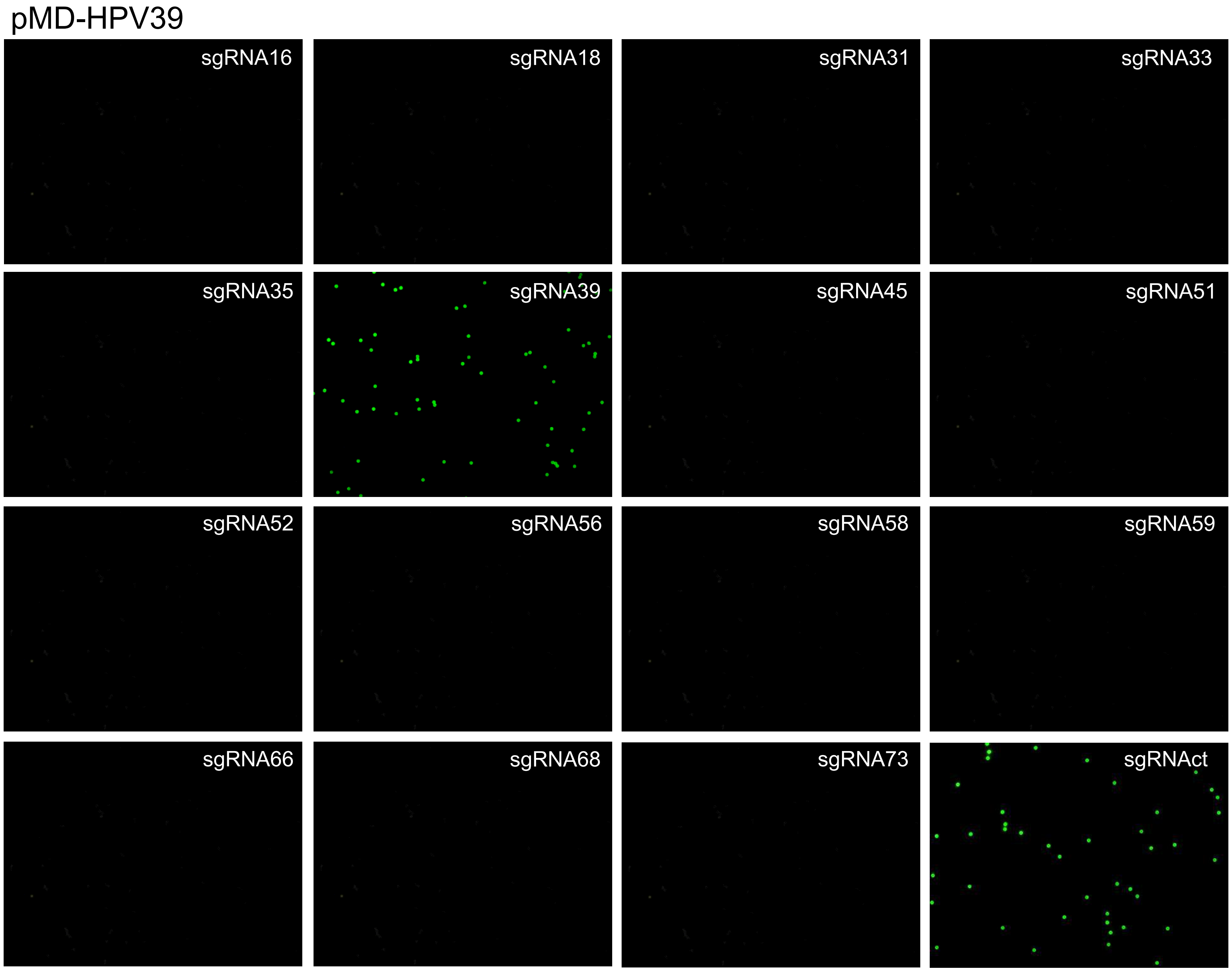
pMD-HPV39
sgRNA16
sgRNA18
sgRNA31
sgRNA33
sgRNA35
sgRNA39
sgRNA45
sgRNA51
sgRNA52
sgRNA56
sgRNA58
sgRNA59
sgRNA66
sgRNA68
sgRNA73
sgRNAct

pMD-HPV45

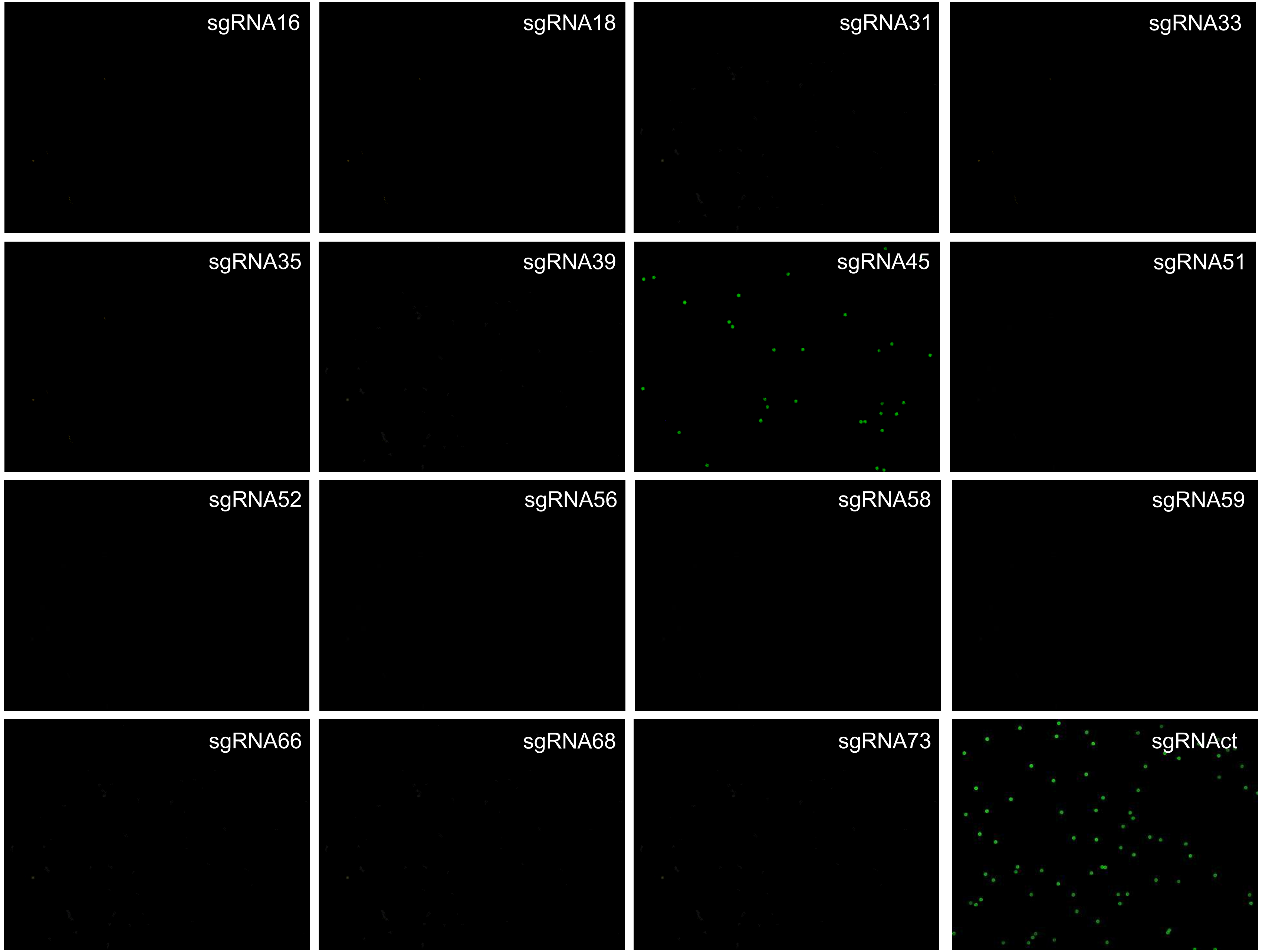
sgRNA16
sgRNA18
sgRNA31
sgRNA33
sgRNA35
sgRNA39
sgRNA45
sgRNA51
sgRNA52
sgRNA56
sgRNA58
sgRNA59
sgRNA66
sgRNA68
sgRNA73
sgRNAct

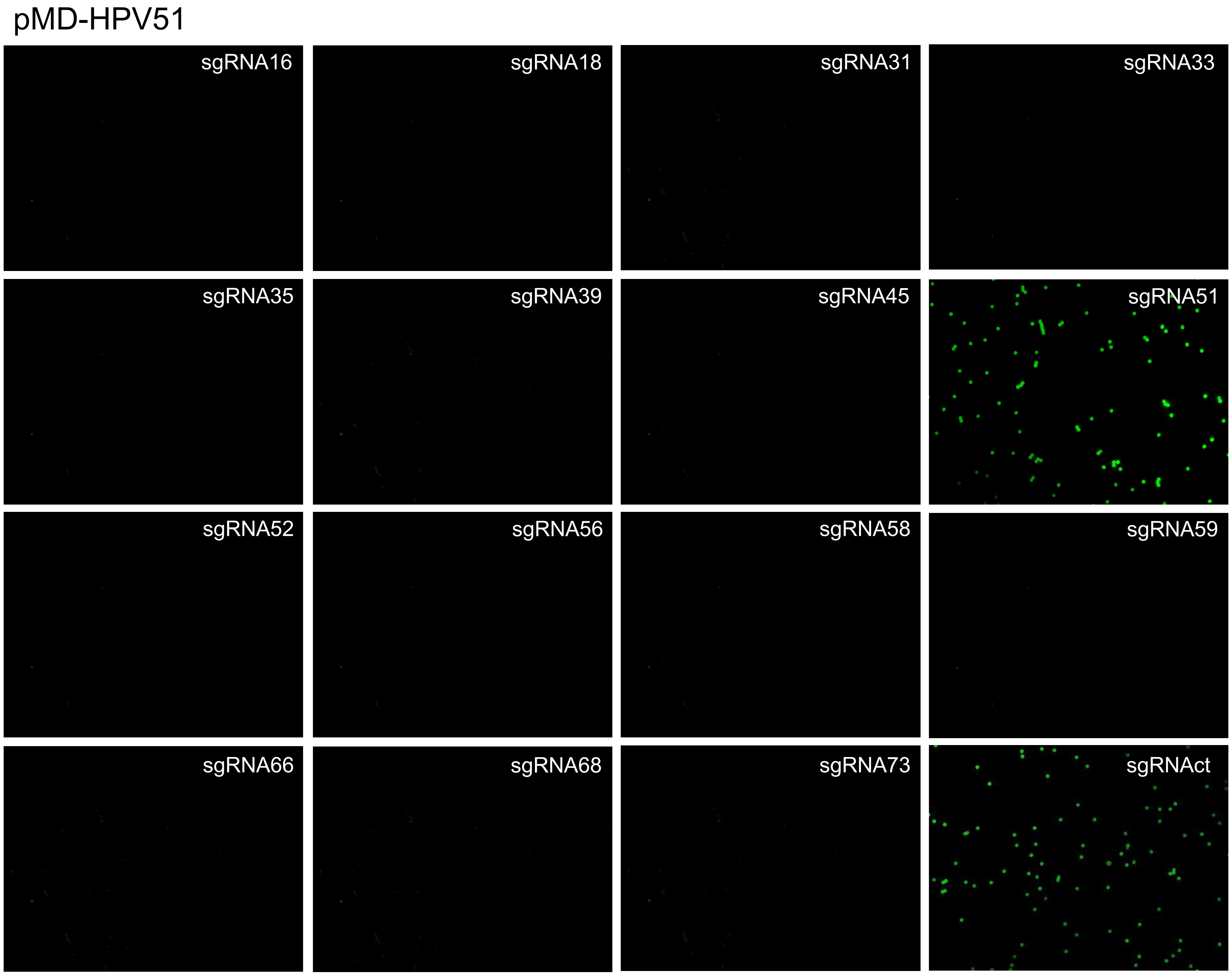
pMD-HPV51
sgRNA16
sgRNA18
sgRNA31
sgRNA33
sgRNA35
sgRNA39
sgRNA45
sgRNA51
sgRNA52
sgRNA56
sgRNA58
sgRNA59
sgRNA66
sgRNA68
sgRNA73
sgRNAct

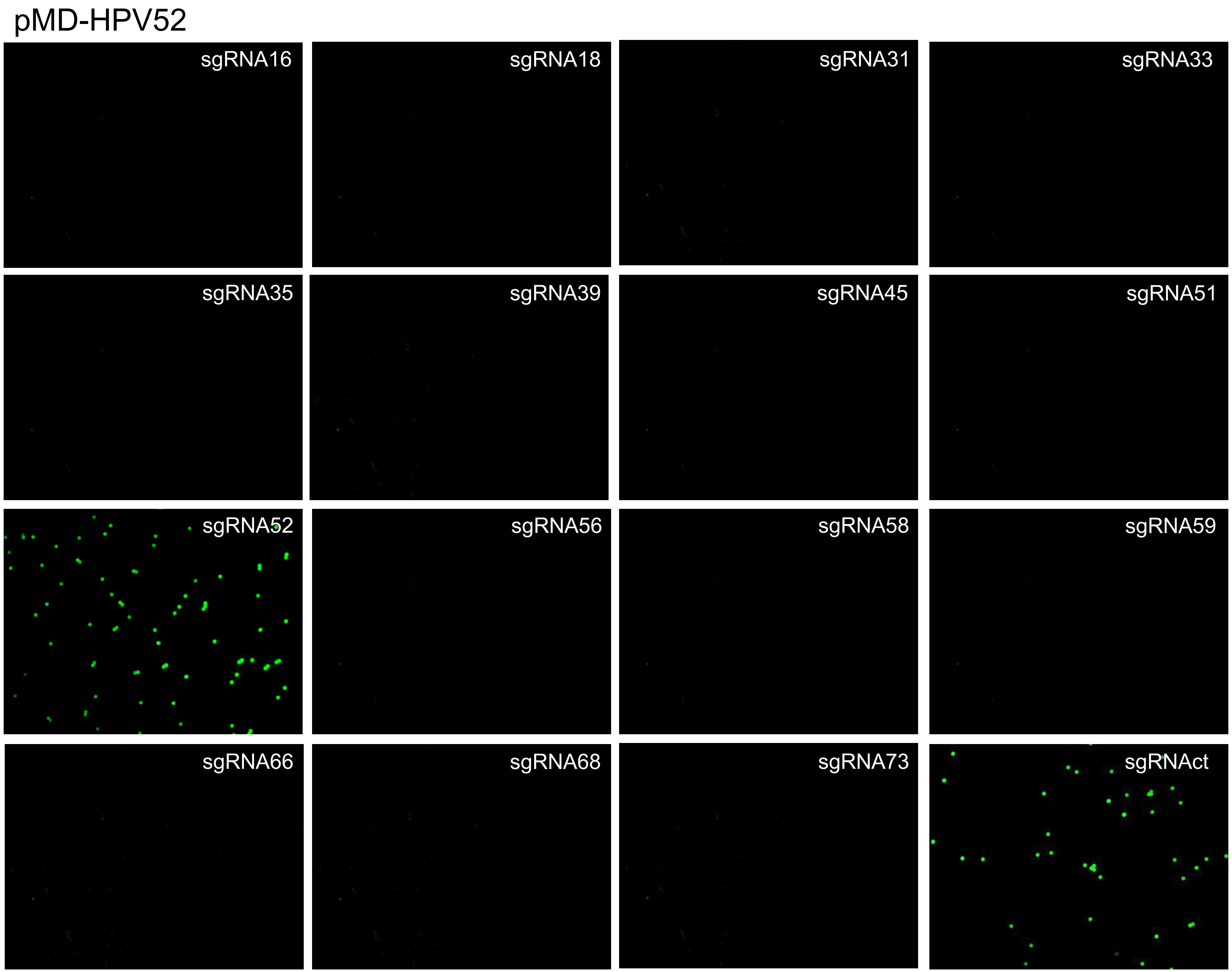
pMD-HPV52
sgRNA16
sgRNA18
sgRNA31
sgRNA33
sgRNA35
sgRNA39
sgRNA45
sgRNA51
sgRNA52
sgRNA56
sgRNA58
sgRNA59
sgRNA66
sgRNA68
sgRNA73
sgRNAct

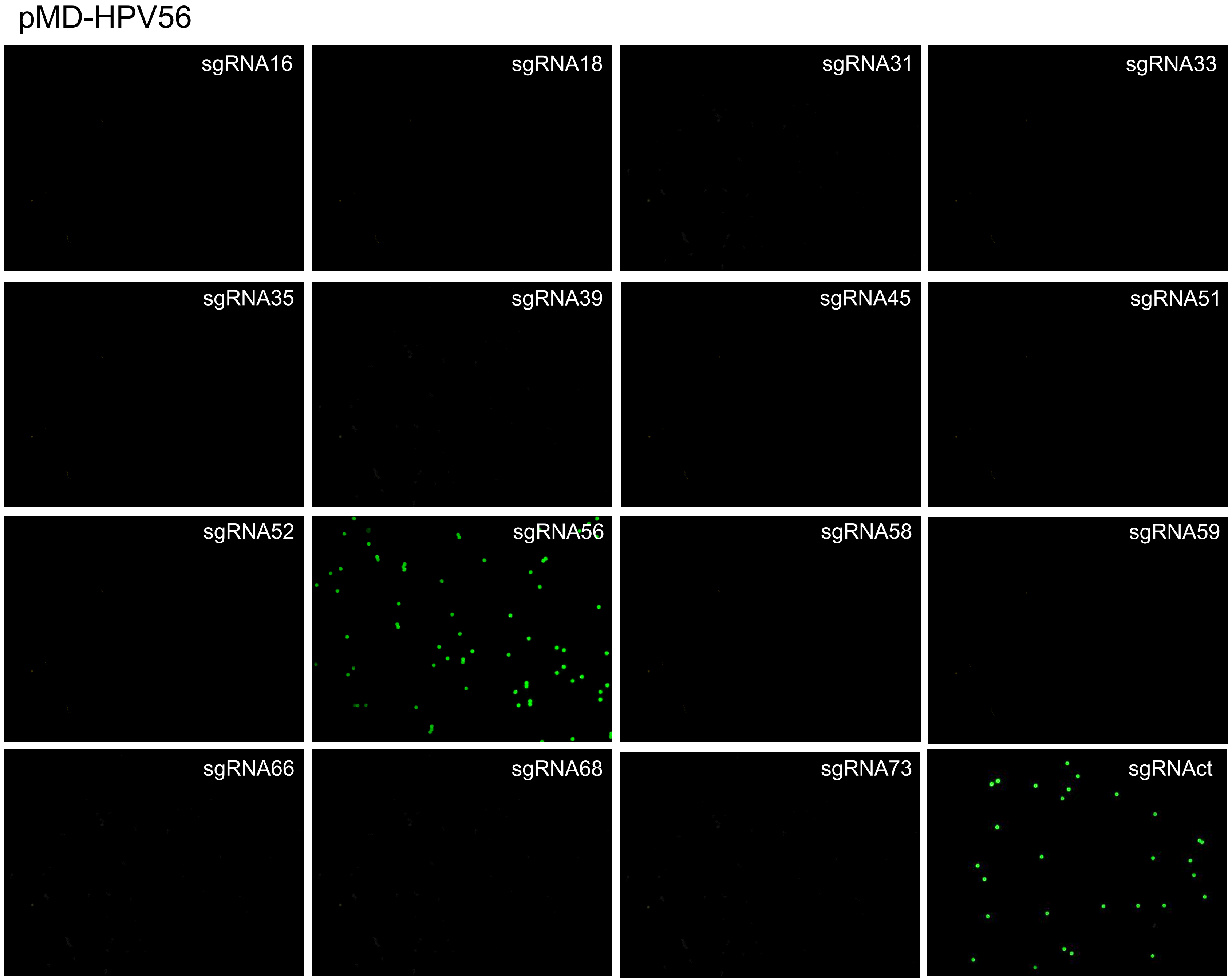
pMD-HPV56
sgRNA16
sgRNA18
sgRNA31
sgRNA33
sgRNA35
sgRNA39
sgRNA45
sgRNA51
sgRNA52
sgRNA56
sgRNA58
sgRNA59
sgRNA66
sgRNA68
sgRNA73
sgRNAct

pMD-HPV58

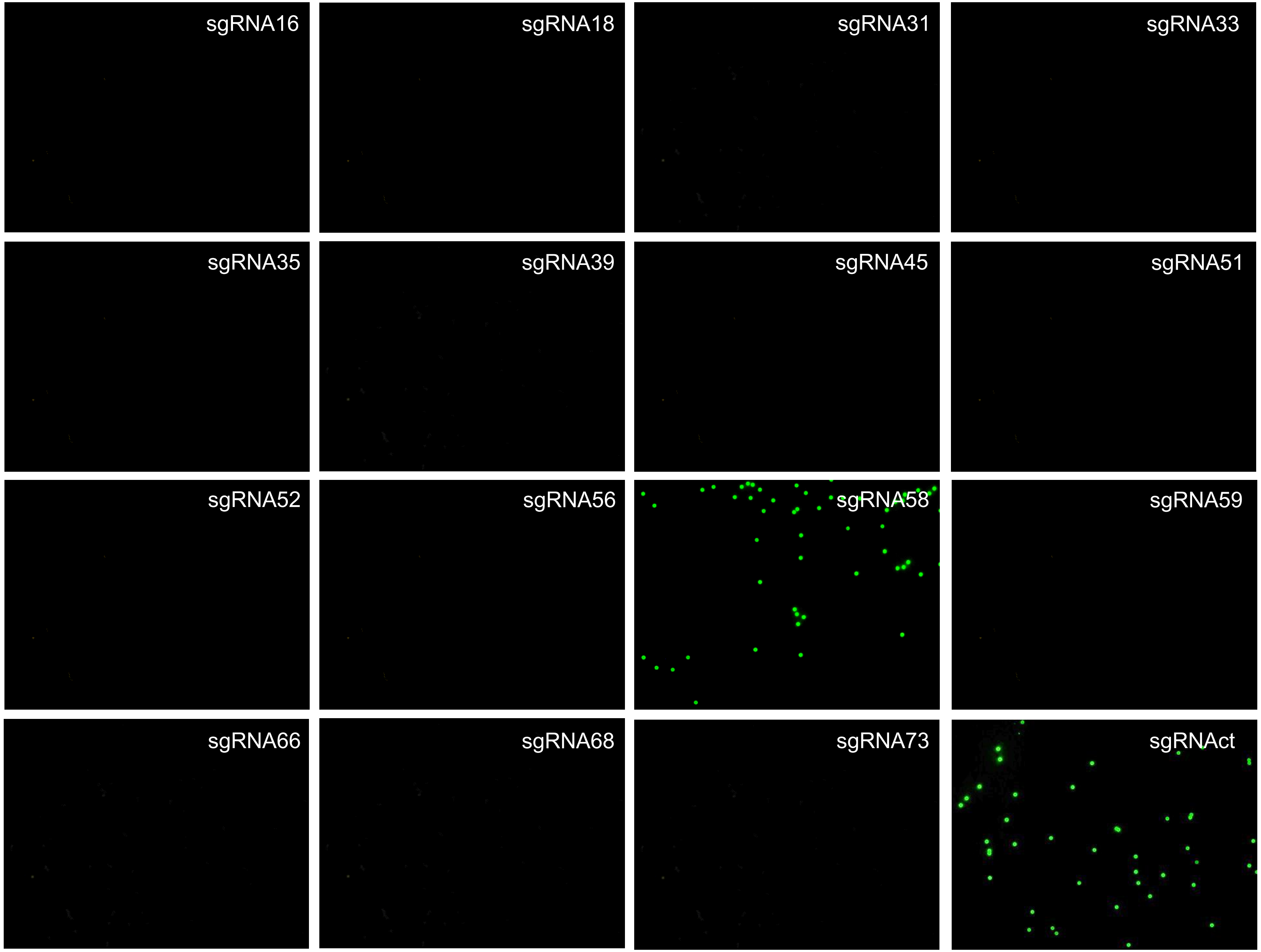
sgRNA16
sgRNA18
sgRNA31
sgRNA33
sgRNA35
sgRNA39
sgRNA45
sgRNA51
sgRNA52
sgRNA56
sgRNA58
sgRNA59
sgRNA66
sgRNA68
sgRNA73
sgRNAct

pMD-HPV59

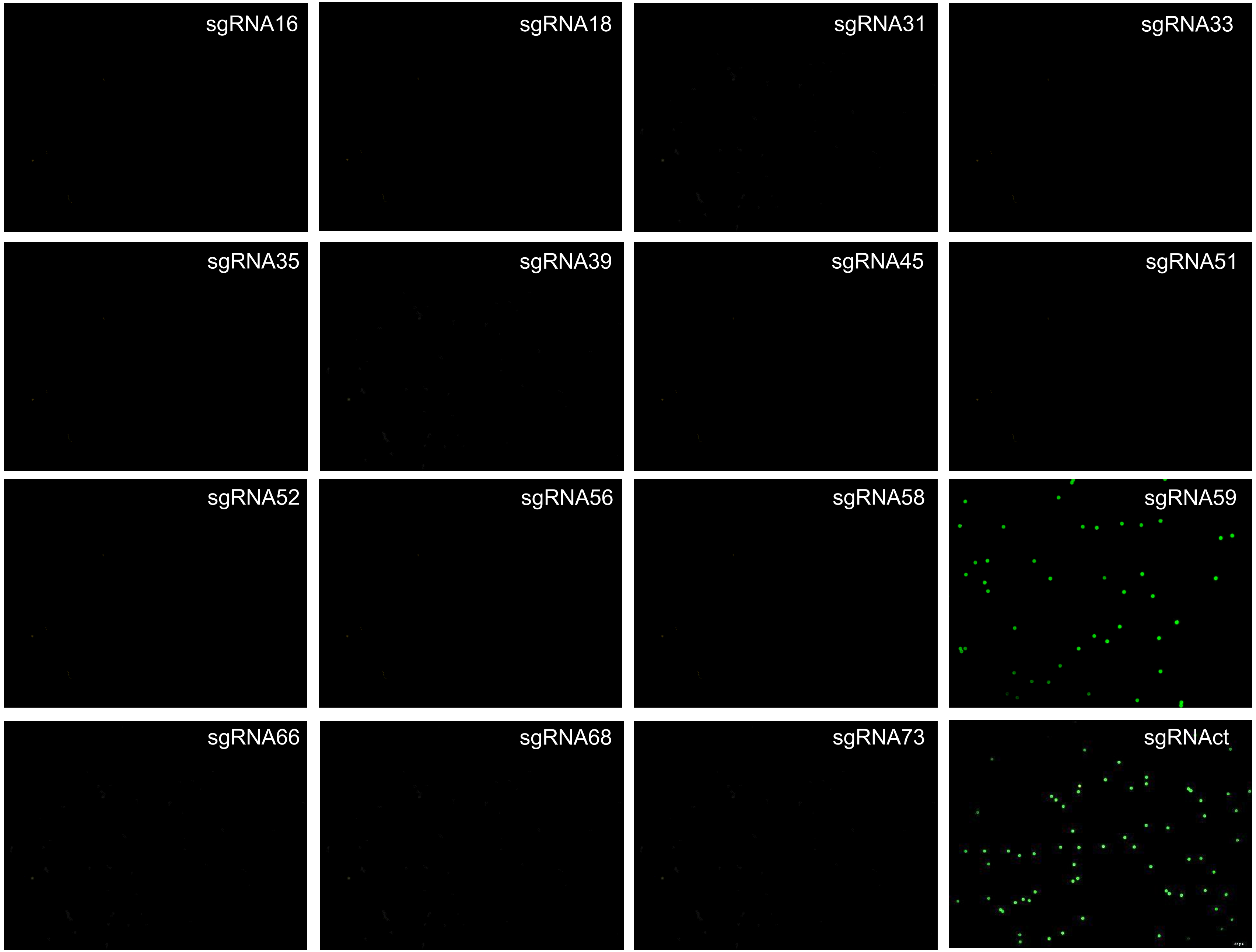
sgRNA16
sgRNA18
sgRNA31
sgRNA33
sgRNA35
sgRNA39
sgRNA45
sgRNA51
sgRNA52
sgRNA56
sgRNA58
sgRNA59
sgRNA66
sgRNA68
sgRNA73
sgRNAct

pMD-HPV66

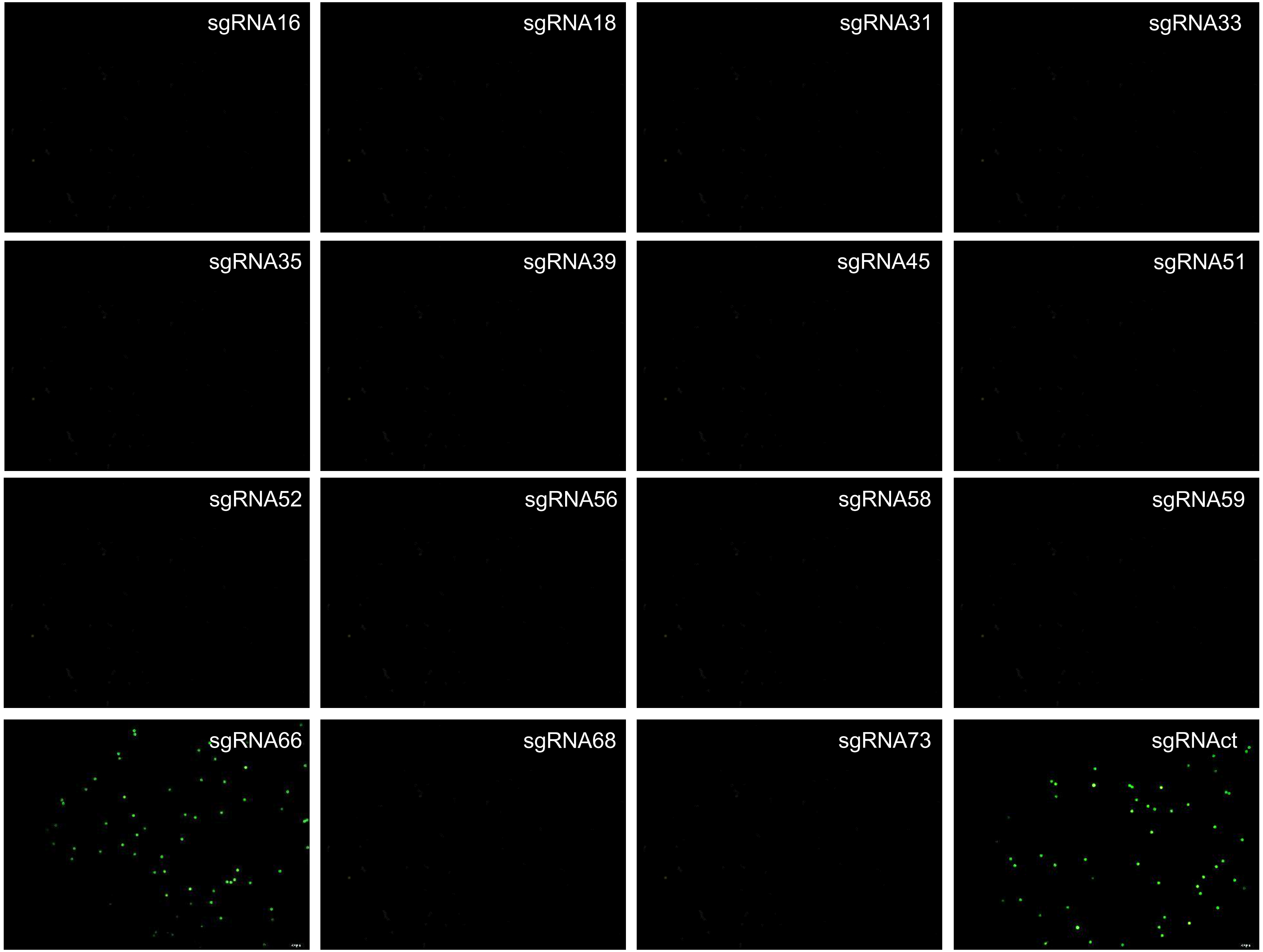
sgRNA16
sgRNA18
sgRNA31
sgRNA33
sgRNA35
sgRNA39
sgRNA45
sgRNA51
sgRNA52
sgRNA56
sgRNA58
sgRNA59
sgRNA66
sgRNA68
sgRNA73
sgRNAct

pMD-HPV68

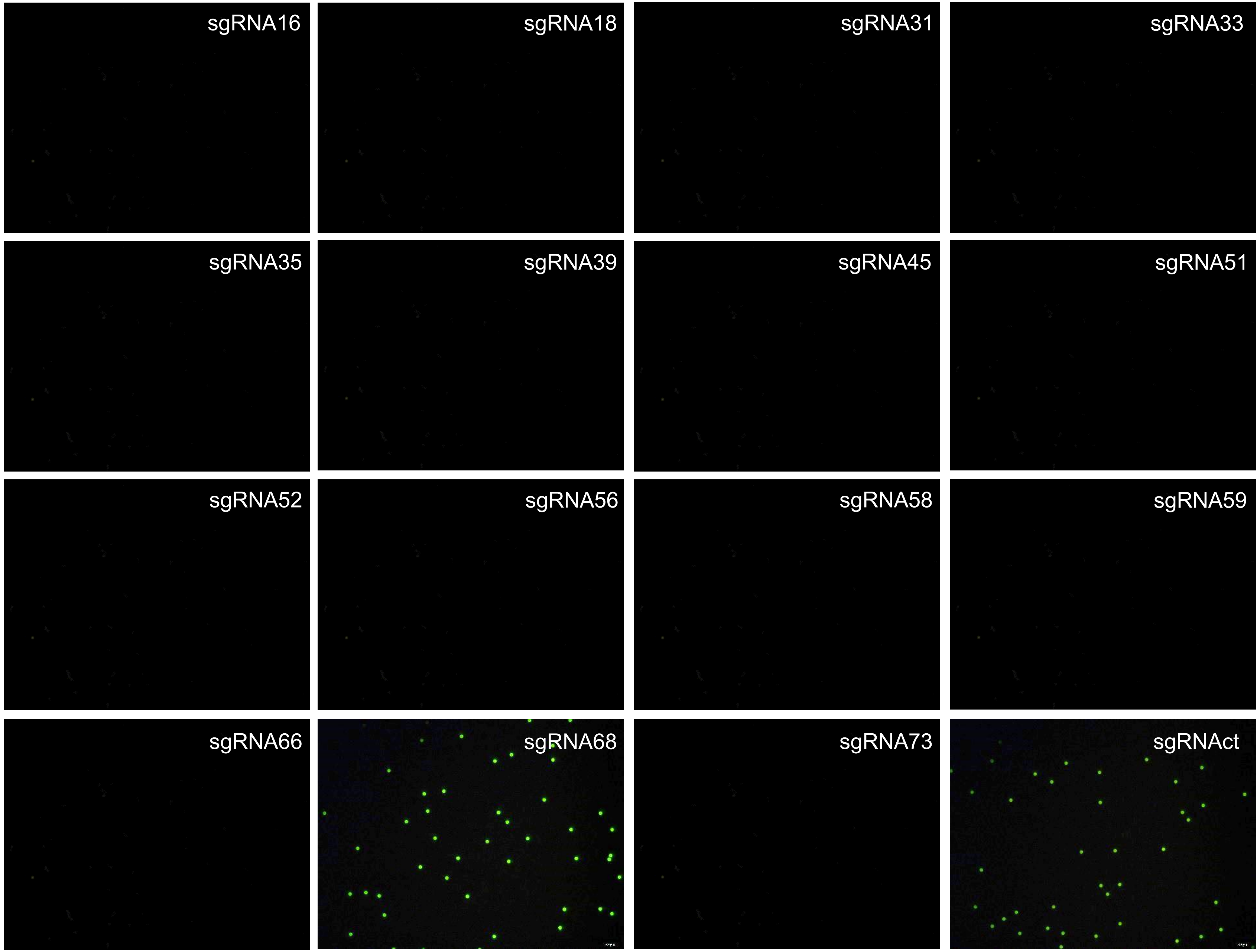
sgRNA16
sgRNA18
sgRNA31
sgRNA33
sgRNA35
sgRNA39
sgRNA45
sgRNA51
sgRNA52
sgRNA56
sgRNA58
sgRNA59
sgRNA66
sgRNA68
sgRNA73
sgRNAct

pMD-HPV73

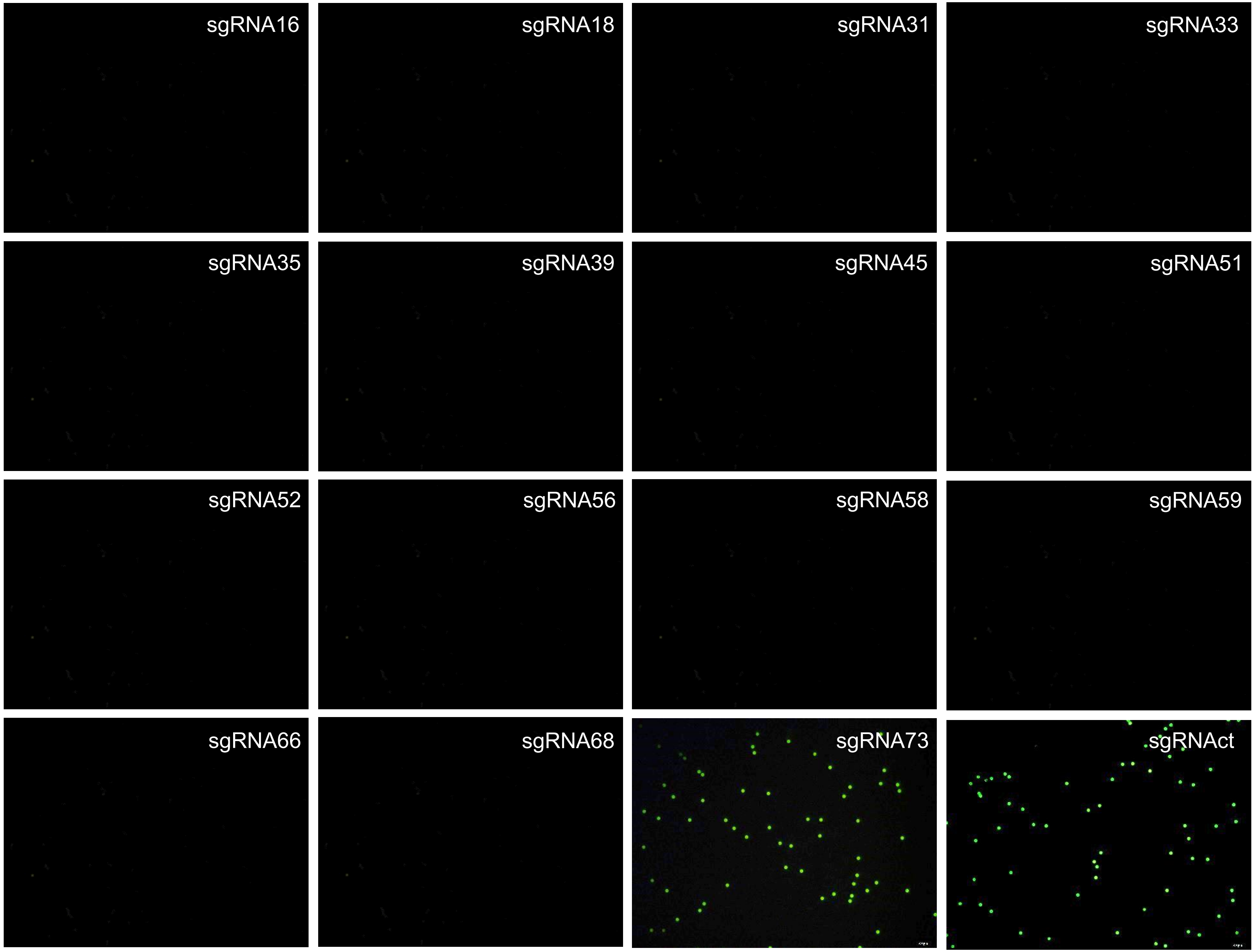
sgRNA16
sgRNA18
sgRNA31
sgRNA33
sgRNA35
sgRNA39
sgRNA45
sgRNA51
sgRNA52
sgRNA56
sgRNA58
sgRNA59
sgRNA66
sgRNA68
sgRNA73
sgRNAct

pMD

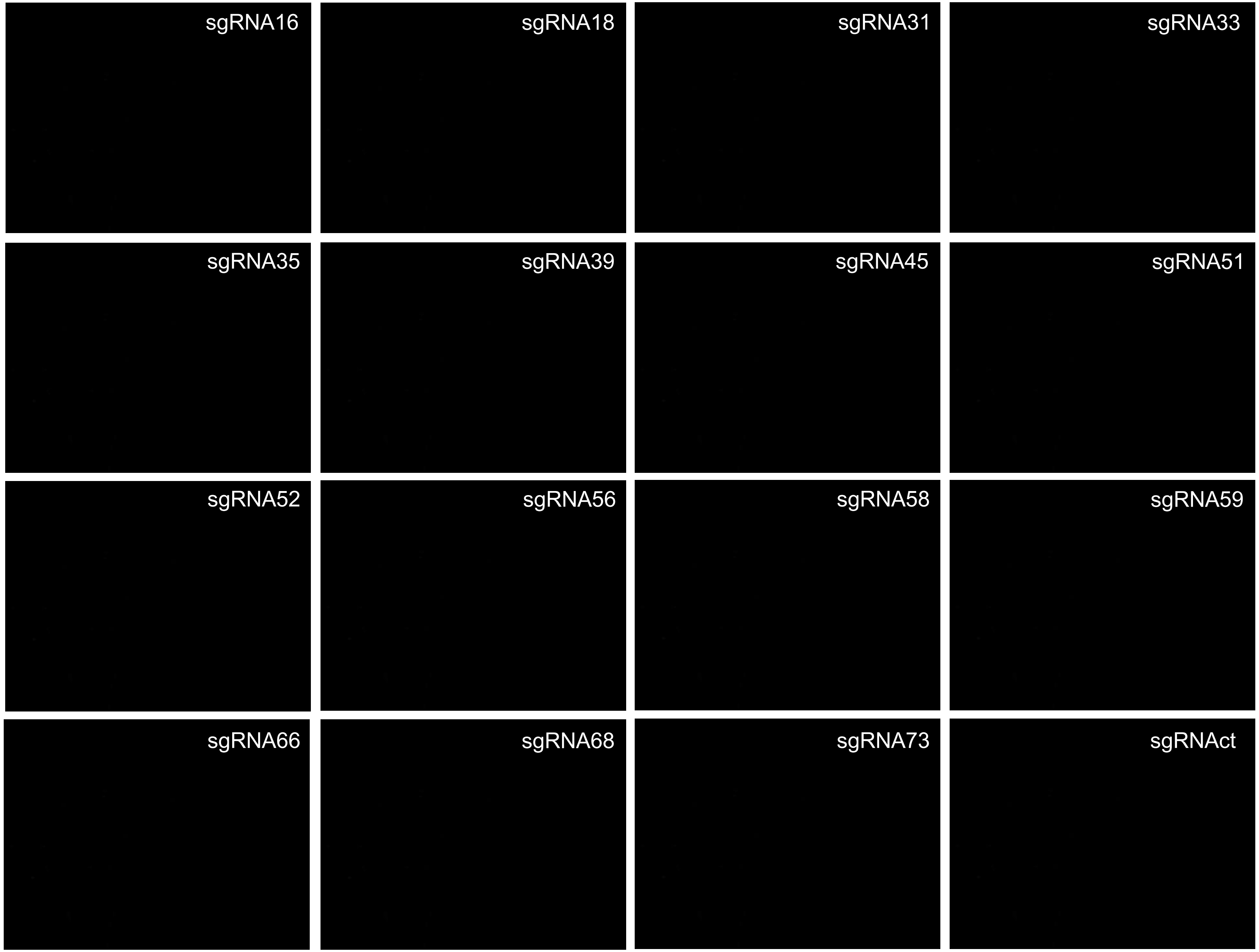
sgRNA16
sgRNA18
sgRNA31
sgRNA33
sgRNA35
sgRNA39
sgRNA45
sgRNA51
sgRNA52
sgRNA56
sgRNA58
sgRNA59
sgRNA66
sgRNA68
sgRNA73
sgRNAct

### File 2

### Detection of clinical samples with Beads-HCR-based CADD

sgRNA16–73: sgRNA specific to L1 fragment of HPV16~73  (sgRNAsp)
sgRNAct: cocktail sgRNA (an equimolar mixture of sgRNA16–73)

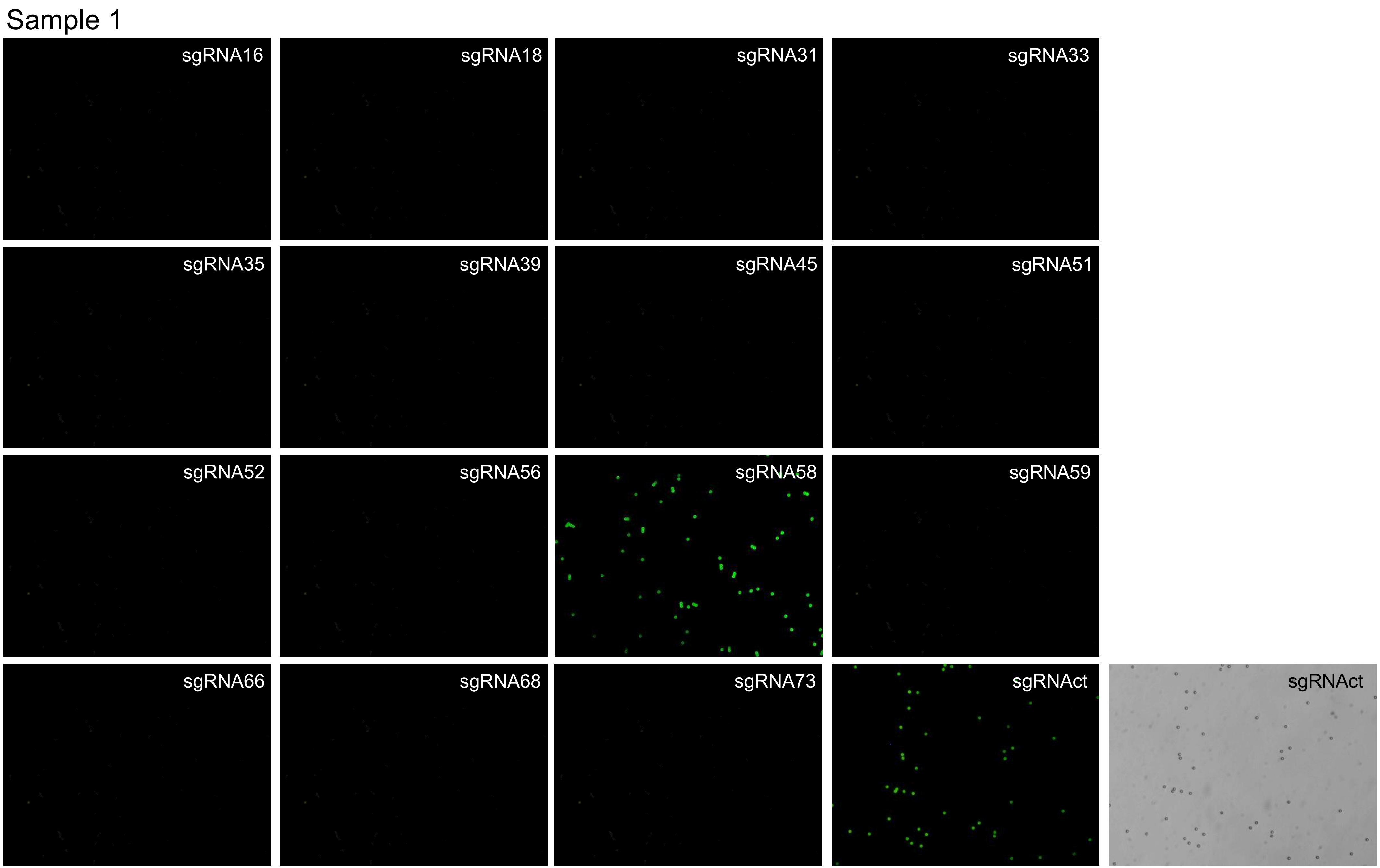
Sample 1
sgRNA16
sgRNA18
sgRNA31
sgRNA33
sgRNA35
sgRNA39
sgRNA45
sgRNA51
sgRNA52
sgRNA56
sgRNA58
sgRNA59
sgRNA66
sgRNA68
sgRNA73
sgRNAct
sgRNAct

Sample 2

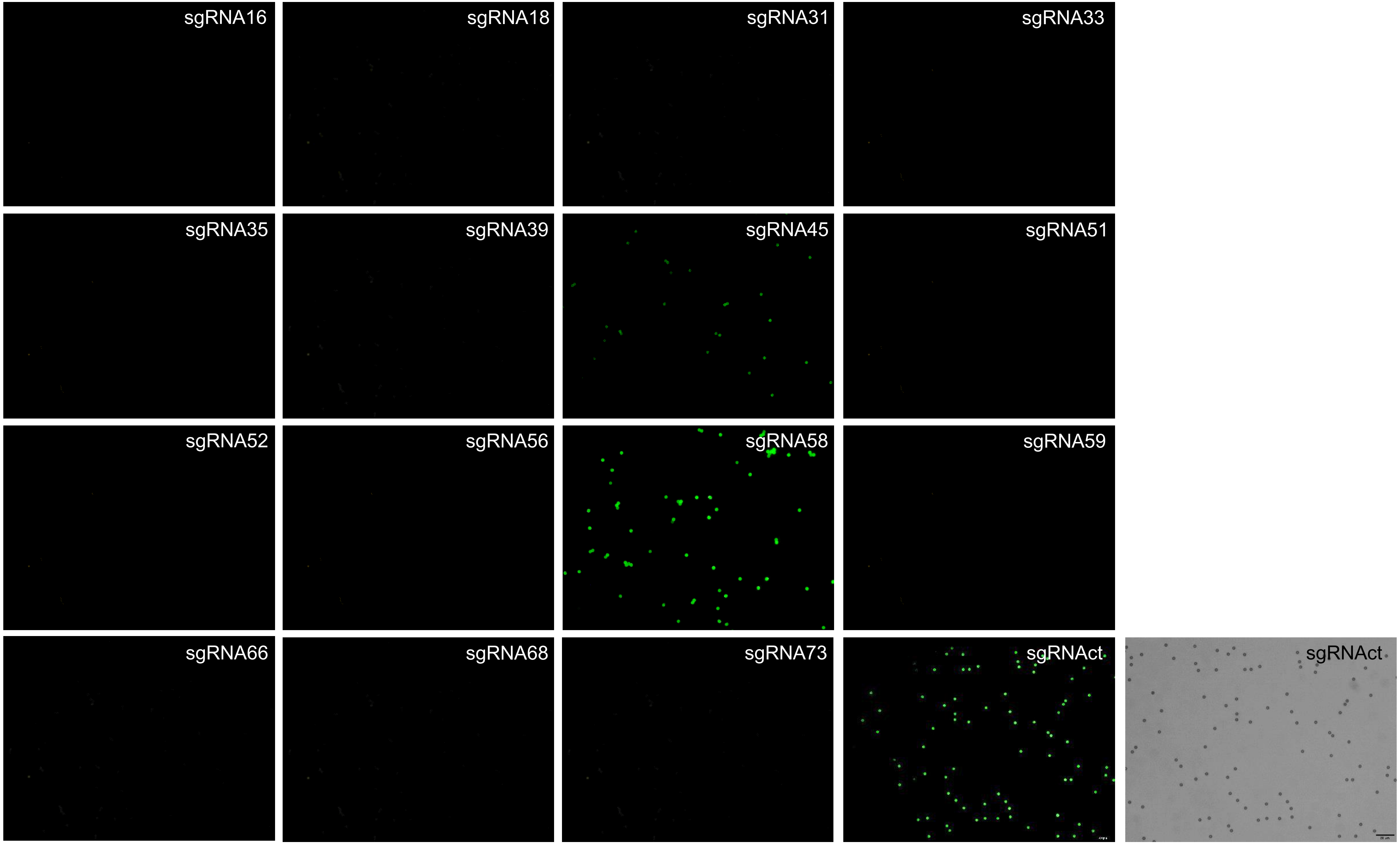

sgRNA16
sgRNA18
sgRNA31
sgRNA33
sgRNA35
sgRNA39
sgRNA45
sgRNA51
sgRNA52
sgRNA56
sgRNA58
sgRNA59
sgRNA66
sgRNA68
sgRNA73
sgRNAct
sgRNAct

Sample 3

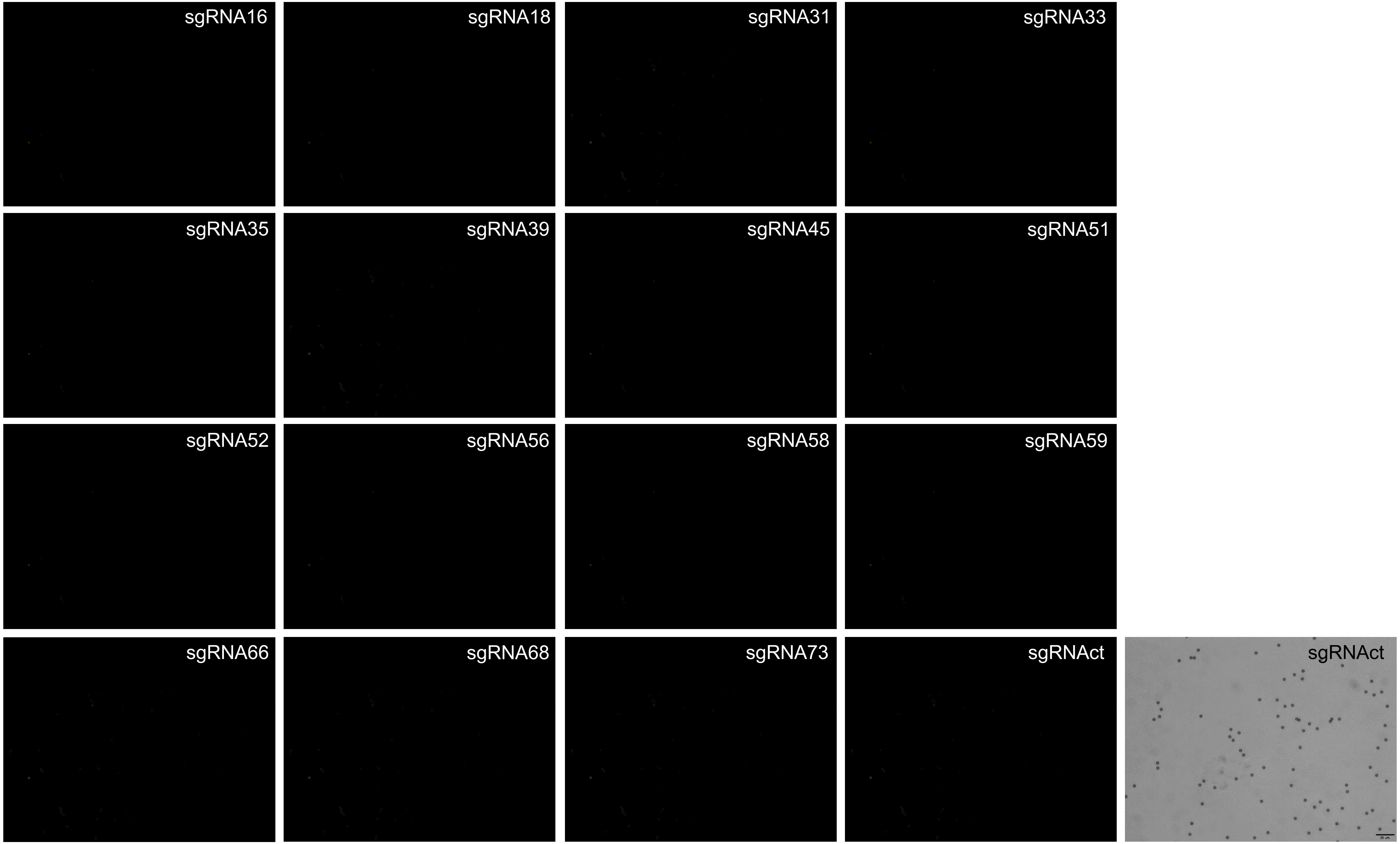
sgRNA16
sgRNA18
sgRNA31
sgRNA33
sgRNA35
sgRNA39
sgRNA45
sgRNA51
sgRNA52
sgRNA56
sgRNA58
sgRNA59
sgRNA66
sgRNA68
sgRNA73
sgRNAct
sgRNAct

Sample 4

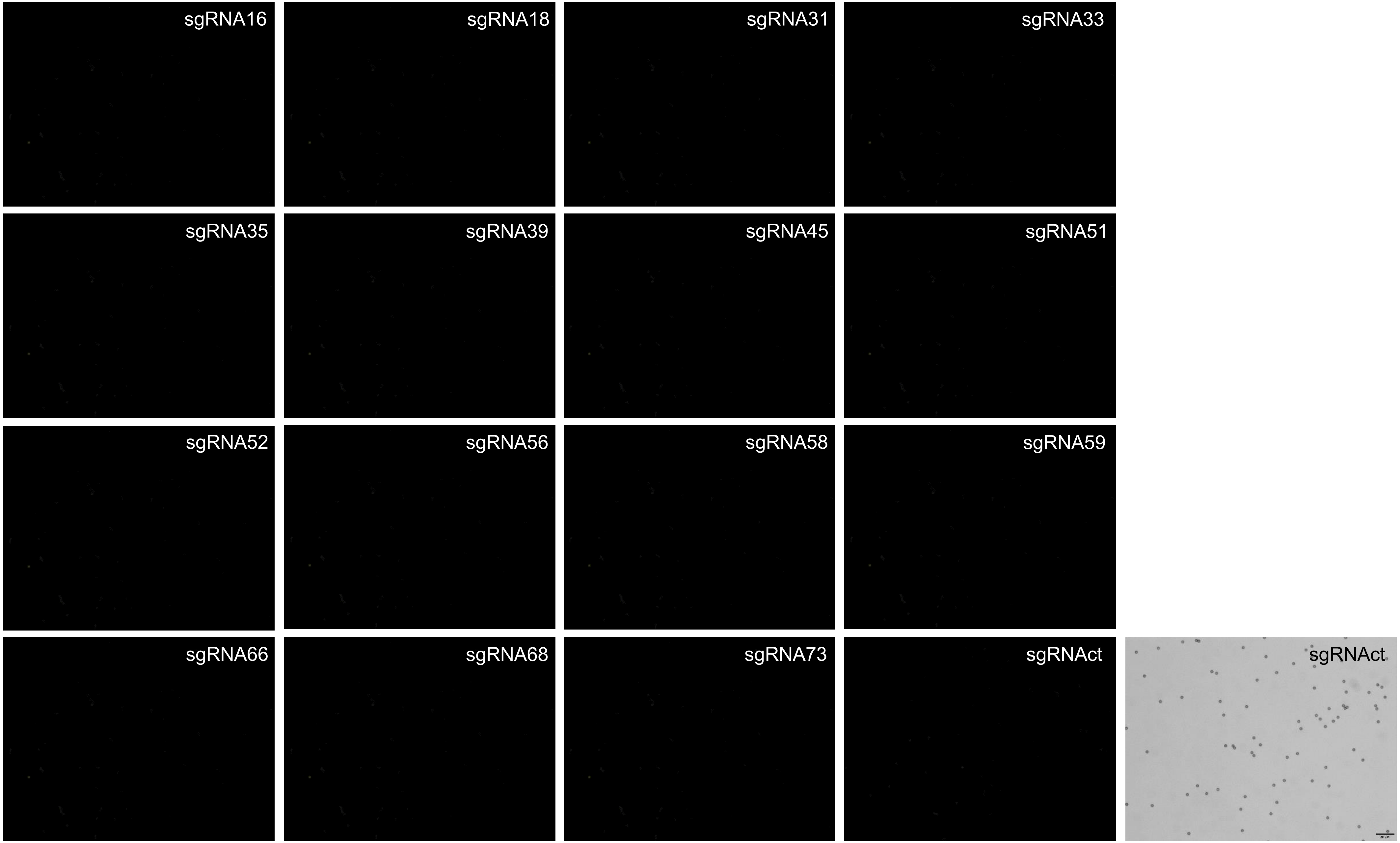
sgRNA16
sgRNA18
sgRNA31
sgRNA33
sgRNA35
sgRNA39
sgRNA45
sgRNA51
sgRNA52
sgRNA56
sgRNA58
sgRNA59
sgRNA66
sgRNA68
sgRNA73
sgRNAct
sgRNAct

Sample 5

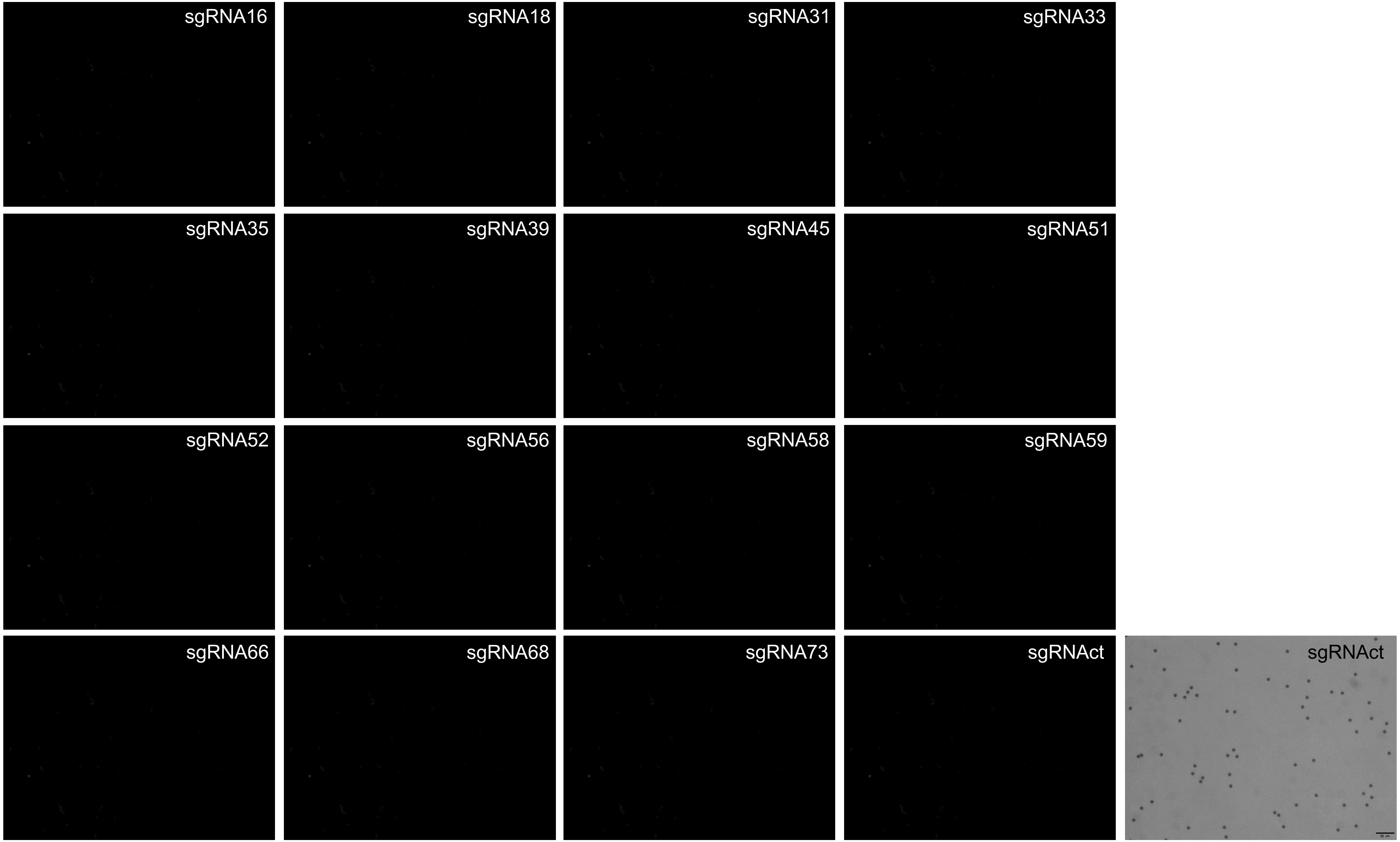
sgRNA16
sgRNA18
sgRNA31
sgRNA33
sgRNA35
sgRNA39
sgRNA45
sgRNA51
sgRNA52
sgRNA56
sgRNA58
sgRNA59
sgRNA66
sgRNA68
sgRNA73
sgRNAct
sgRNAct

Sample 6

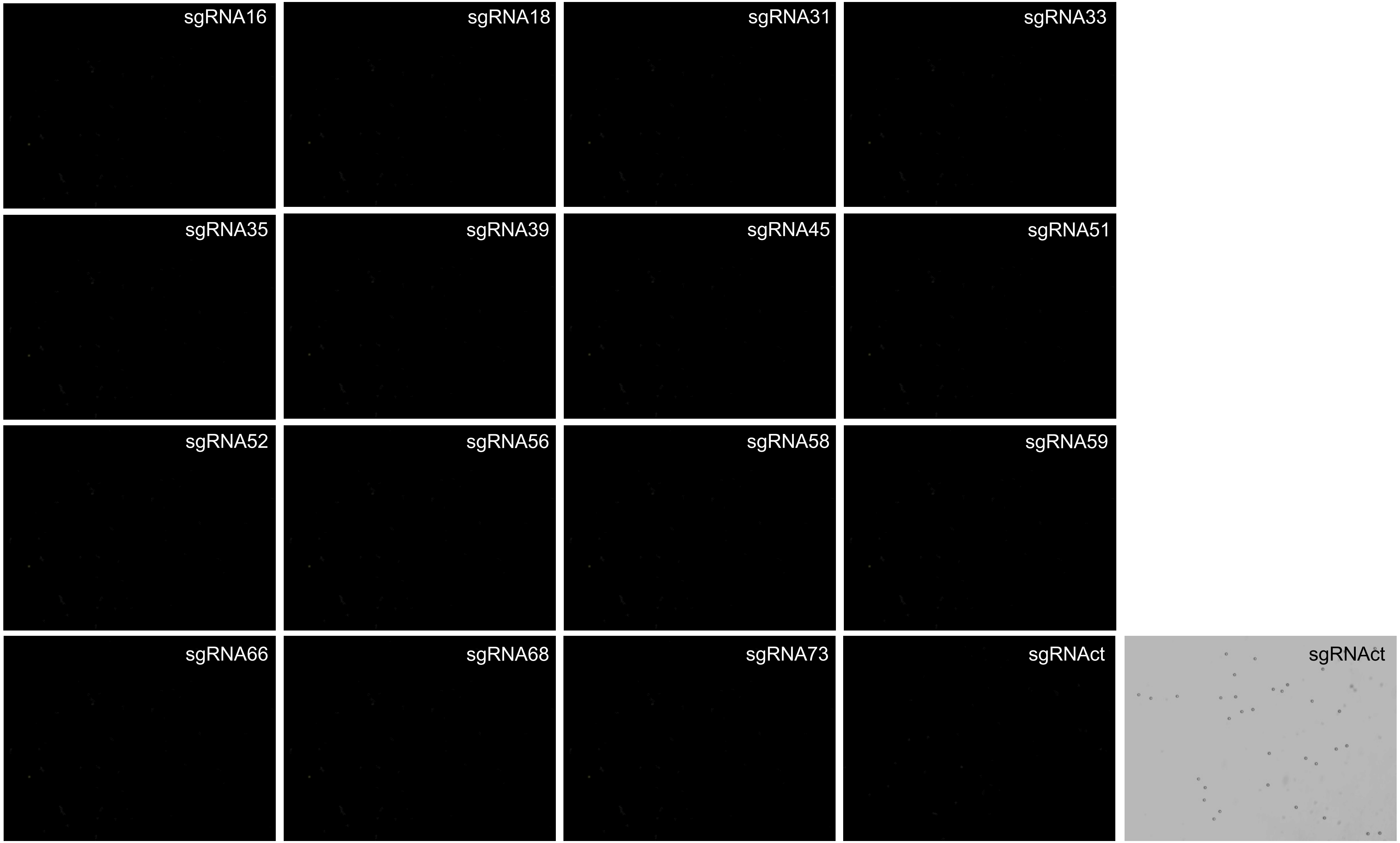
sgRNA16
sgRNA18
sgRNA31
sgRNA33
sgRNA35
sgRNA39
sgRNA45
sgRNA51
sgRNA52
sgRNA56
sgRNA58
sgRNA59
sgRNA66
sgRNA68
sgRNA73
sgRNAct
sgRNAct

Sample 7

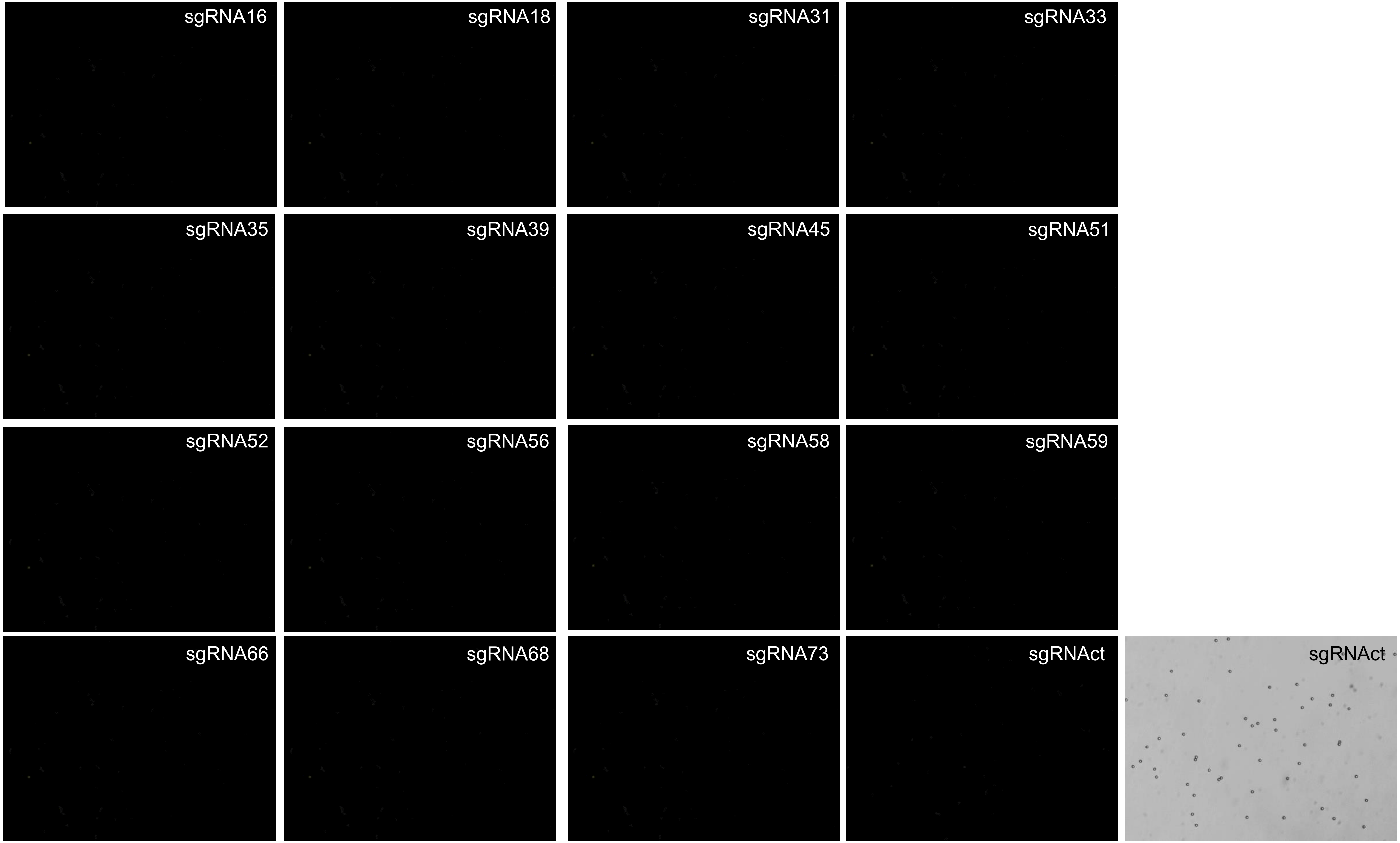
sgRNA16
sgRNA18
sgRNA31
sgRNA33
sgRNA35
sgRNA39
sgRNA45
sgRNA51
sgRNA52
sgRNA56
sgRNA58
sgRNA59
sgRNA66
sgRNA68
sgRNA73
sgRNAct
sgRNAct

Sample 8

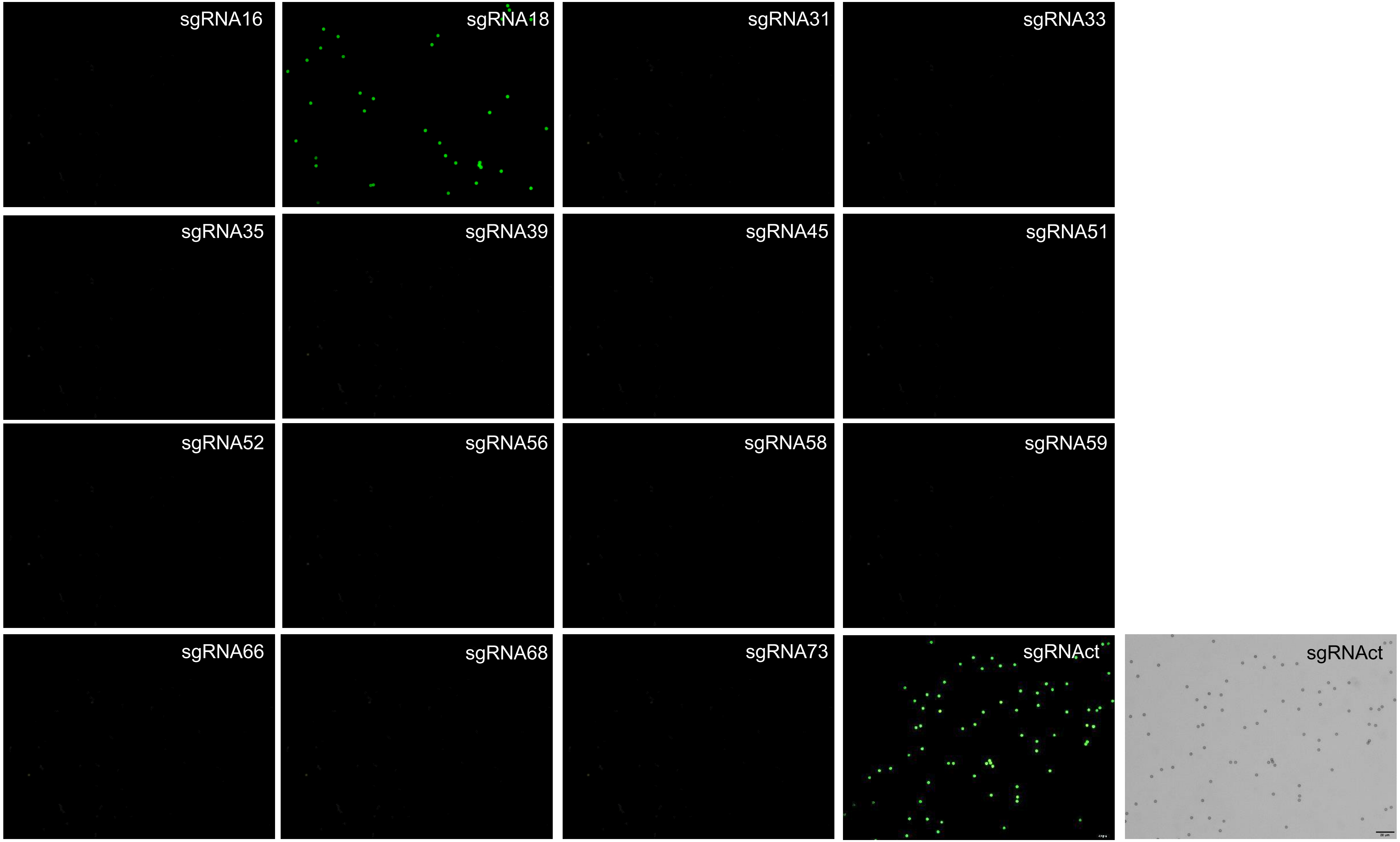
sgRNA16
sgRNA18
sgRNA31
sgRNA33
sgRNA35
sgRNA39
sgRNA45
sgRNA51
sgRNA52
sgRNA56
sgRNA58
sgRNA59
sgRNA66
sgRNA68
sgRNA73
sgRNAct
sgRNAct

Sample 11

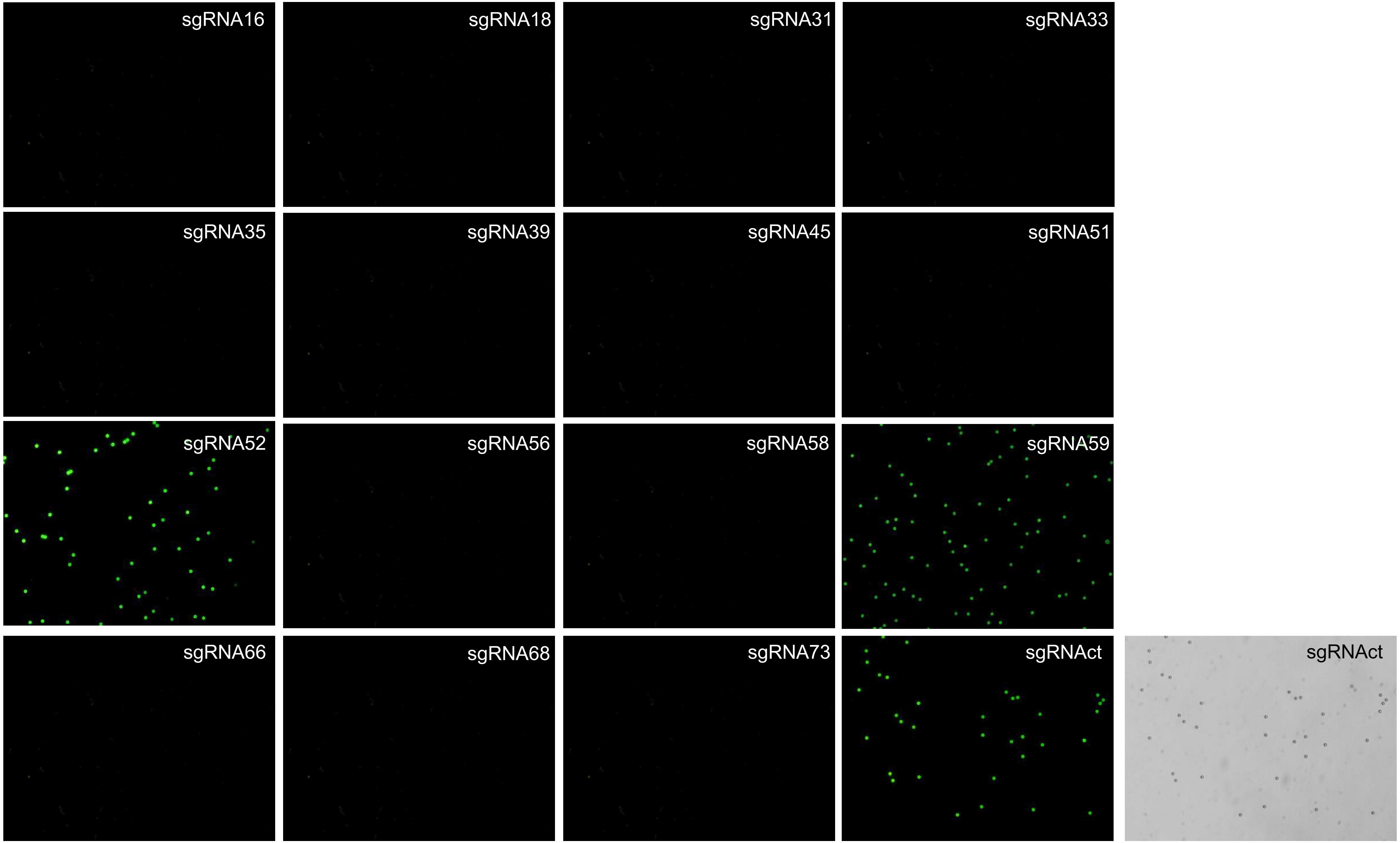
sgRNA16
sgRNA18
sgRNA31
sgRNA33
sgRNA35
sgRNA39
sgRNA45
sgRNA51
sgRNA52
sgRNA56
sgRNA58
sgRNA59
sgRNA66
sgRNA68
sgRNA73
sgRNAct
sgRNAct

Sample 16

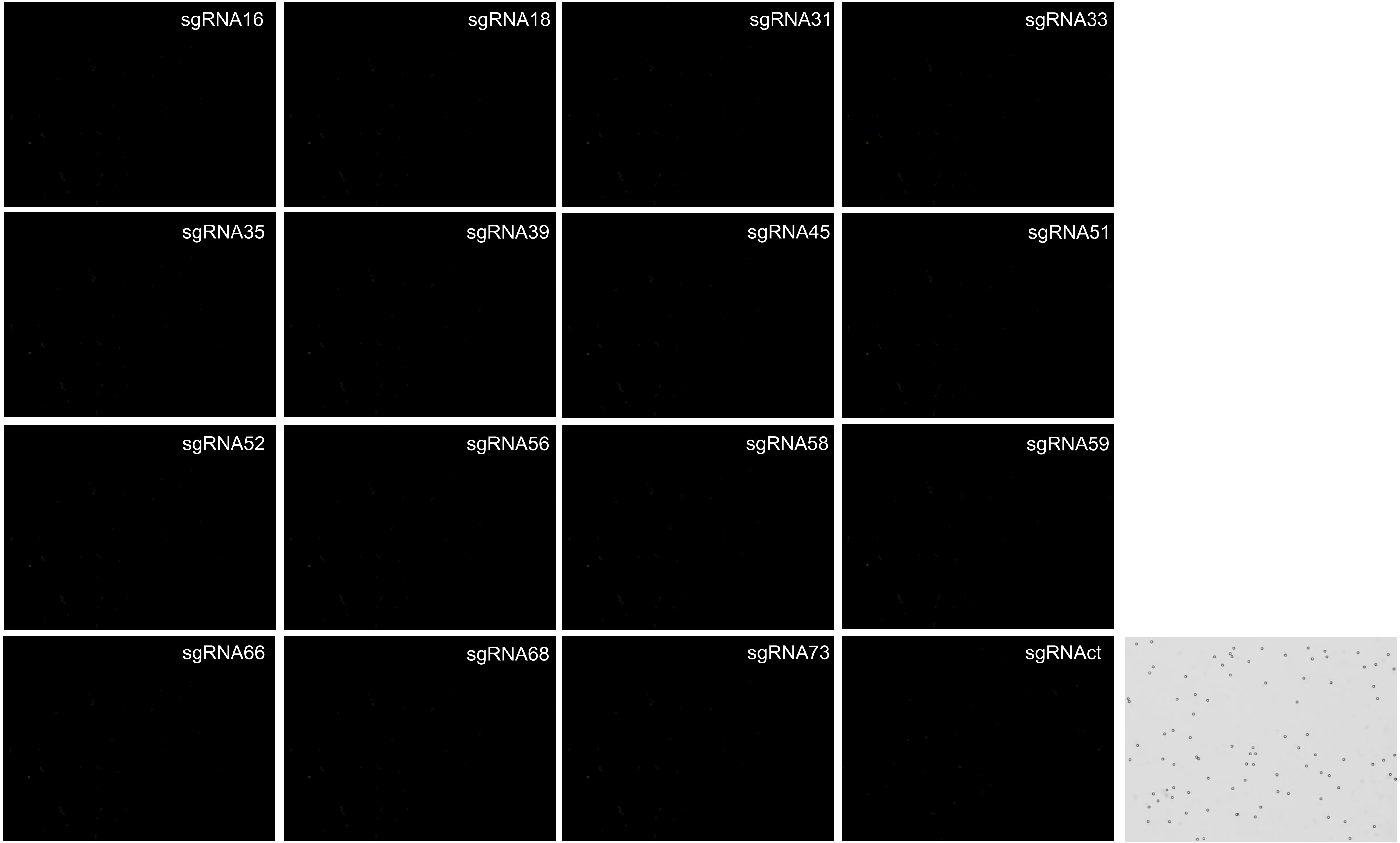
sgRNA16
sgRNA18
sgRNA31
sgRNA33
sgRNA35
sgRNA39
sgRNA45
sgRNA51
sgRNA52
sgRNA56
sgRNA58
sgRNA59
sgRNA66
sgRNA68
sgRNA73
sgRNAct

Sample 19

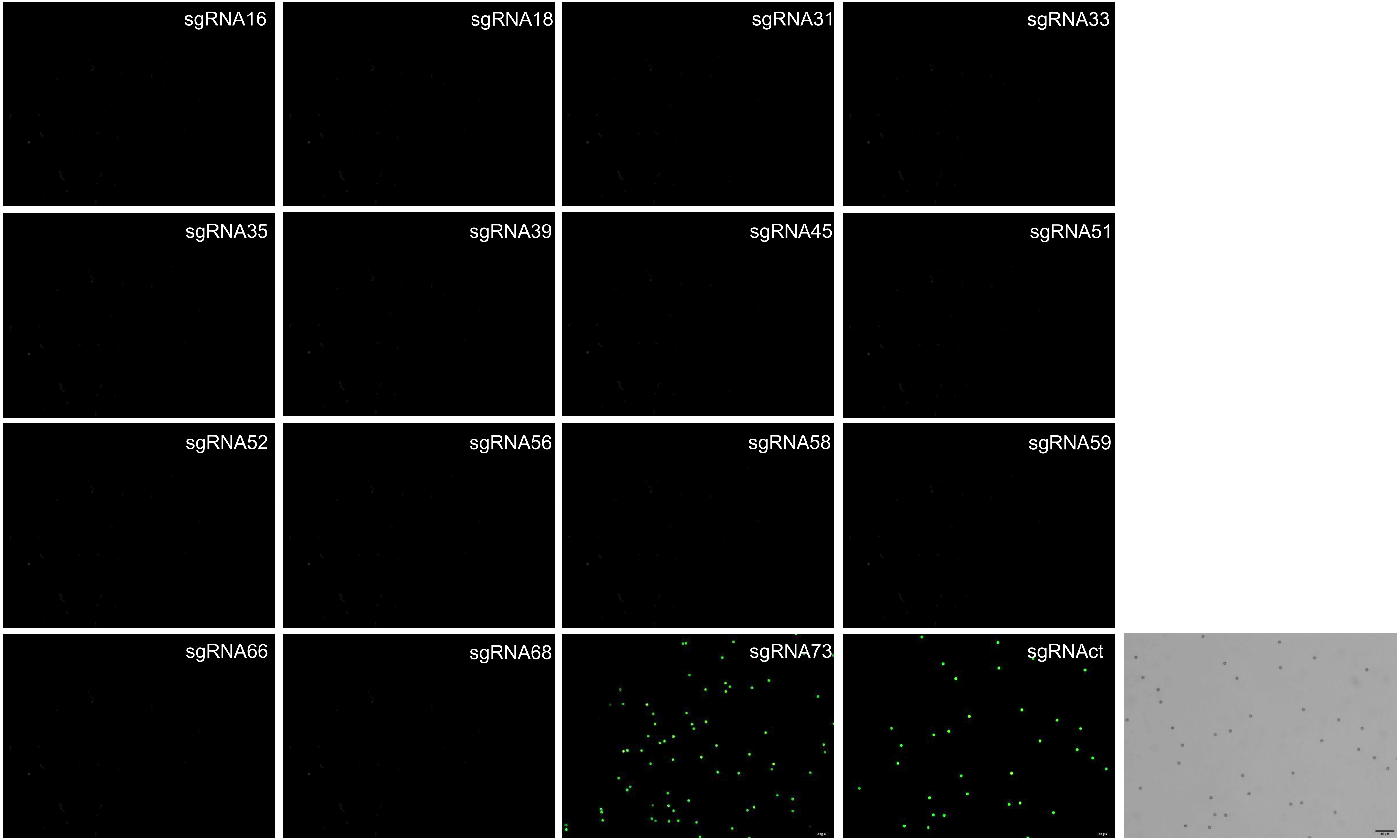
sgRNA16
sgRNA18
sgRNA31
sgRNA33
sgRNA35
sgRNA39
sgRNA45
sgRNA51
sgRNA52
sgRNA56
sgRNA58
sgRNA59
sgRNA66
sgRNA68
sgRNA73
sgRNAct

Sample 26

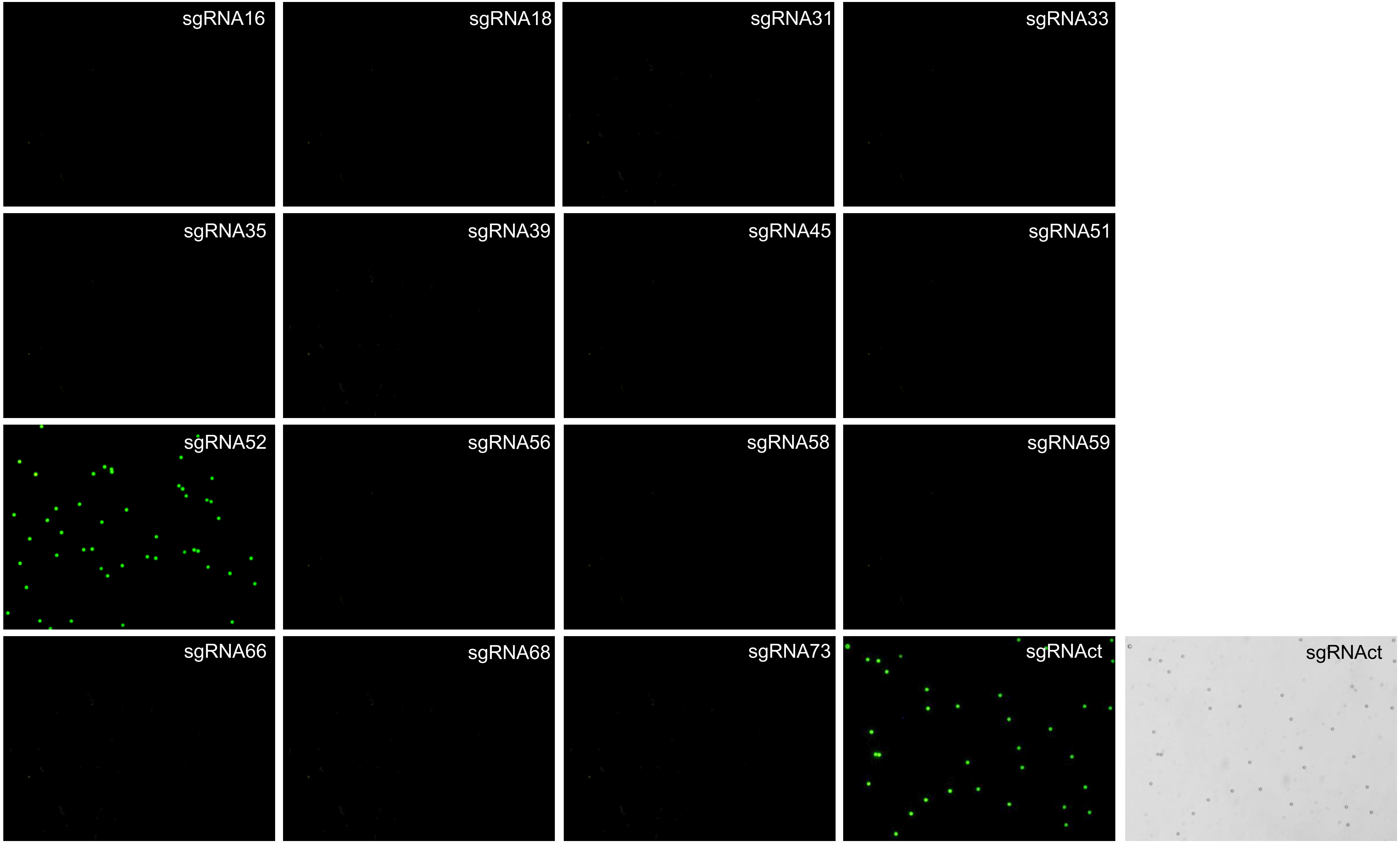
sgRNA16
sgRNA18
sgRNA31
sgRNA33
sgRNA35
sgRNA39
sgRNA45
sgRNA51
sgRNA52
sgRNA56
sgRNA58
sgRNA59
sgRNA66
sgRNA68
sgRNA73
sgRNAct
sgRNAct

Sample 35

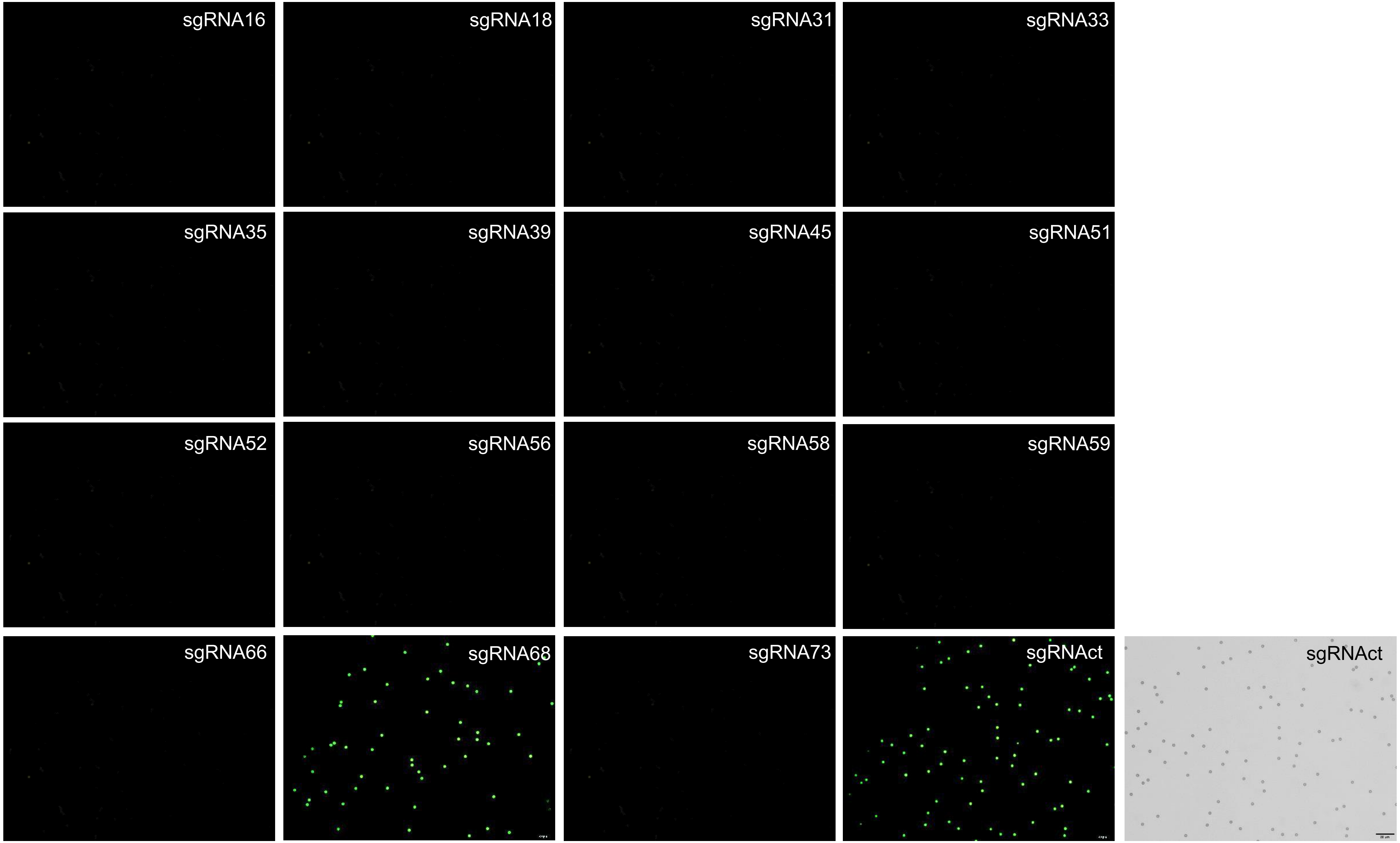
sgRNA16
sgRNA18
sgRNA31
sgRNA33
sgRNA35
sgRNA39
sgRNA45
sgRNA51
sgRNA52
sgRNA56
sgRNA58
sgRNA59
sgRNA66
sgRNA68
sgRNA73
sgRNAct
sgRNAct

Sample 36

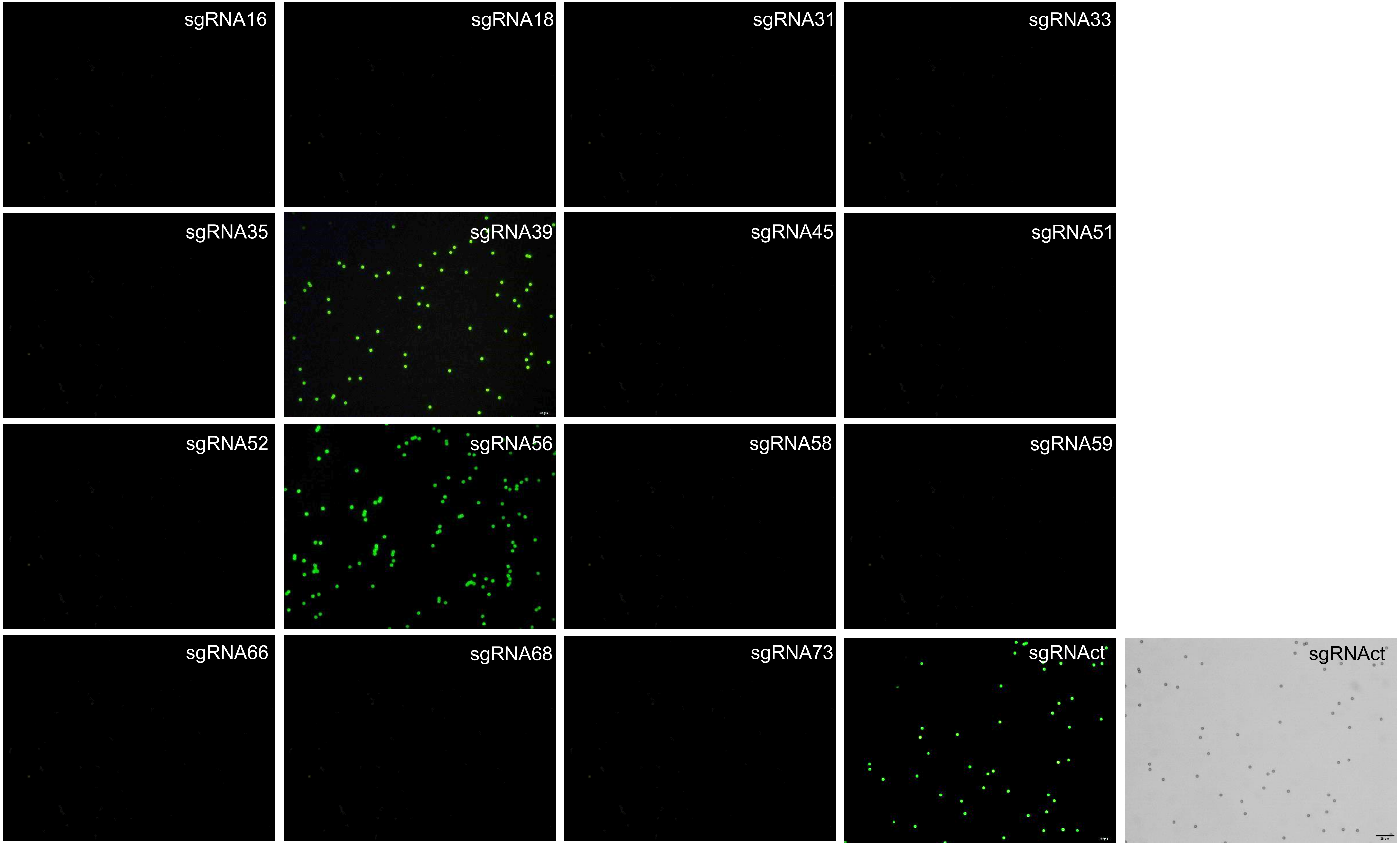
sgRNA16
sgRNA18
sgRNA31
sgRNA33
sgRNA35
sgRNA39
sgRNA45
sgRNA51
sgRNA52
sgRNA56
sgRNA58
sgRNA59
sgRNA66
sgRNA68
sgRNA73
sgRNAct
sgRNAct

Sample 37

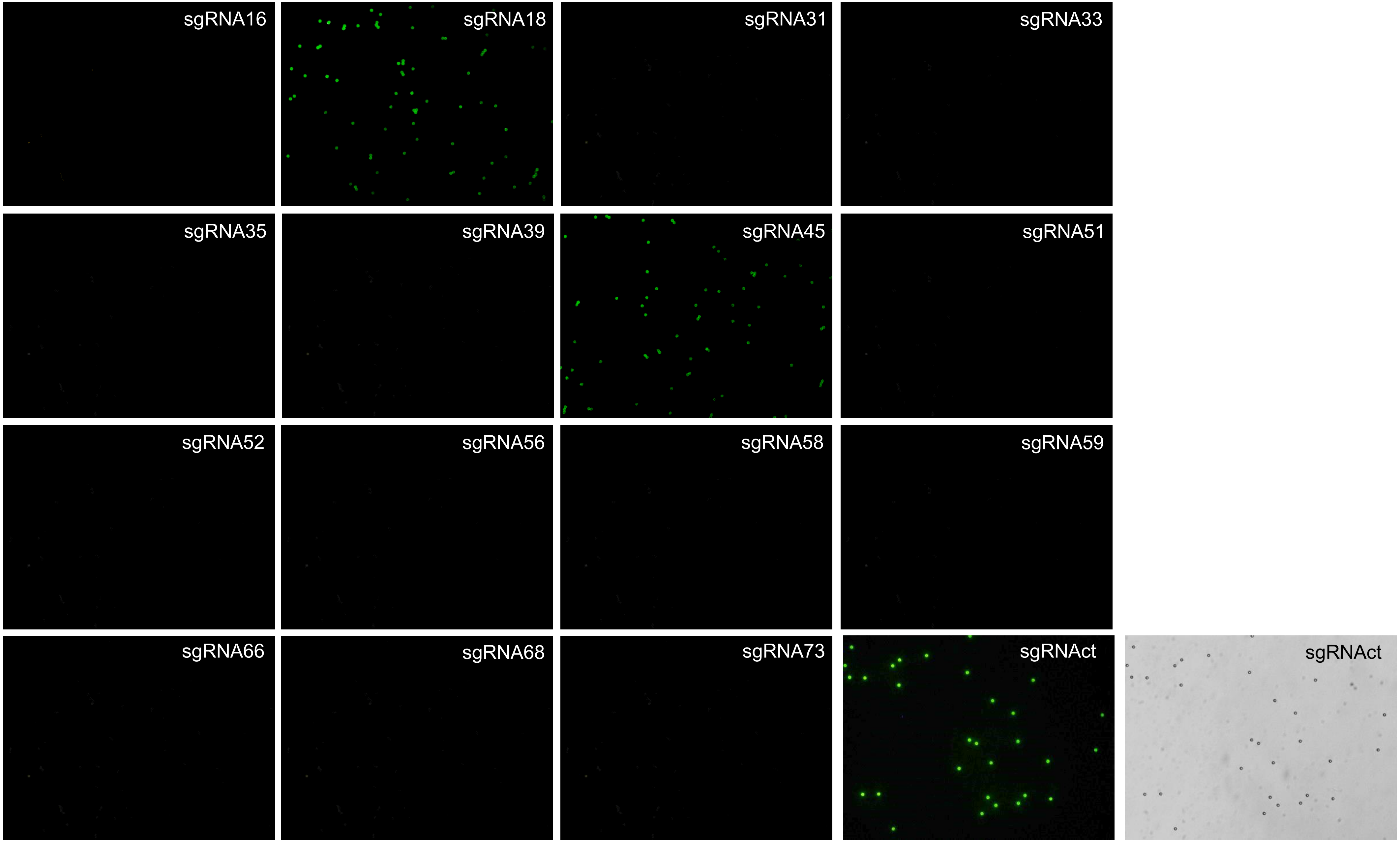
sgRNA16
sgRNA18
sgRNA31
sgRNA33
sgRNA35
sgRNA39
sgRNA45
sgRNA51
sgRNA52
sgRNA56
sgRNA58
sgRNA59
sgRNA66
sgRNA68
sgRNA73
sgRNAct
sgRNAct

Sample 39

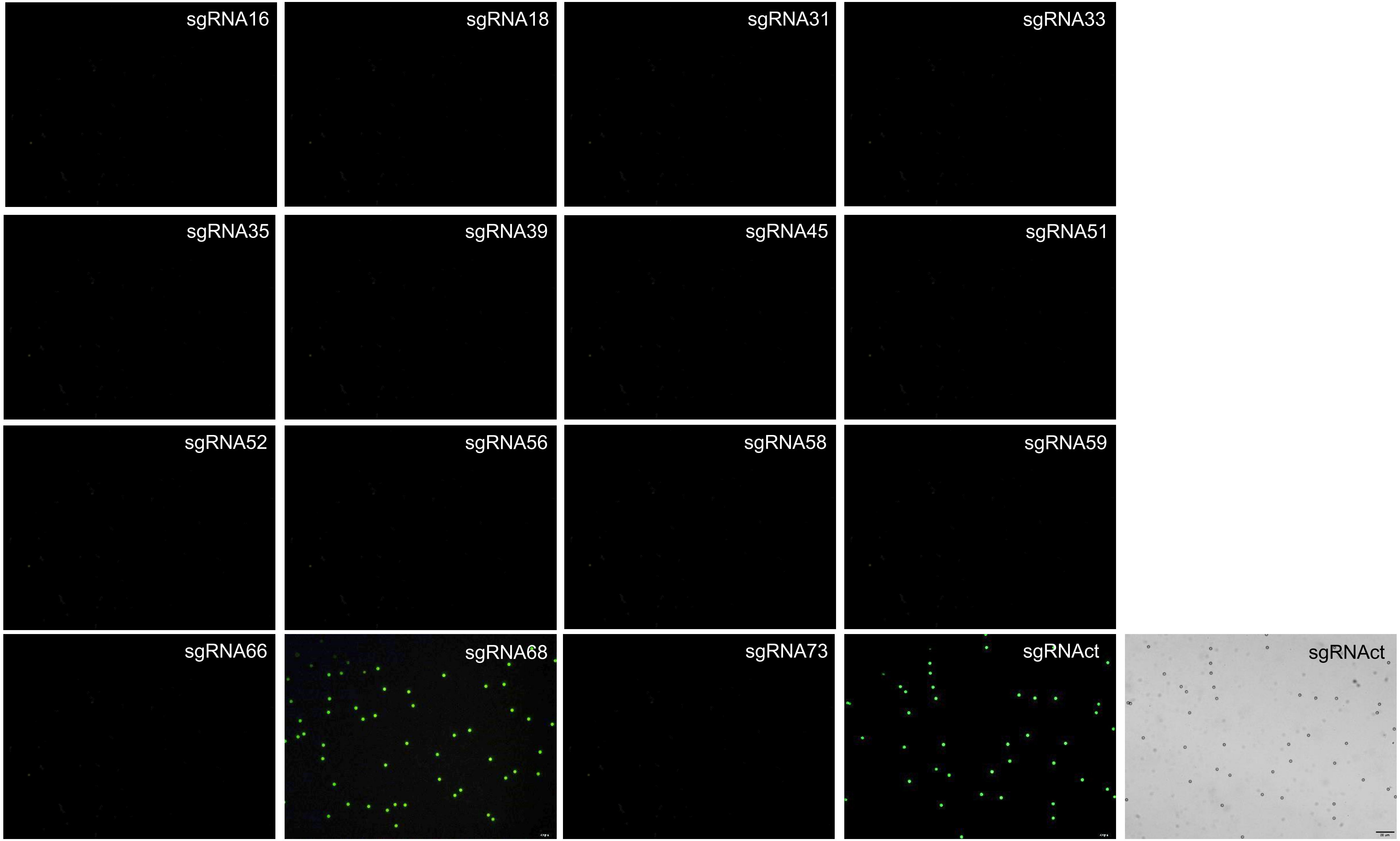
sgRNA16
sgRNA18
sgRNA31
sgRNA33
sgRNA35
sgRNA39
sgRNA45
sgRNA51
sgRNA52
sgRNA56
sgRNA58
sgRNA59
sgRNA66
sgRNA68
sgRNA73
sgRNAct
sgRNAct

Sample 42

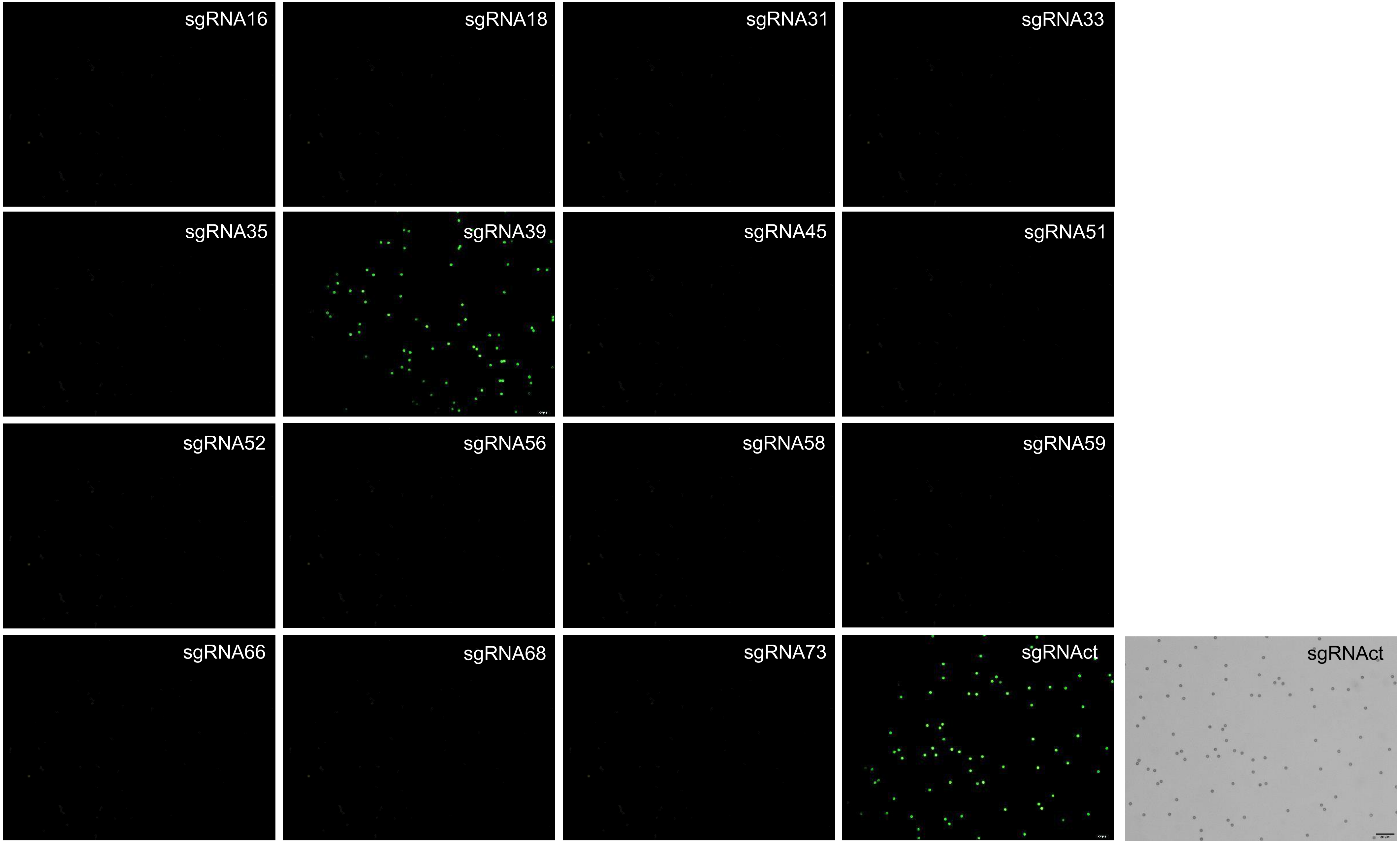

sgRNA16
sgRNA18
sgRNA31
sgRNA33
sgRNA35
sgRNA39
sgRNA45
sgRNA51
sgRNA52
sgRNA56
sgRNA58
sgRNA59
sgRNA66
sgRNA68
sgRNA73
sgRNAct
sgRNAct

Sample 44

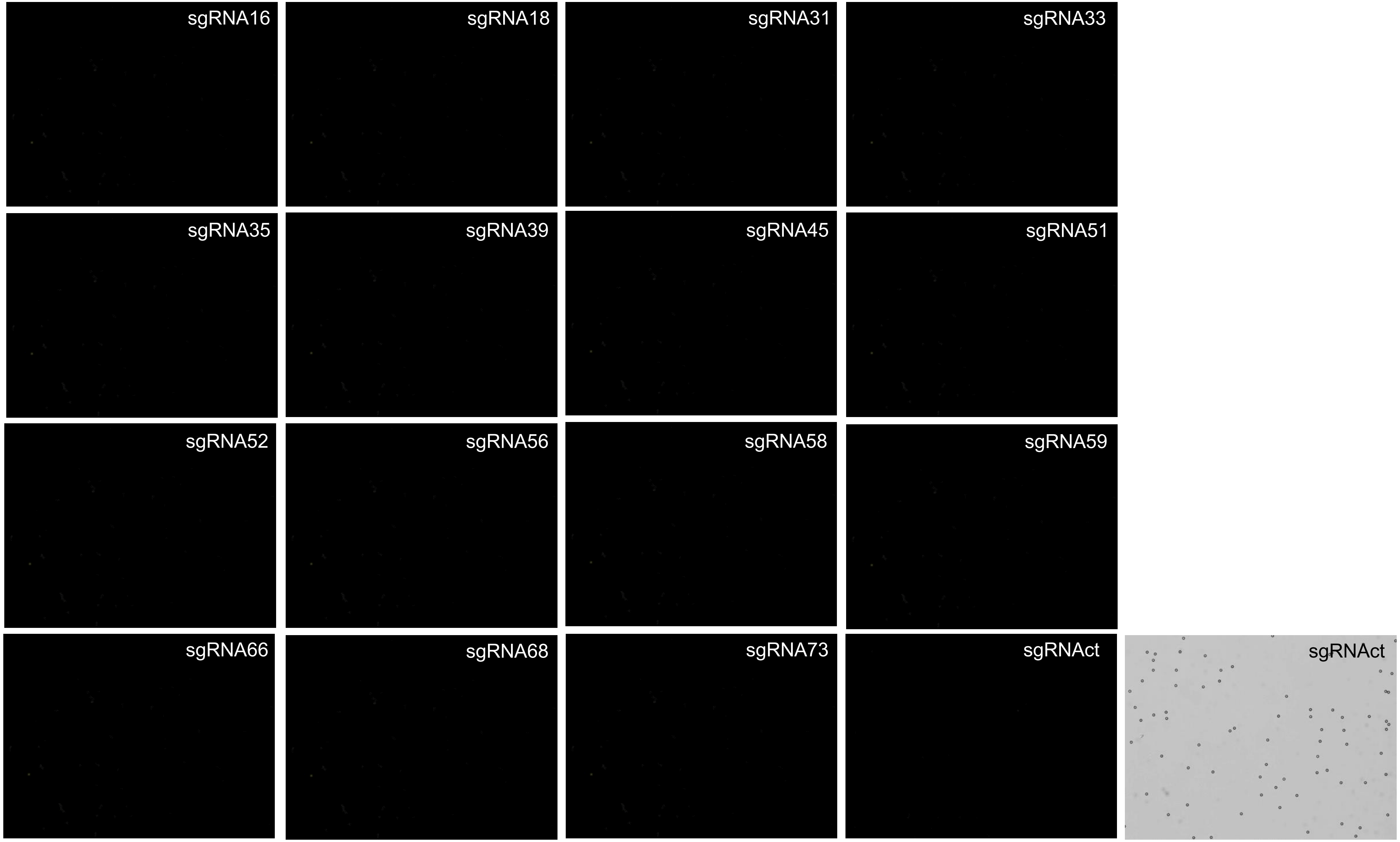
sgRNA16
sgRNA18
sgRNA31
sgRNA33
sgRNA35
sgRNA39
sgRNA45
sgRNA51
sgRNA52
sgRNA56
sgRNA58
sgRNA59
sgRNA66
sgRNA68
sgRNA73
sgRNAct
sgRNAct

Sample 50

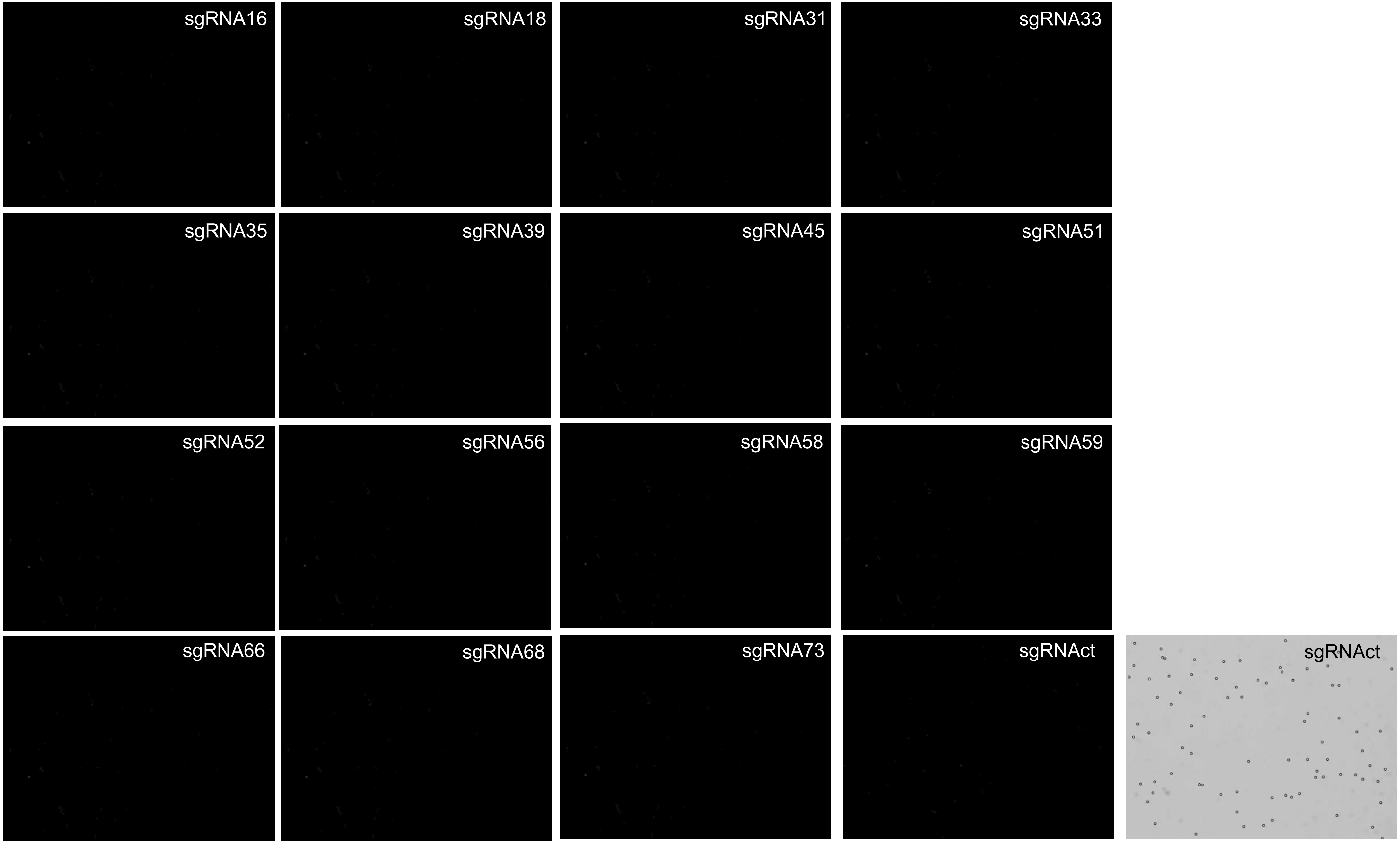
sgRNA16
sgRNA18
sgRNA31
sgRNA33
sgRNA35
sgRNA39
sgRNA45
sgRNA51
sgRNA52
sgRNA56
sgRNA58
sgRNA59
sgRNA66
sgRNA68
sgRNA73
sgRNAct
sgRNAct

Sample 57

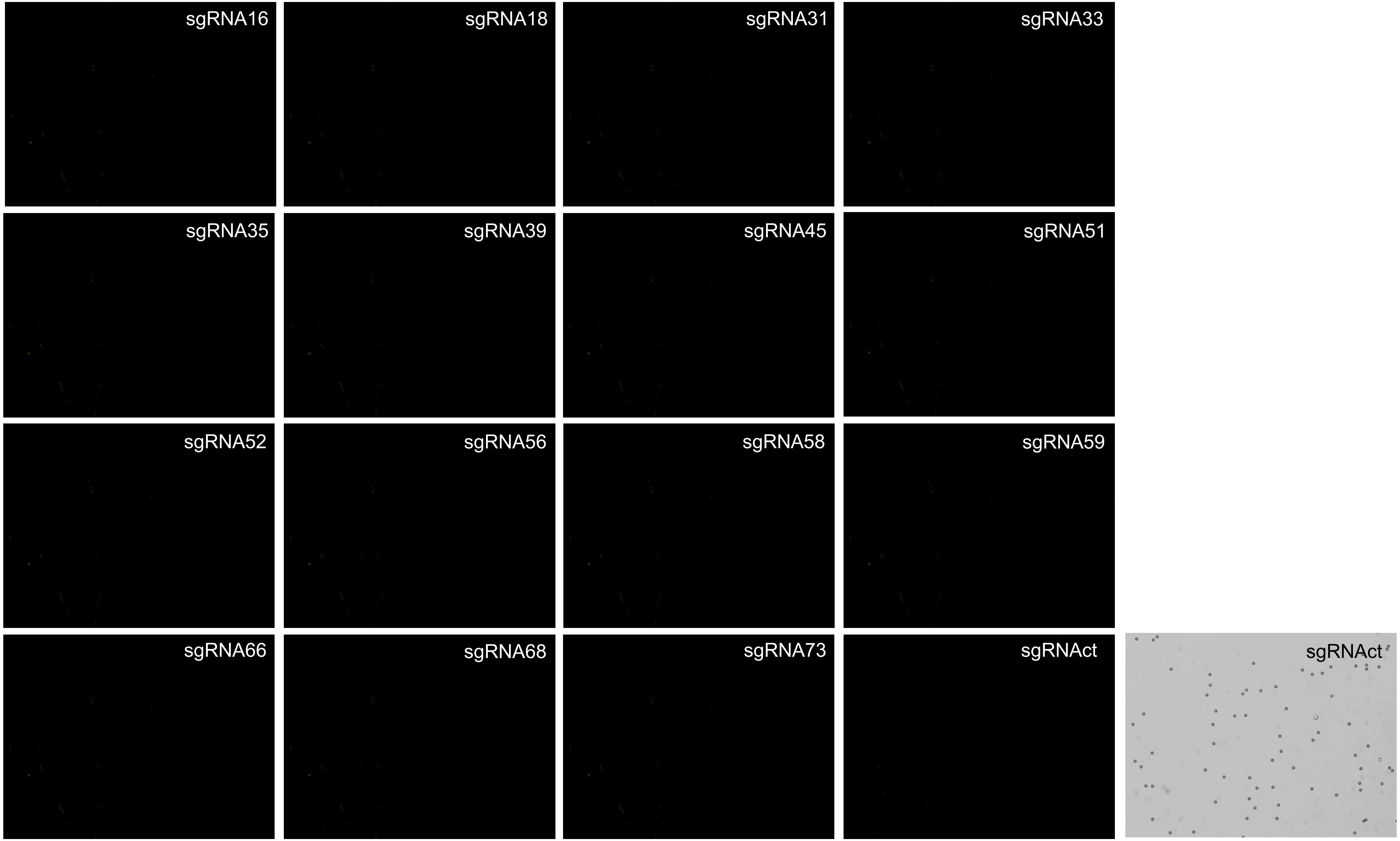
sgRNA16
sgRNA18
sgRNA31
sgRNA33
sgRNA35
sgRNA39
sgRNA45
sgRNA51
sgRNA52
sgRNA56
sgRNA58
sgRNA59
sgRNA66
sgRNA68
sgRNA73
sgRNAct
sgRNAct

Sample 61

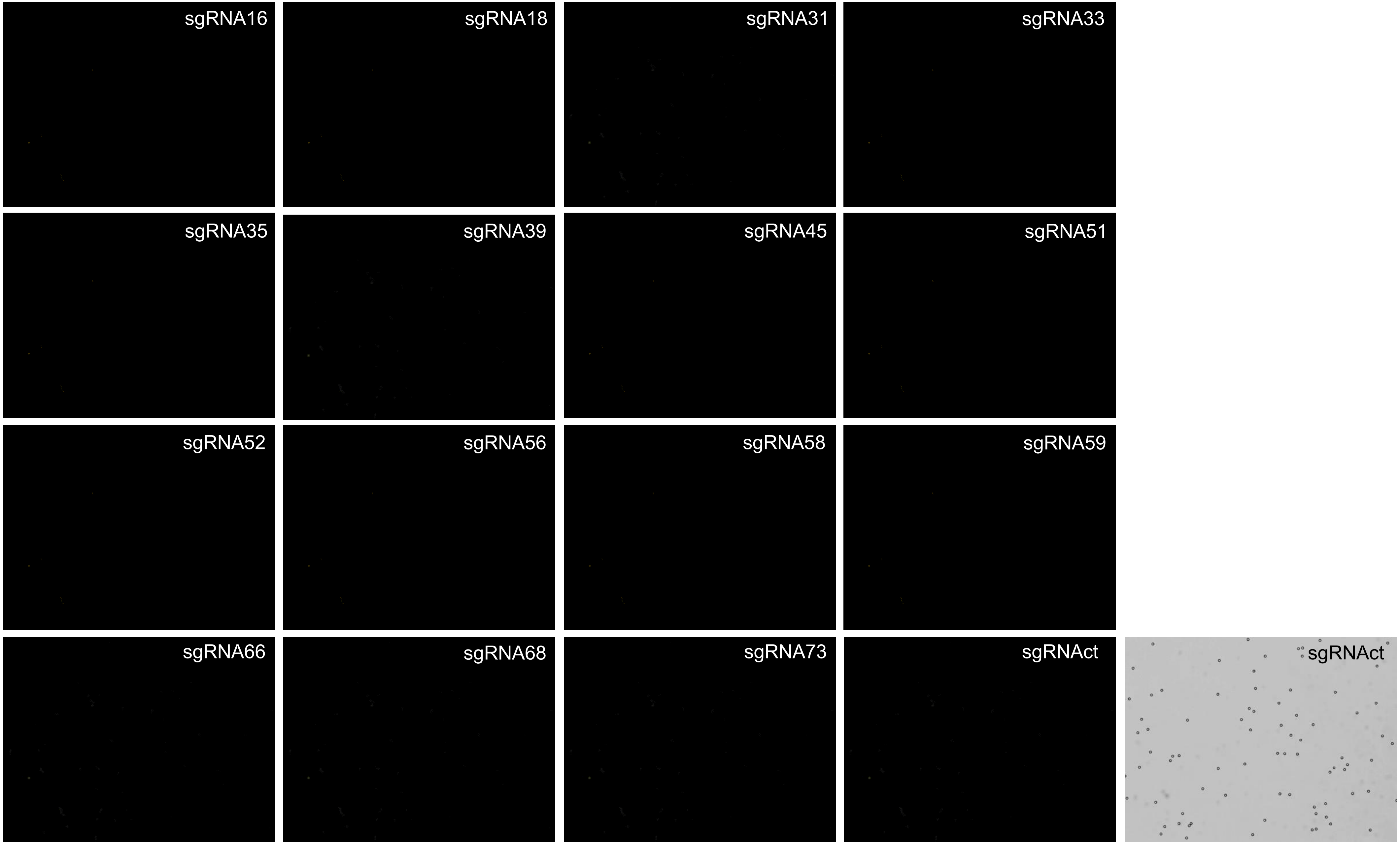
sgRNA16
sgRNA18
sgRNA31
sgRNA33
sgRNA35
sgRNA39
sgRNA45
sgRNA51
sgRNA52
sgRNA56
sgRNA58
sgRNA59
sgRNA66
sgRNA68
sgRNA73
sgRNAct
sgRNAct

Sample 66

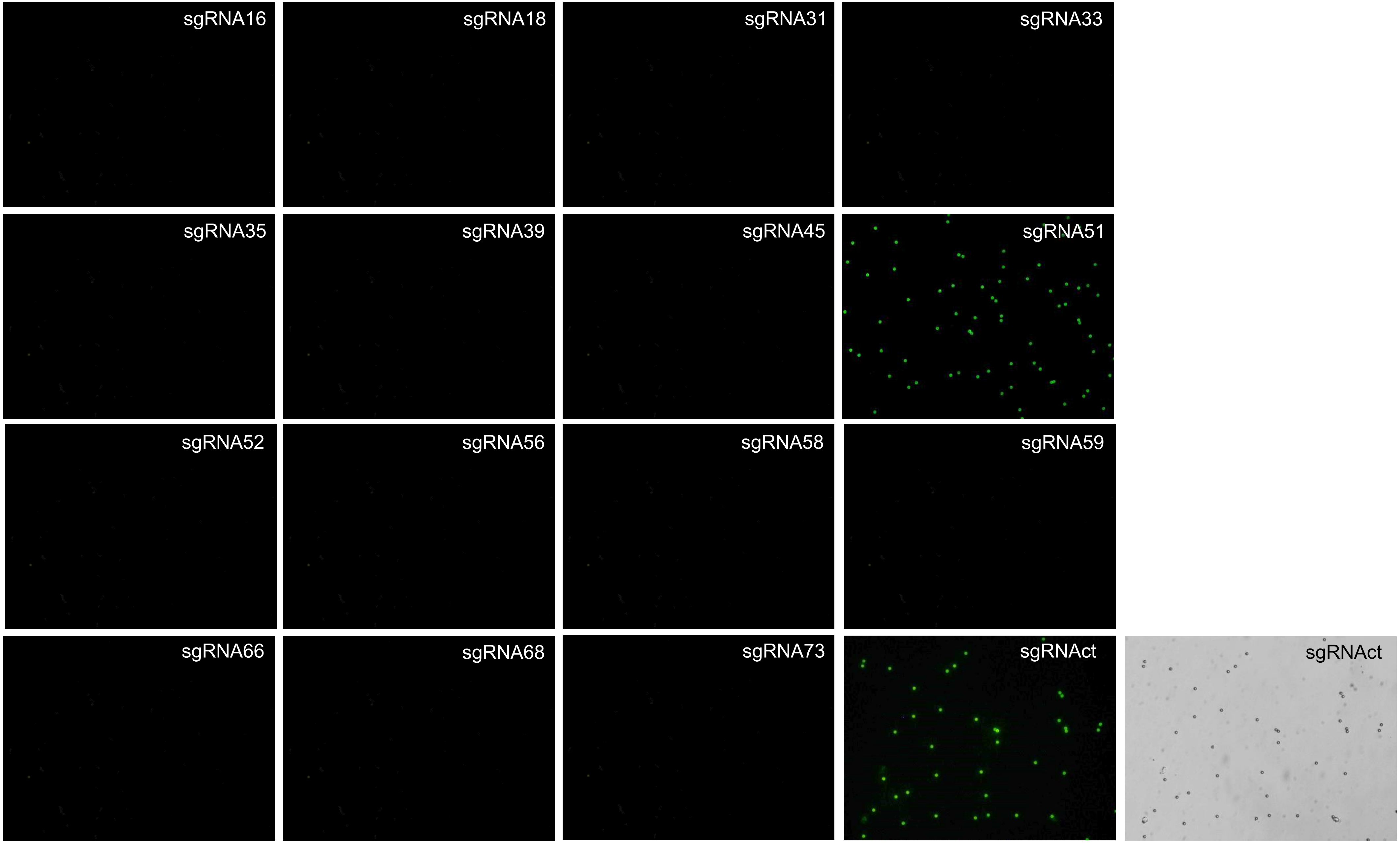
sgRNA16
sgRNA18
sgRNA31
sgRNA33
sgRNA35
sgRNA39
sgRNA45
sgRNA51
sgRNA52
sgRNA56
sgRNA58
sgRNA59
sgRNA66
sgRNA68
sgRNA73
sgRNAct
sgRNAct

Sample 67

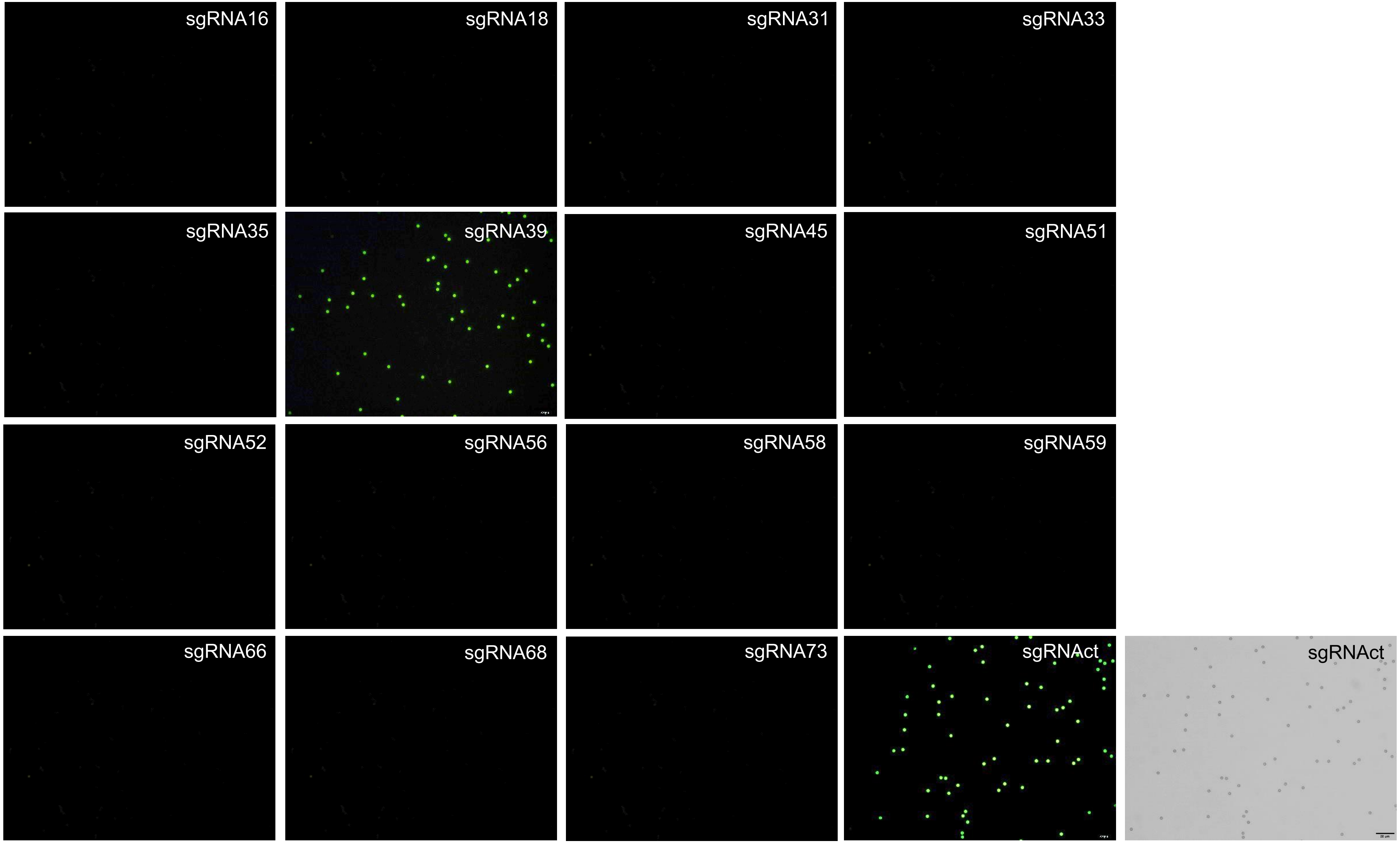
sgRNA16
sgRNA18
sgRNA31
sgRNA33
sgRNA35
sgRNA39
sgRNA45
sgRNA51
sgRNA52
sgRNA56
sgRNA58
sgRNA59
sgRNA66
sgRNA68
sgRNA73
sgRNAct
sgRNAct

Sample 68

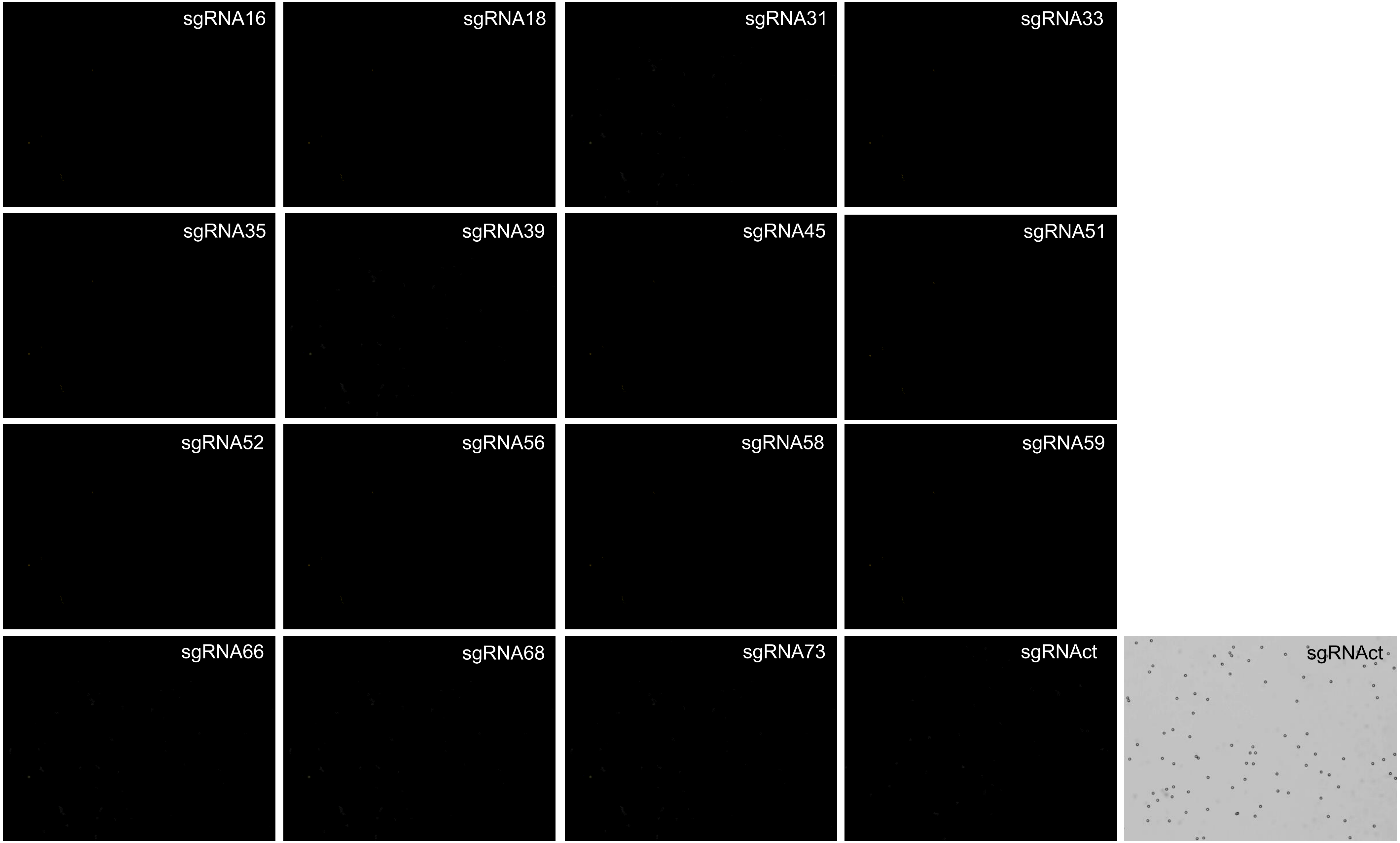
sgRNA16
sgRNA18
sgRNA31
sgRNA33
sgRNA35
sgRNA39
sgRNA45
sgRNA51
sgRNA52
sgRNA56
sgRNA58
sgRNA59
sgRNA66
sgRNA68
sgRNA73
sgRNAct
sgRNAct

Sample 70

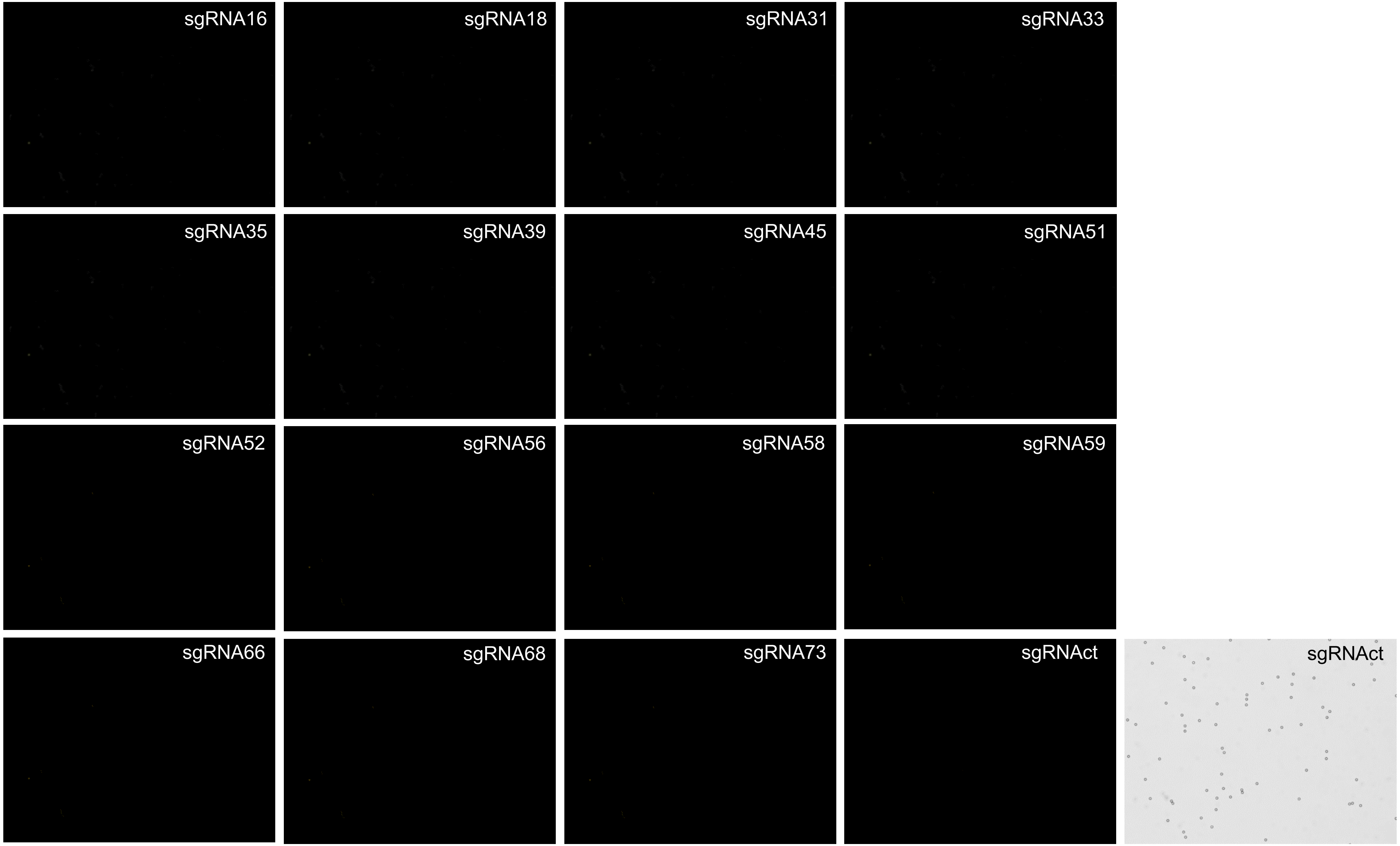
sgRNA16
sgRNA18
sgRNA31
sgRNA33
sgRNA35
sgRNA39
sgRNA45
sgRNA51
sgRNA52
sgRNA56
sgRNA58
sgRNA59
sgRNA66
sgRNA68
sgRNA73
sgRNAct
sgRNAct

Sample 71

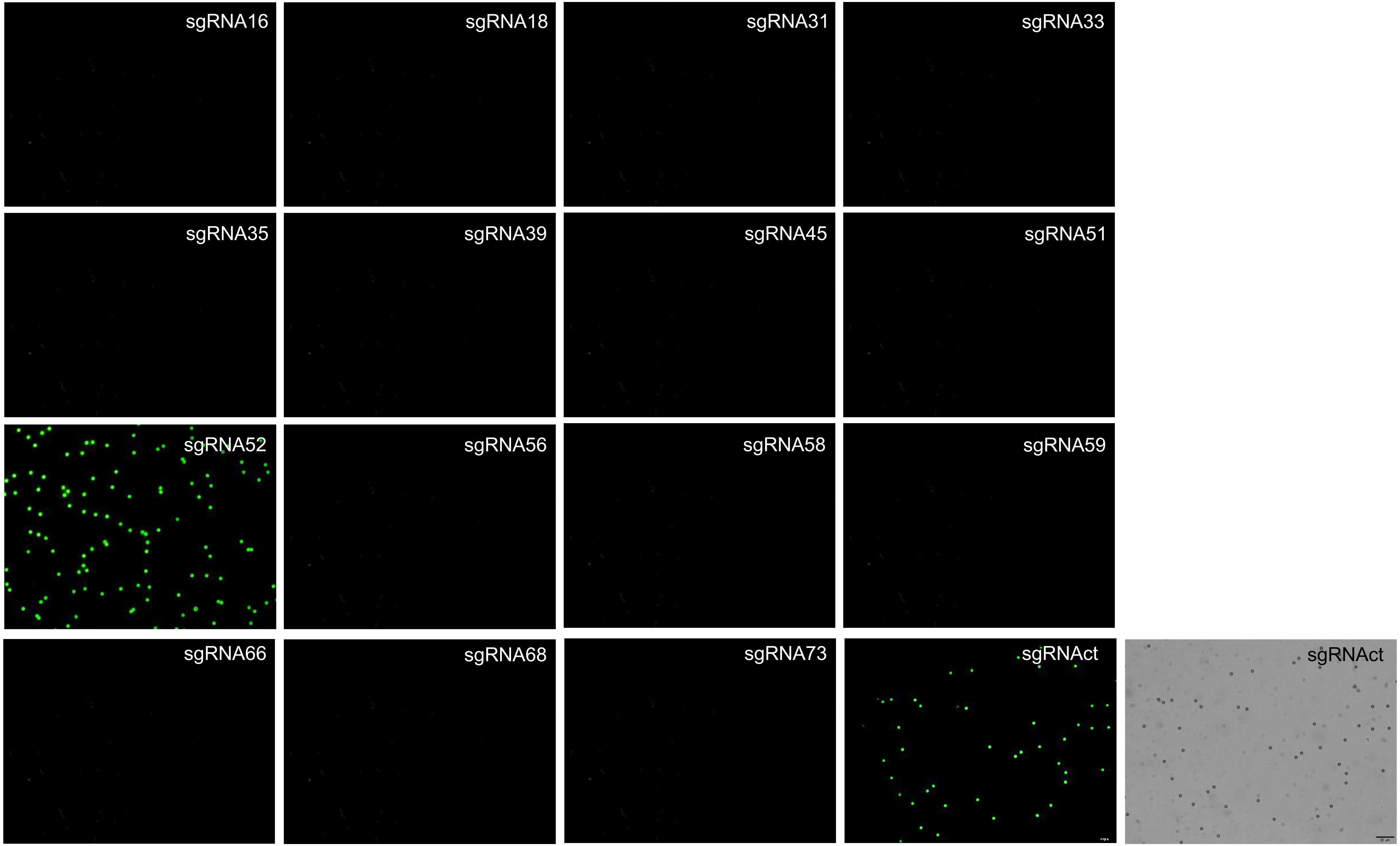
sgRNA16
sgRNA18
sgRNA31
sgRNA33
sgRNA35
sgRNA39
sgRNA45
sgRNA51
sgRNA52
sgRNA56
sgRNA58
sgRNA59
sgRNA66
sgRNA68
sgRNA73
sgRNAct
sgRNAct

Sample N17

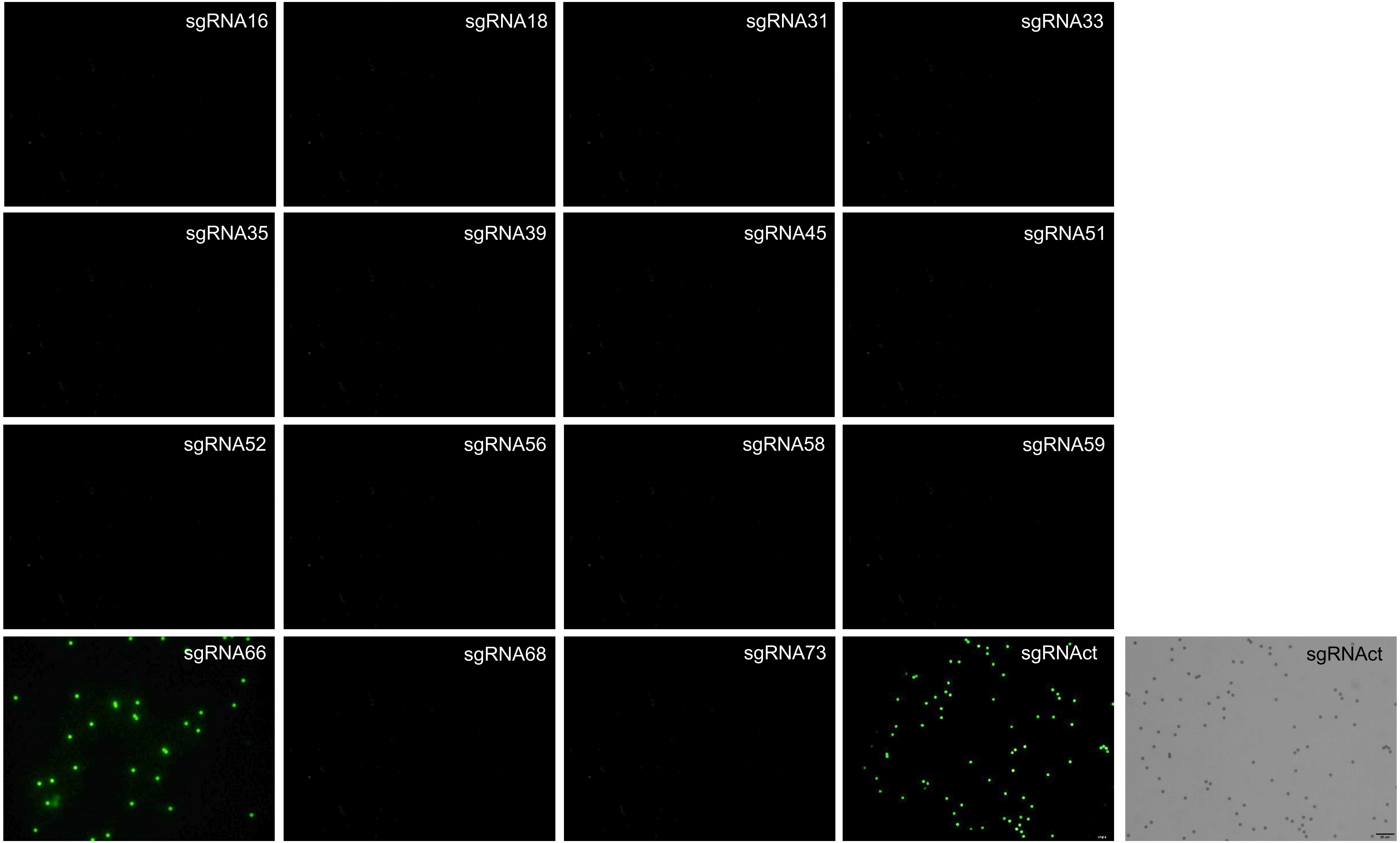
sgRNA16
sgRNA18
sgRNA31
sgRNA33
sgRNA35
sgRNA39
sgRNA45
sgRNA51
sgRNA52
sgRNA56
sgRNA58
sgRNA59
sgRNA66
sgRNA68
sgRNA73
sgRNAct
sgRNAct

Sample N26

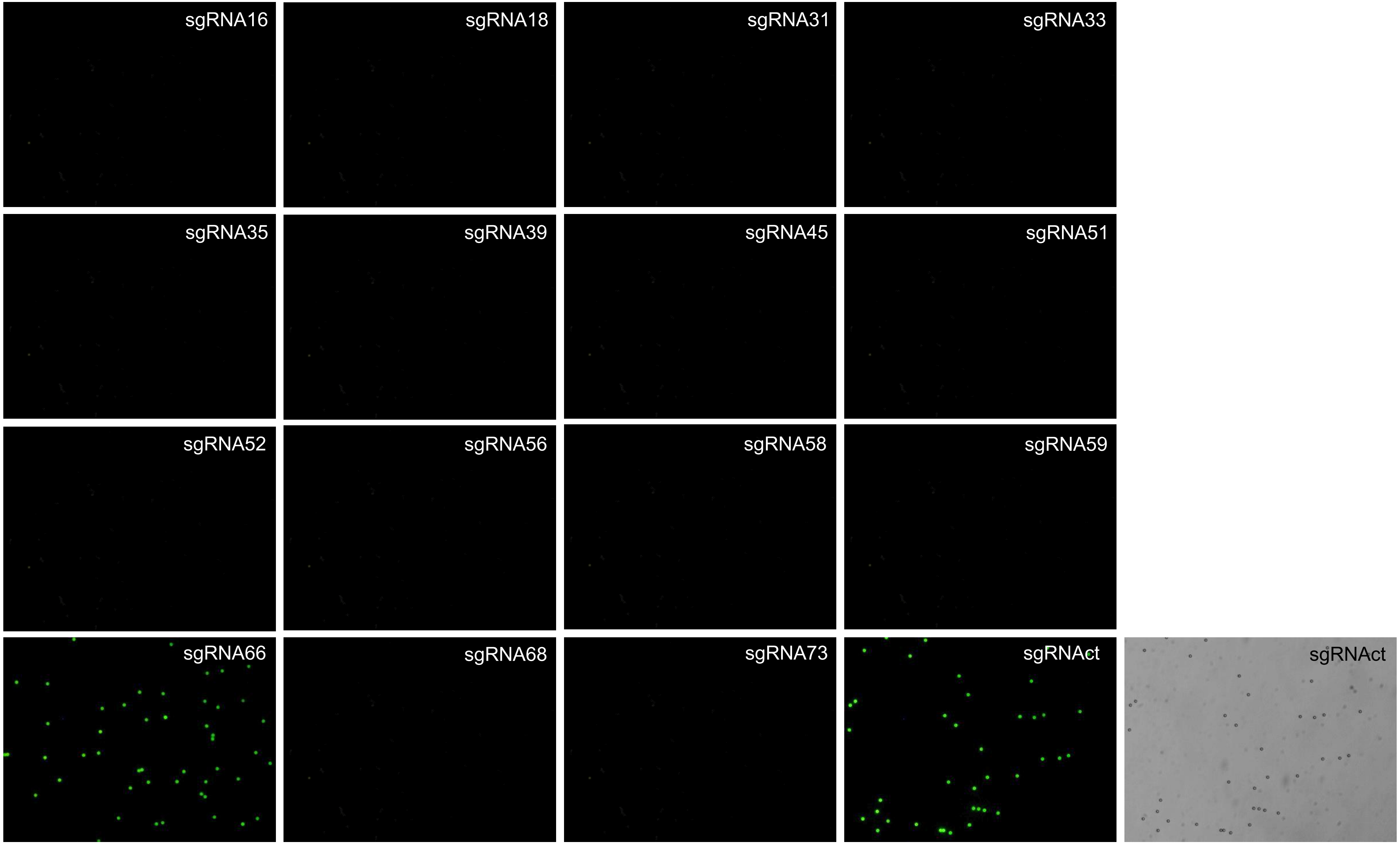
sgRNA16
sgRNA18
sgRNA31
sgRNA33
sgRNA35
sgRNA39
sgRNA45
sgRNA51
sgRNA52
sgRNA56
sgRNA58
sgRNA59
sgRNA66
sgRNA68
sgRNA73
sgRNAct
sgRNAct

Sample N34

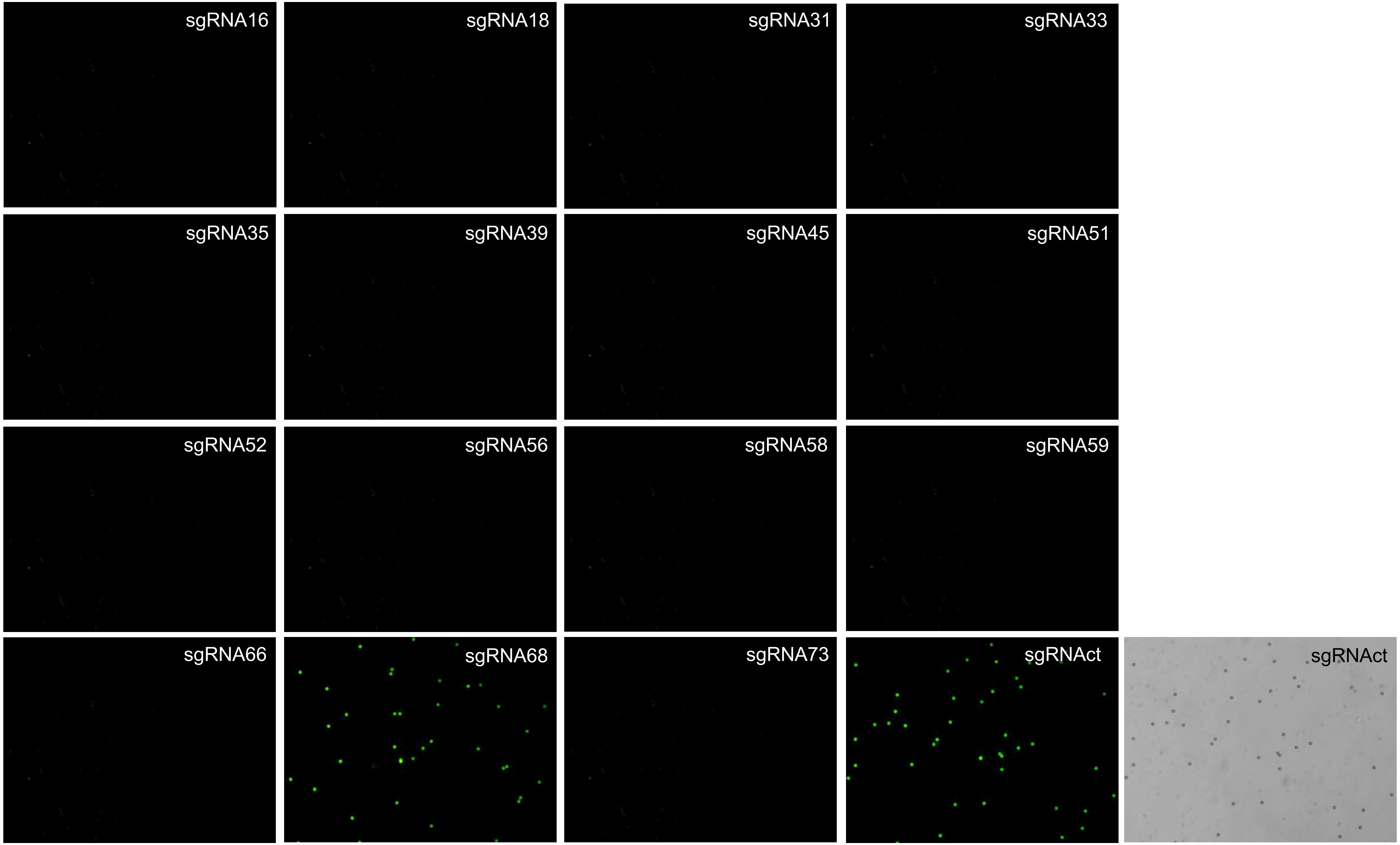
sgRNA16
sgRNA18
sgRNA31
sgRNA33
sgRNA35
sgRNA39
sgRNA45
sgRNA51
sgRNA52
sgRNA56
sgRNA58
sgRNA59
sgRNA66
sgRNA68
sgRNA73
sgRNAct
sgRNAct

Sample N60

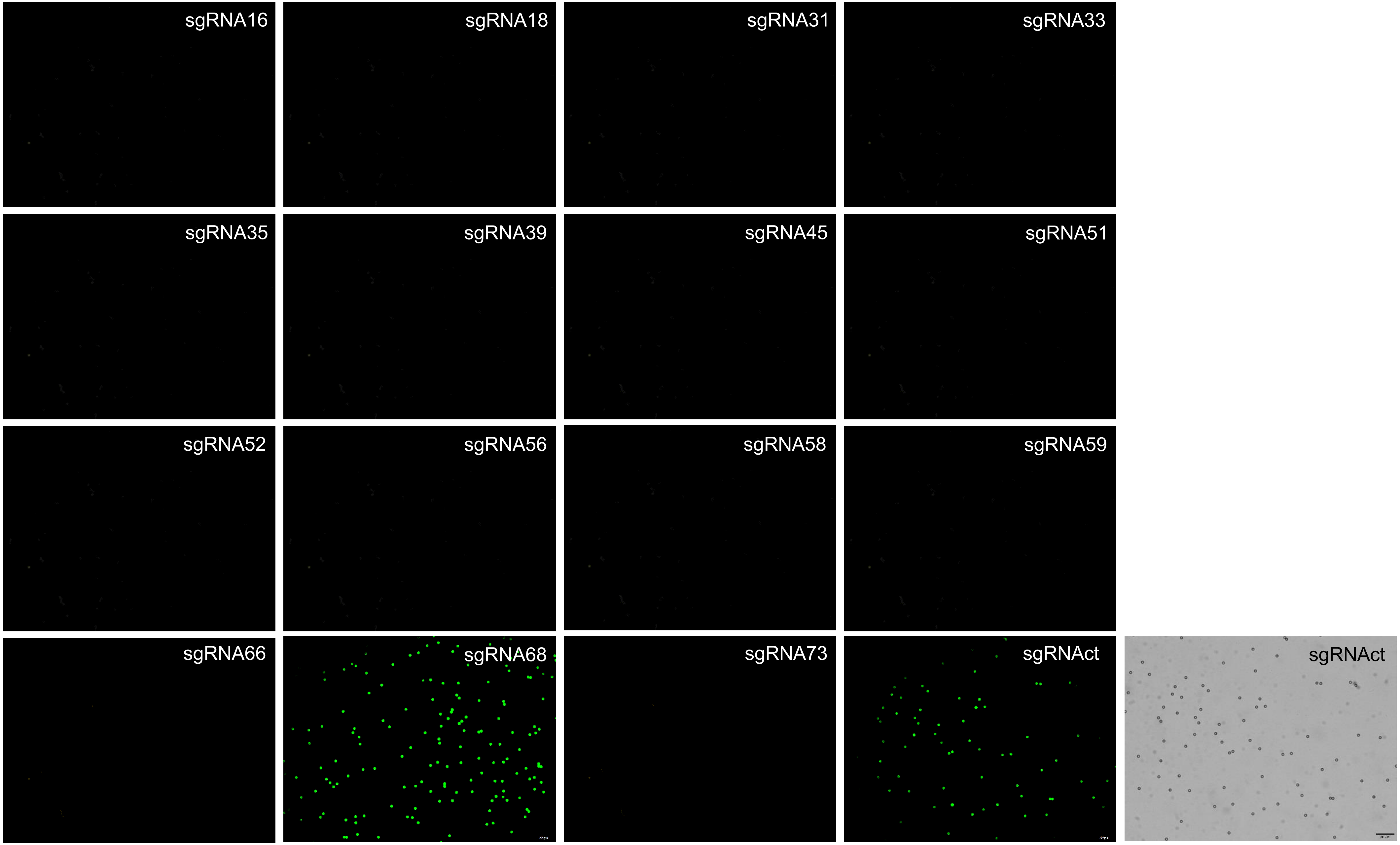
sgRNA16
sgRNA18
sgRNA31
sgRNA33
sgRNA35
sgRNA39
sgRNA45
sgRNA51
sgRNA52
sgRNA56
sgRNA58
sgRNA59
sgRNA66
sgRNA68
sgRNA73
sgRNAct
sgRNAct

Sample N79

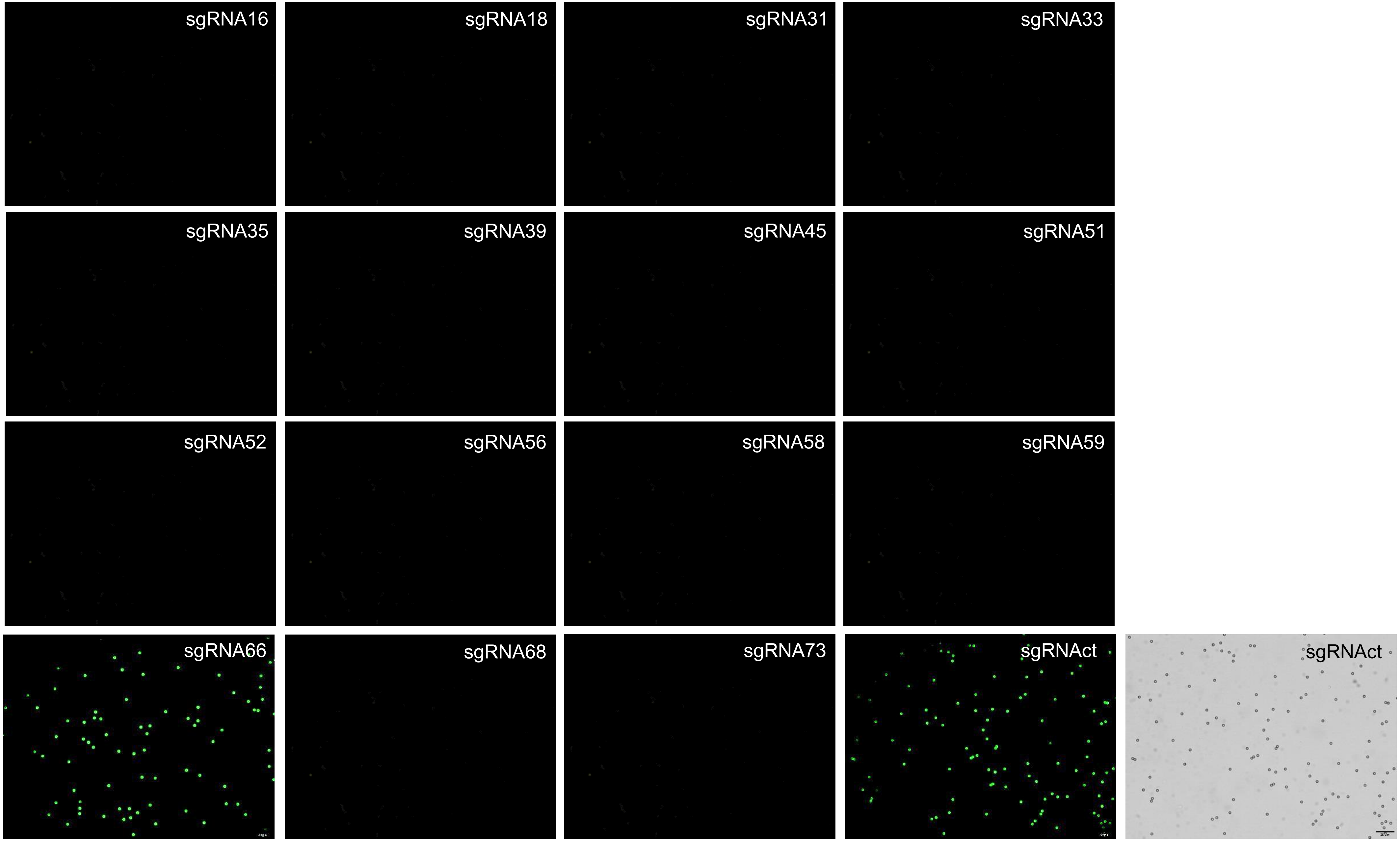
sgRNA16
sgRNA18
sgRNA31
sgRNA33
sgRNA35
sgRNA39
sgRNA45
sgRNA51
sgRNA52
sgRNA56
sgRNA58
sgRNA59
sgRNA66
sgRNA68
sgRNA73
sgRNAct
sgRNAct
